# Efficient Search of Ultra-Large Synthesis On-Demand Libraries with Chemical Language Models

**DOI:** 10.1101/2025.09.04.674350

**Authors:** Karl Heyer, David Yang, Daniel J. Diaz

## Abstract

Ultra-large ‘building block’ catalogs provide inexpensive access to billions of synthesis-on-demand molecules, but the combinatorial scale renders conventional virtual screening impractical. We present Vector Virtual Screen (VVS), a score-function-agnostic machine learning framework for efficient navigation of combinatorial libraries and rapid identification of promising molecules for experimental validation. VVS comprises four key innovations: (i) the Embedding Decomposer, which factors molecules into building blocks in latent space; (ii) ChemRank, a correlation-based loss that improves retrieval precision; (iii) BBKNN, an algorithm for nearest-neighbor search directly in building block space; and (iv) a multi-scale hill-climbing algorithm for gradient-based navigation of molecular embedding vector databases. Across diverse scoring functions, VVS consistently outperforms existing methods in retrieving high-scoring molecules while evaluating only a fraction of the library, achieving orders-of-magnitude runtime improvements. By turning ultra-large libraries into tractable search spaces, VVS enables virtual screening to keep pace with the rapid expansion of chemical space and adapt seamlessly to future advances in scoring functions.

## 1 Introduction

Ultra-large combinatorial libraries have emerged as a transformative resource in drug discovery, providing inexpensive access to billions of highly diverse synthesison-demand molecules [1–4]. These libraries are constructed from “building block” molecules that can be combinatorially assembled with a fixed set of reactions, implicitly spanning chemical spaces of 10^8^–10^12^ compounds for commercial catalogs such as Enamine and WuXi [5, 6], and up to 10^26^ for proprietary libraries [7]. Typical synthesis costs remain low (*≈* 200 USD with *≈* 80% success rates [8]), making these catalogs a cheap and practical source of chemical diversity. However their unprecedented scale renders conventional virtual screening impractical: scoring even a fraction of the space quickly exceeds computational budgets [7, 9]. Efficient strategies are therefore needed to navigate these libraries without exhaustive enumeration and evaluation.

Several classes of methods address this challenge. Similarity-based analog retrieval methods fragment query molecules and match fragments to vendor catalogs using cheminformatics rules and fingerprints [10–13], as well as synthon/fragments workflows [14, 15] and graph-edit–based matching [16]. However, these implementations remain largely proprietary and rigid. Embedding-based approaches leverage chemical language models for fast nearest-neighbor retrieval [17, 18] and scalable vector indexes [19], yet require full library enumeration, which becomes intractable at gigascale. In parallel, several ML methods for general retrosynthesis have been proposed [20–24], but they target total synthesis rather than catalog-constrained analog retrieval. Fragment-based methods reduce the search space by evaluating individual building blocks in the context of a protein pocket, then assembling molecules from high-scoring fragments [25–27]. Some machine learning variants extend this concept by “growing” ligands within a protein pocket one fragment at a time, using the docking score of the growing ligand as a reward signal [28–31], but these methods rely on the assumption that fragment scores transfer to full molecules, which is often an invalid assumption [32]. Active Learning (AL) [33] methods reduce the search compute burden by fitting a fast surrogate model to the target score function, but this approach demands repeated retraining of the surrogate function and re-scoring over enumerated spaces [34–38], which is intractable at the scale of modern libraries. While there are methods to reduce the inference burden of active learning [39, 40], these remain insufficient at modern scales.

Taken together, existing methods either (i) require full enumeration, (ii) impose heavy computational overhead, (iii) depend on complex rules-based systems, or (iv) are tied to specific scoring functions. We provide an in depth summary of these method classes in Supp. Notes 4.1.

To overcome these limitations, we introduce three innovations (Fig. 1). First, the Embedding Decomposer model learns to decompose molecules into building blocks directly in embedding space, replacing handcrafted retrosynthetic rules with end-to-end learning. Second, Building Block K-Nearest Neighbors (BBKNN) algorithm enables efficient analog retrieval within combinatorial spaces by searching directly in building block space, avoiding explicit enumeration. Third, we present Vector Virtual Screen (VVS), a gradient-based optimization framework that integrates arbitrary scoring functions with BBKNN to efficiently explore gigascale chemical spaces. Finally, we release the VVS platform, which leverages modern software and deep learning infrastructure to deliver scalable performance and modular design, establishing VVS as a flexible foundation for virtually any future virtual screening application. We demonstrate that VVS identifies high-scoring, structurally diverse compounds across diverse objectives—including antibiotic activity prediction, 3D molecular similarity, binding affinity, and docking—while achieving orders-of-magnitude runtime improvements over existing approaches. Together, these results establish VVS as a powerful and general framework for tractable navigation of ultra-large building block libraries.

**Fig. 1.**
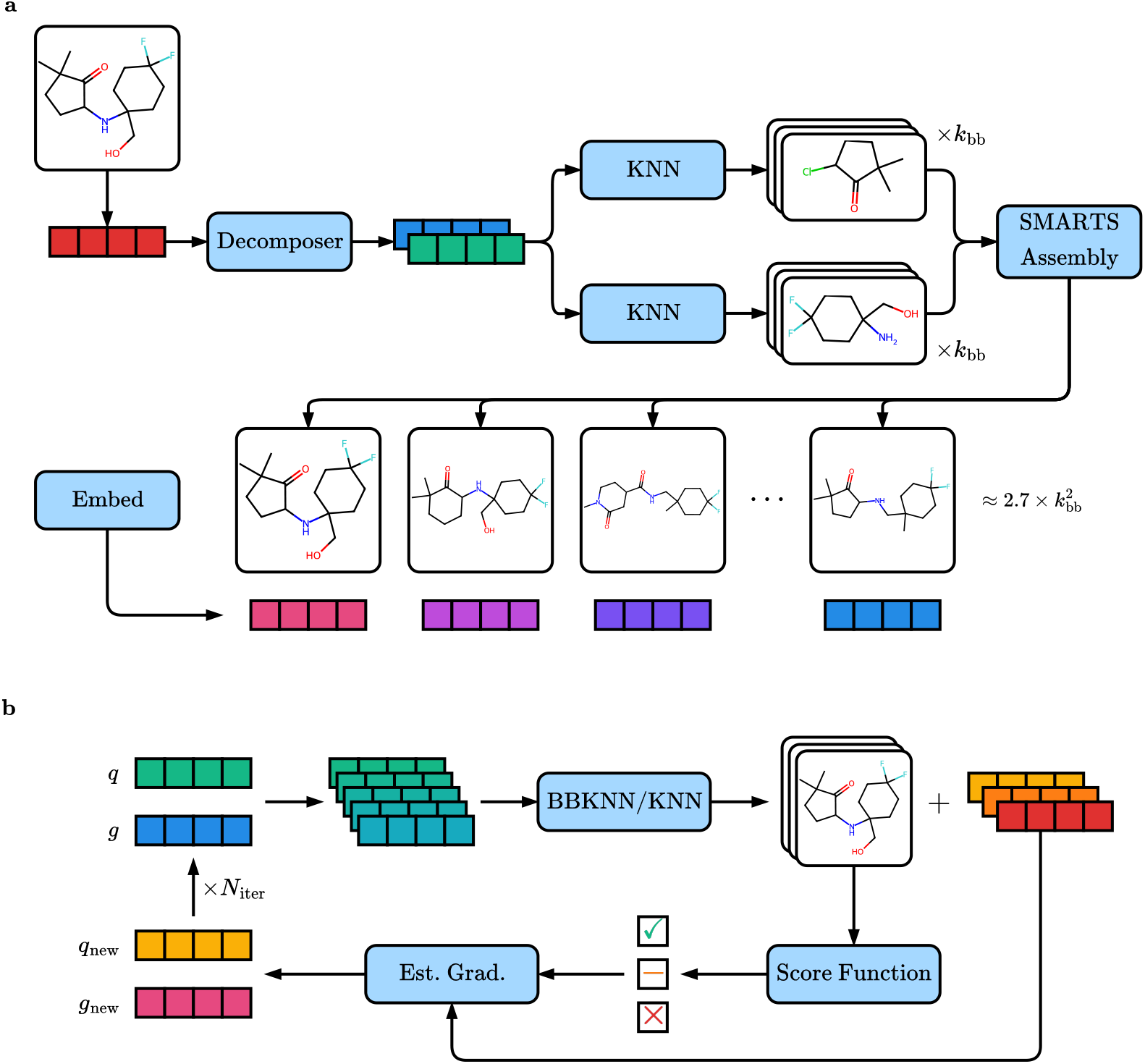
Overview of the Vector Virtual Screening framework. **a)** The Building Block K-Nearest Neighbor (BBKNN) algorithm. The Embedding Decomposer model decomposes a query embedding into two predicted building block embeddings, which are used to retrieve similar building blocks from a vector database. The retrieved building blocks are assembled into molecules via SMARTs reactions. **b)** The Vector Virtual Screening (VVS) algorithm. A query embedding and corresponding gradient are used to generate multi-scale embedding queries, which are used to retrieve nearest neighbors via BBKNN for combinatorial spaces or KNN for enumerated spaces. The retrieved results are scored with the target scoring function. The score information is used to estimate the gradient of the query and generate a new query embedding via hill-climbing.

## 2 Results

### 2.1 Machine Learning-based Building Block Decomposition

The underlying innovation that enables VVS is the Embedding Decomposer — an MLP-based model trained with self-supervised metric learning to decompose molecules into building blocks in latent space by predicting two constituent building block embeddings from a product molecule embedding, replacing complex cheminformatics heuristics with a scalable end-to-end data-driven framework. The model is trained on a dataset of 50M (building block, product molecule) pairs (Supp. Notes 4.2) embedded with a pretrained RoBERTa-ZINC model [41] using the ChemRank correlation loss, a self-supervised metric learning loss we developed for this task (Fig. 2b, Methods 4.4).

**Fig. 2.**
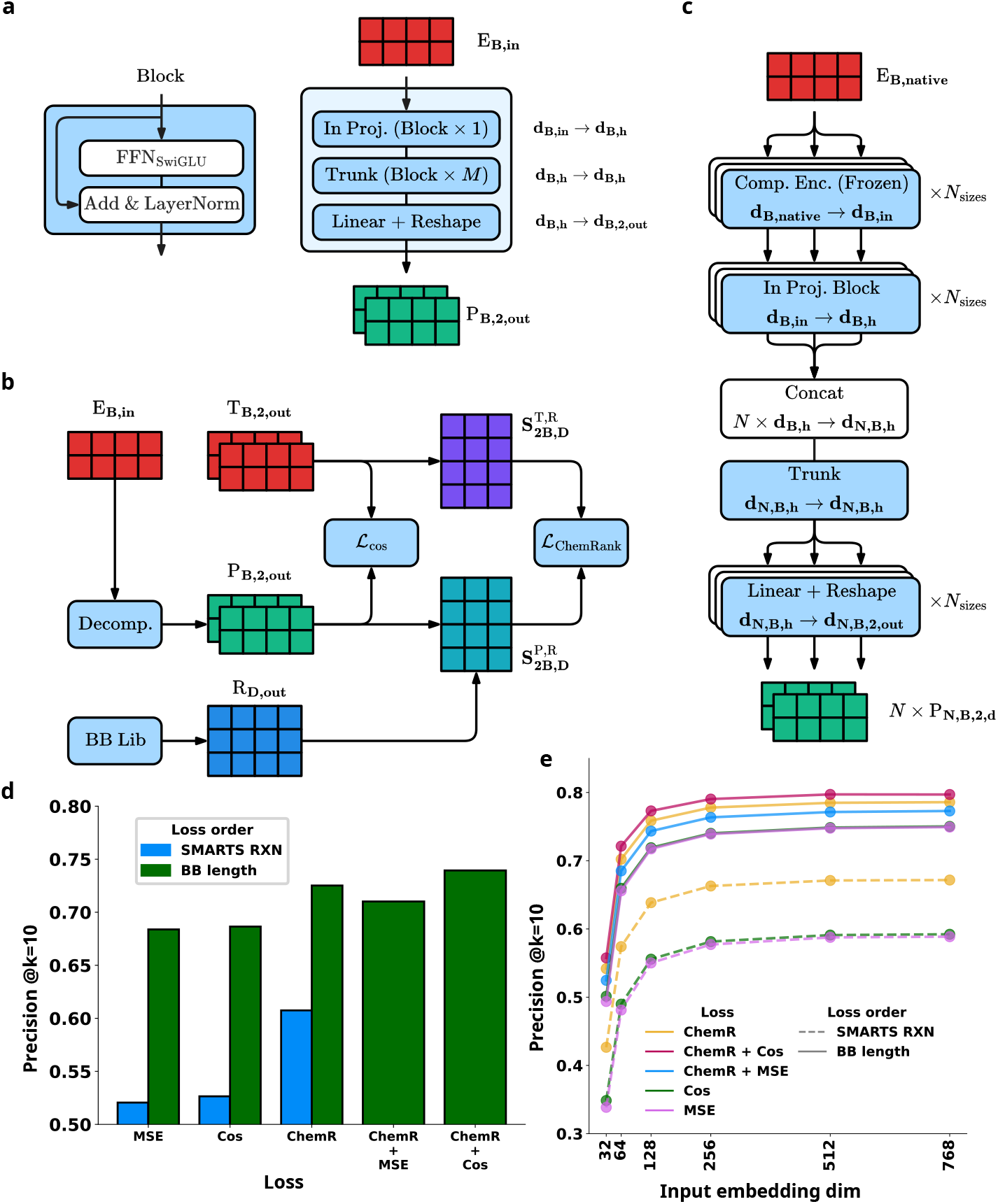
Overview of the Embedding Decomposer model. **a)** Model architecture. The input embedding *d*_in_ is projected to hidden dimension *d*_*h*_, processed by a trunk of *M* layers, and output to (2, *d*_out_). **b)** ChemRank loss. Predictions are trained with cosine similarity (*ℒ*_cos_) and an in-batch ranking loss (*ℒ* _ChemRank_) computed against randomly sampled reference embeddings. **c)** Multi-scale training. The model is trained across all (*d*_in_, *d*_out_) pairs from compressed embeddings using the frozen Embedding Compression encoders. Compressed inputs are projected to a common hidden size, aggregated into a single batch, and mapped to outputs for all sizes (*N × N* predictions). Inference uses a single (*d*_in_, *d*_out_) pair. **d)** Loss ablations. Comparison of training losses shows that ChemRank (ChemR) consistently improves retrieval precision (Precision@10) over mean-squared error (MSE) and cosine similarity losses. Performance is further enhanced when building blocks are ordered by SMILES length (BB length) during training, rather than by the original SMARTS reaction order. **e)** Embedding size analysis. For all loss functions, performance is stable for sizes *≥* 256, but drops markedly for sizes *≤* 64.

We evaluated the performance of the Embedding Decomposer using retrieval accuracy at *k* (defined as recovering both target building blocks within the result set) and retrieval precision at *k* (defined by comparing the building blocks retrieved by the predicted embedding to those retrieved by the ground truth embedding). We found retrieval accuracy of 0.973 and retrieval precision of 0.804 at *k* = 10 using 768-dim embeddings (Supp. Fig. 4). Interestingly, we observed that retrieval performance was not permutation invariant to the order of the building block SMILES strings and found that sorting the building blocks by SMILES length resulted in a *∼* 25% performance boost over the default ordering of reactants in the SMARTS string, irrespective of the loss function (Fig. 2d). We investigated failure modes and found most errors were confined to short SMILES strings or building block pairs with near-identical SMILES string lengths (Supp. Fig. 5). Qualitative analysis (Supp. Fig. 16,17,18) shows example molecule decompositions and failure modes. Full details of Embedding Decomposer training and loss ablations can be found at Methods 4.6.

**Fig. 3.**
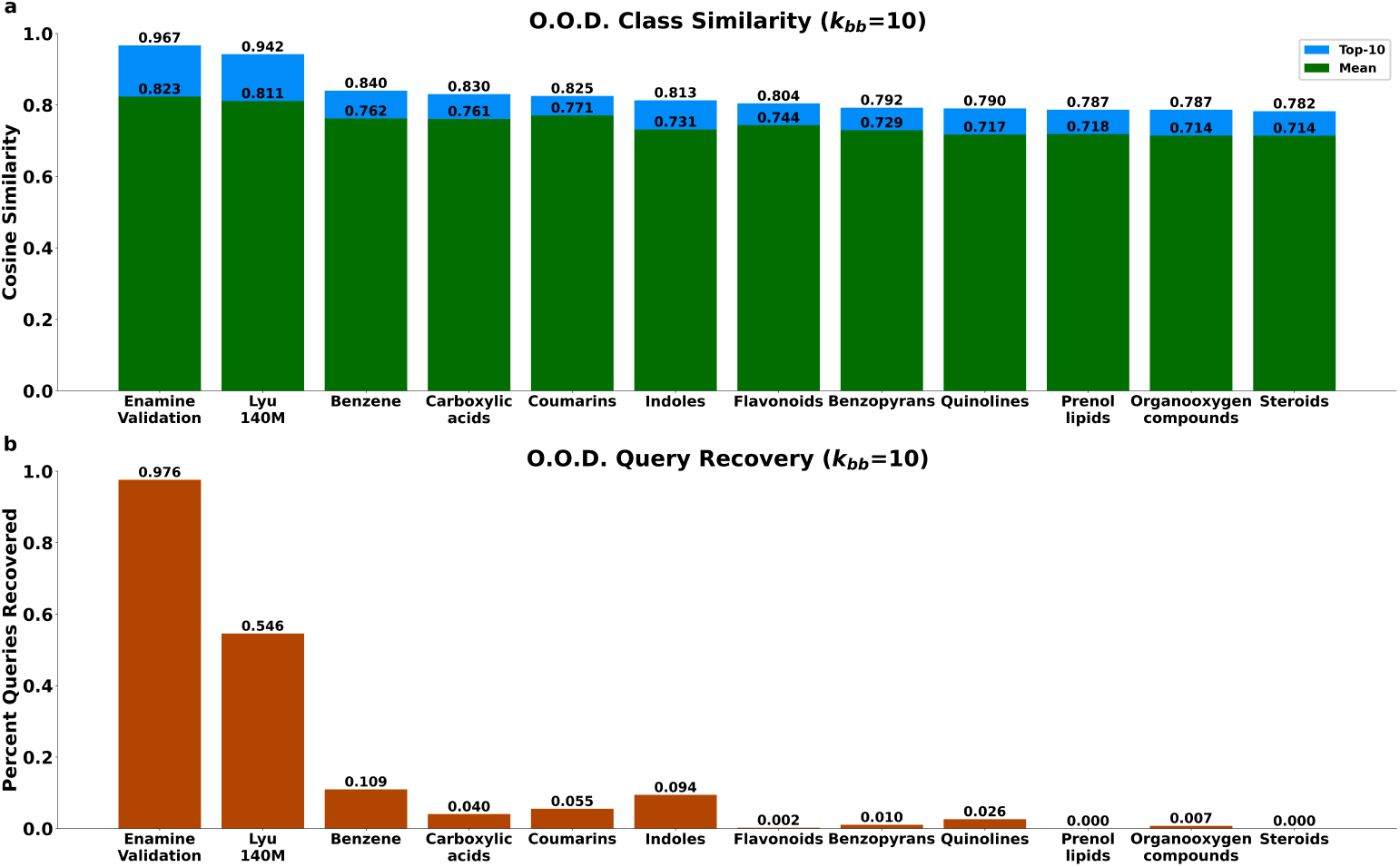
BBKNN generalization to different molecule classes. All evaluation use *k*_bb_ = 10. **(a)** Query to result similarity of all results (green) and the top-*N* = 10 most similar results (blue). “Enamine Validation” denotes molecules from the Enamine Assembled dataset. “Lyu 140M” denotes molecules from the Lyu 140M dataset. All other columns are different natural product classes from the COCONUT 2.0 dataset. **(b)** Average recovery rate (denoting the exact query molecule found in the results set) for different molecule classes.

**Fig. 4.**
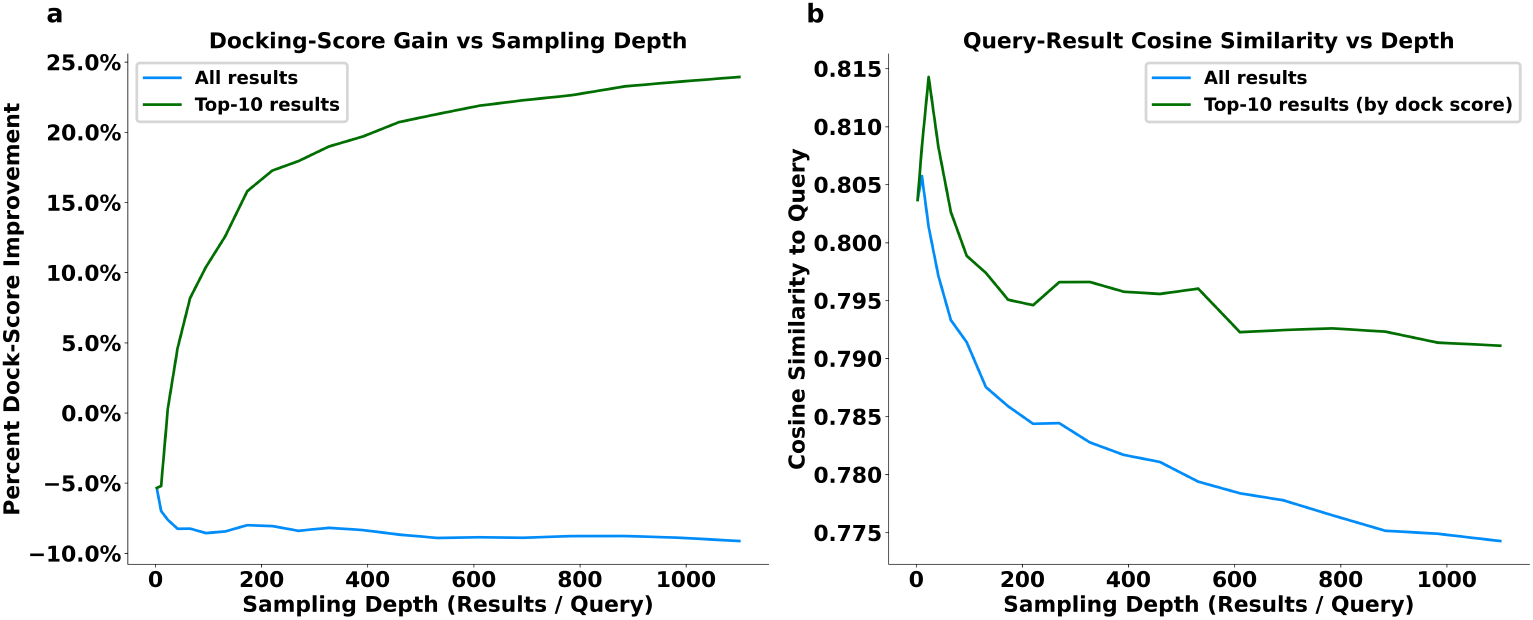
Evaluation of EGFR “analog-by-retrieval” generation capability of BBKNN. Analogs are retrieved from querying 50 known EGFR ligands against the Enamine Building Block dataset. **(a)** Docking score percent improvements over the query molecule vs BBKNN sampling depth. Average results (blue) are 5-10% worse than the query molecule, while the top 10 results in the result set (green) show a docking score improvement of 10% at sampling depth of 100, rising to 24% at sampling depth of 1100. **(b)** Cosine similarity of the results against the query molecule vs sampling depth, also shown for the average (blue) and top-10 (green) results by docking score.

**Fig. 5.**
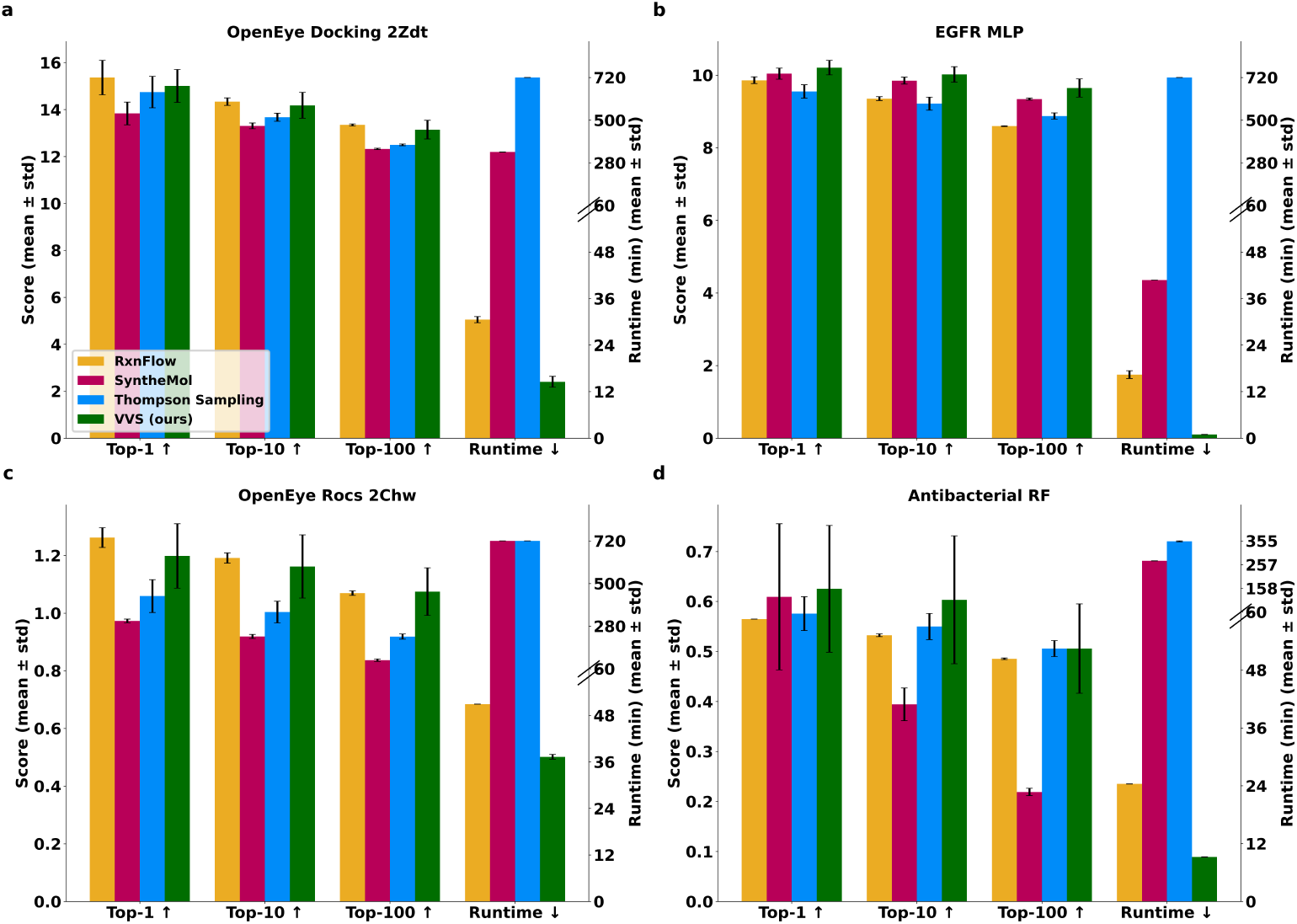
Performance evaluation against various score functions on the Enamine Building Block dataset. All experiments a budget of *≤* 50,000 score evaluations, and a 12-h wall-time limit. Each bar shows the mean ± s.d. of five independent runs for each search method against **(a)** OpenEye Docking, **(b)** EGFR pIC_50_ MLP, **(c)** OpenEye ROCS, and **(d)** Antibacterial Random Forest scoring functions. The “Top- *{*1, 10, 100*}*” reports the average score of the best k molecules found in each run (higher = better ↑). Score magnitudes reflect the specific scoring function used. “Runtime” is the wall-clock time in minutes until the evaluation limit is reached (lower = better ↓); a value of 720 minutes indicates the method exhausted the 12-hour budget before completing all 50k evaluations.

Additionally, the Embedding Decomposer embeddings are used in downstream applications for nearest-neighbor retrieval from a vector database (Results 2.4, 2.5). This process is performance-limited by the embedding dimension (Supp. Fig. 3). To enable ultra-fast and memory-efficient retrieval, we train compression models that reduce RoBERTa-ZINC embeddings from the native 768 dimensions to *d ∈ {* 32, 64, 128, 256, 512*}* (Methods 4.5, Supp. Fig. 1), using the ChemRank loss. Qualitative comparison of embedding retrieval performance across different compression sizes is shown in Supp. Fig. 14, 15.

During development of the compression models, we found that RoBERTa-ZINC embeddings could be compressed from 768 to 128 dimensions with minimal loss in retrieval performance (Supp. Fig. 2). In the final implementation, the Embedding Decomposer is trained jointly across all (*d*_in_, *d*_out_) size combinations (Fig. 2c), enabling flexible use of any compression size in downstream applications. We find performance of the Embedding Decomposer is primarily driven by input embedding size, and the model achieves retrieval accuracy of *≥* 96% and retrieval precision of *≥* 79% at *k* = 10 for embedding input sizes *≥* 256 (Supp. Fig. 4).

### 2.2 Nearest-Neighbor Search in Combinatorial Libraries

The BBKNN algorithm (Methods 4.7, Supp. Algorithm 1) is a *k*-nearest-neighbor procedure that retrieves nearest-neighbor molecules from a building block library without requiring explicit library enumeration of all product molecules. Specifically, for each query molecule *q*, BBKNN (i) uses the Embedding Decomposer model to predict two building block embeddings, (ii) retrieves the *k*_bb_ nearest-neighbors for each predicted embedding from a vector database of purchasable building blocks, (iii) exhaustively cross-combines them using SMARTS reaction templates to yield a set of *N* assembled result molecules, and (iv) ranks the results via embedding cosine similarity to the initial query molecule (Fig. 1a).

On a test set of 1,000 random validation molecules from the Enamine Assembled dataset (Supp. Notes 4.2), BBKNN returns on average 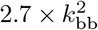 synthesizable analogs with mean cosine similarity to the query of 0.82 at *k*_bb_ = 10 (Supp. Fig. 6). The exact query compound is recovered for 97.5% of queries (Fig. 3b), demonstrating that the BBKNN method can reconstruct a virtual catalog of *∼* 10^10^ product molecules with high fidelity. The fidelity of BBKNN results relies on the underlying Embedding Decomposer model, raising the question of how BBKNN generalizes to molecular scaffolds not present in the Enamine Assembled training dataset. Thus, we test the generalization of the algorithm by applying BBKNN to 1,000 molecules sampled from the Lyu 140M dataset [42] (moderate similarity to the Enamine Assembled dataset used to train the Embedding Decomposer model) and 5,000 natural product molecules from the COCONUT 2.0 dataset [43] sampled from ten structural classes (highly dissimilar to the Enamine Assembled dataset). We find the cosine similarity of BBKNN results is roughly the same for Lyu 140M dataset queries (0.81) and moderately degraded for natural products (0.71–0.76 depending on molecule class), illustrating the ability for BBKNN to retrieve structurally similar molecules for out-of-distribution (O.O.D.) queries (Fig. 3a). However, we find the recovery rate drops moderately for the Lyu 140M dataset sample (55.4%) and substantially for the COCONUT 2.0 dataset (0.0%–10.7% depending on chemical class), as shown in Fig. 3b. The pronounced drop in the recovery rate for natural product molecules is expected, since many query structures contain scaffolds absent from the Enamine Building Block—the source of analog retrieval—and therefore cannot be exactly reconstructed. Qualitative examples of BBKNN retrieval results are shown in Supp. Figs. 19, 20. Full evaluation details can be found in Methods 4.7.2.

**Fig. 6.**
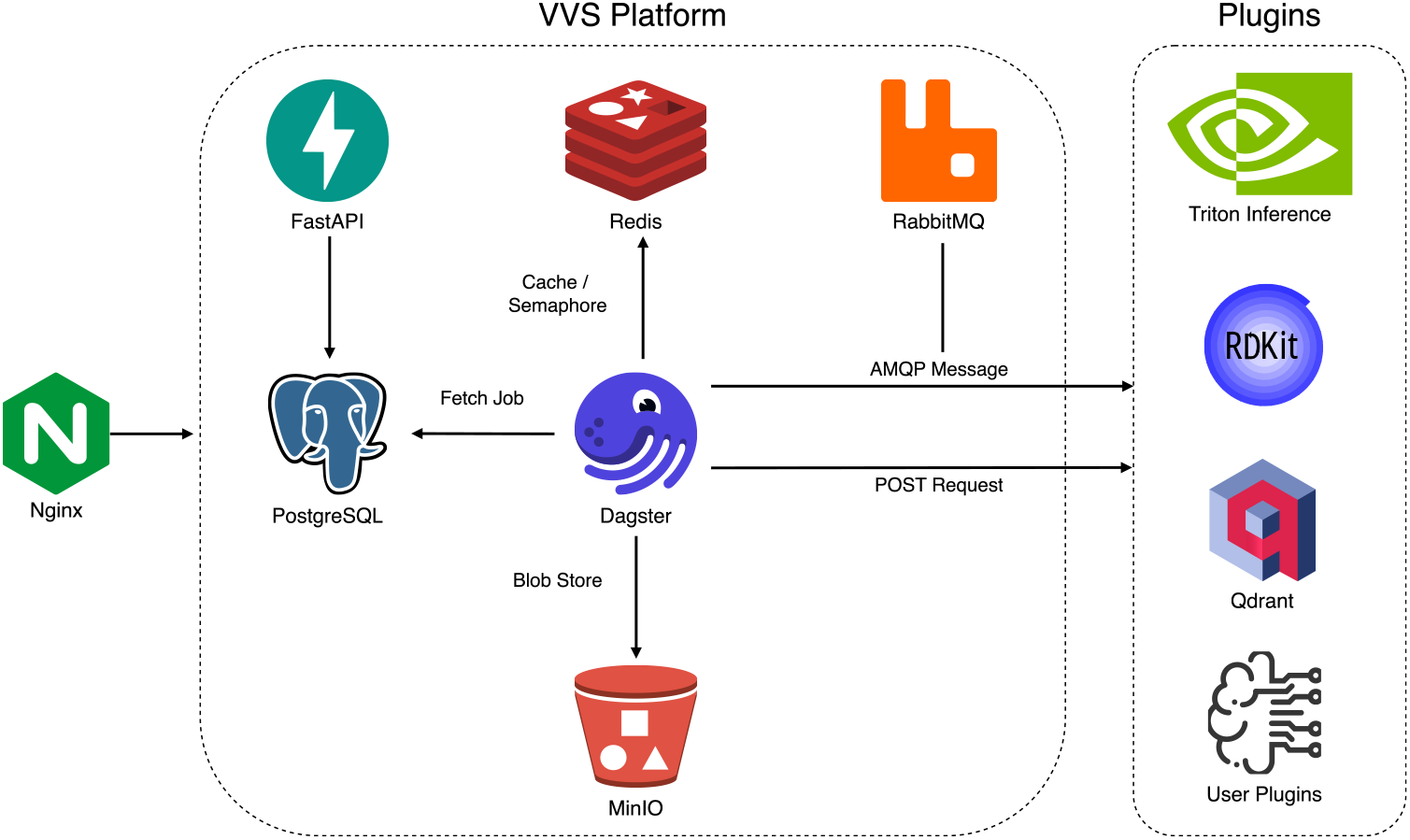
Architecture of the VVS Platform. The core system (left) consists of containerized microservices including a FastAPI server for REST API endpoints, PostgreSQL for state management, RabbitMQ for inter-service messaging, Redis for distributed caching and semaphore control, Dagster for job orchestration, and MinIO for blob storage. The platform supports a flexible plug-in architecture (right) with six canonical roles that can be served via HTTP POST or AMQP messaging. Builtin plugins include Triton Inference Server for neural network models, RDKit for cheminformatics operations, and Qdrant for vector similarity search. Users can extend functionality by deploying custom plugins that adhere to the defined request/response schemas.

### 2.3 Identifying EGFR Analogs by Retrieval with BBKNN

One promising application of BBKNN is to find synthesis-on-demand analogs for arbitrary query molecules. To test this application, we applied BBKNN to 50 known EGFR ligands and evaluated the result molecules using OpenEye molecular docking against the EGFR structure (Methods 4.8). We find that each result set reliably contains molecules that score better than the query. At a sampling depth of 100 results (*k*_bb_ = 5), the top-10 scoring results show an average 10% score improvement over the query, rising to 18% at sampling depth of 270 (*k*_bb_ = 10) and 24% at sampling depth of 1,100 (*k*_bb_ = 20) (Fig. 4). The entire case study was run in a single afternoon on a single consumer CPU with 64 cores. These results demonstrate that BBKNN is a fast and practical “analog-by-retrieval” generator drug design tool that enables medicinal chemists to turn arbitrary query molecules into synthesis-on-demand analogs available for immediate purchase. Example analogs are shown in Supp. Fig. 23.

### 2.4 Virtual Screening with VVS

Virtual screening is the task of computationally identifying small molecules that maximize a target scoring function. To enable virtual screening of ultra-large libraries without enumeration, we introduce the VVS algorithm (Fig. 1b): a gradient-directed hill climbing algorithm for efficient search of combinatorial chemical spaces that rapidly identifies synthesis-on-demand molecules that optimize an arbitrary scoring function. To accomplish this, VVS algorithm extends BBKNN by incorporating a key insight: evaluating the BBKNN nearest-neighbor results with the target scoring function enables estimation of the gradient of the score function with respect to the query embedding (Supp. Algorithm 2). These gradients then guide an iterative, multi-scale search (Supp. Algorithms 3, 4).

In brief, given a query embedding **q**, corresponding gradient **g**, and a list of learning rates *α*_*j*_ *∈* ***α***, VVS generates multiple “gradient step queries” via **q**^(*j*)^ = **q** *− α*_*j*_**g** where each **q**^(*j*)^ is an embedding representing a hypothetical molecule in continuous latent space. Each **q**^(*j*)^ is used as the query to BBKNN, which resolves the hypothetical embeddings back into discrete synthesis on-demand molecules. The retrieved results are then scored with the target scoring function. The embedding of the top-scoring result is selected as the next query, and the gradient of the new query is estimated via Supp. Algorithm 2. A detailed description of the algorithm can be found in Methods 4.9.

During the development of VVS, we conducted hyperparameter sweeps over 1,400 independent VVS searches using the EGFR pIC_50_ MLP scoring function to explore the impact of different parameters on performance (Supp. Fig. 9). Notably we find that multiple gradient steps per iteration are critical to performance, with a maximum learning rate of 5,000 yielding the best results (Supp. Figs. 9b,d). Learning-rate sweeps (Supp. Fig. 8) show the VVS method can tolerate high learning rates due to the retrieval properties of the BBKNN method-BBKNN always returns valid molecules, even for “off manifold” queries, enabling large exploratory jumps in chemical embedding space without divergence. Finally, the 1,400 ablation runs produced a cumulative set of 50 million assembled molecules. In analyzing this dataset, we find 83% of these results were unique (Supp. Fig. 10), underscoring that VVS consistently produces reliably diverse results.

Next, we compared VVS (using the 256-dim compressed embeddings) against three state-of-the-art search methods that operate on building block libraries without enumeration: SyntheMol [44], Thompson Sampling (TS) [8], and RxnFlow [45], against four diverse scoring functions: OpenEye docking score against the c-Jun N-terminal kinase 3 (JNK3) crystal 2ZDT [46], OpenEye 3D ROCS similarity to the 46C ligand bound to 2ZDT, a random-forest antibiotic classifier [44], and a 13M-parameter MLP predicting pIC_50_ for EGFR (Supp. Notes 4.4) on the Enamine Building Block building block dataset with results summarized in Fig. 5, Supp. Table 5. Each method was allowed 50,000 molecule evaluations and a 12 hour wall time on equal hardware. For each method, we first perform hyperparameter sweeps using the EGFR pIC_50_ MLP scoring function (Supp. Table 3) and use the best parameters for final evaluation. See Methods 4.3 and Methods 4.10 for details on the scoring function and benchmark implementations.

Across all scoring functions, VVS achieves substantially shorter run-times than other methods, completing searches 1.4*×* –17*×* faster than the next fastest method and several orders of magnitude faster than the slowest (Fig. 5, Supp. Table 5). Molecule quality, measured by the top- *{* 1, 10, 100*}* scores, shows that VVS is consistently best or competitive with the best method, indicating that its speed advantage does not come at the expense of molecule quality. The magnitude of this improvement is most pronounced for fast scoring functions (e.g., EGFR pIC_50_ MLP and antimicrobial RF), where method overhead constitutes a larger fraction of the total runtime, further underscoring the efficiency of the core method. Supp. Figs. 12,13 show the evolution of the top-100 score for each benchmark run as a function of inference budget and runtime, respectively.

For completeness, we also evaluate VVS on enumerated spaces, replacing the BBKNN step with standard k-nearest-neighbors, using an HNSW index for nearest neighbor retrieval (Supp. Fig. 11, Supp. Table 6). We compare VVS on an enumerated set of 100M molecules from the Lyu 140M dataset against the Retrieval Augmented Docking method (RAD) [47]. We observe that VVS delivers performance on par with RAD, showing only a slight reduction in average top- *{*1, 10, 100 *}* retrieval scores—while consistently running 2.2 *×* –15*×* faster than RAD. Together, these results establish that VVS not only accelerates searches on enumerated molecular libraries without compromising performance, but also bridges building block and enumerated libraries, underscoring its versatility and utility.

### 2.5 VVS Platform

In addition to the core VVS algorithm, we also release the VVS Platform, a lightweight, scalable, plug-and-play software tool designed to scale to the growing size of combinatorial libraries (Fig. 6). At its core is a plug-in architecture that turns the system into a “bring-your-own-algorithm” sandbox. The Platform recognizes six canonical plug-in roles—embedding, data-source, filter, score, mapper, and assembly—each conforming to a strict request/response schema that can be served over HTTP POST or as a RabbitMQ routing-key–based consumer. To add a new method, users deploy a lightweight FastAPI microservice or RabbitMQ worker on the same network; each plug-in declares its batch size, concurrency, and timeout so utilization can be tuned to available hardware.

The distribution includes several built-in plug-ins: a Qdrant vector database [48] for HNSW search [49]; an RDKit bundle for SMARTS-based filtering, reaction assembly, and property scoring; and a Triton GPU inference server for the RoBERTa-ZINC, Embedding Compression, and Embedding Decomposer models. Integration of Embedding Compression and Embedding Decomposer is particularly advantageous at scale, where compressed embeddings substantially reduce query overhead during vector-database queries (Supp. Fig. 3).

Operationally, the Platform ships as a Docker Compose stack in which each capability is a containerized microservice that can be scaled independently: a FastAPI server exposes a public REST API, PostgreSQL stores state, RabbitMQ handles AMQP messaging, and an Nginx reverse proxy fronts external traffic. VVS hill-climbing workflows are orchestrated by Dagster [50]. Within each workflow, a shared Redis instance serves as both a distributed cache and a semaphore layer that enforces concurrent access limits on plug-in resources, allowing the core backend to remain stateless while ensuring usage limits are respected across processes.

We anticipate that this modular, cloud-ready architecture will facilitate rapid adoption of VVS across the virtual screening community by lowering deployment barriers and enabling seamless integration with diverse computational environments. More broadly, the platform’s scalability and plug-in design position VVS as a flexible foundation that can readily incorporate future scoring functions, workflows, and advances in machine learning—ensuring its continued relevance as chemical space and screening technologies evolve.

## 3 Discussion

The development of synthesis-on-demand catalogs has expanded virtual libraries from millions to tens of billions of compounds in just a few years. This growth has outpaced brute-force enumeration, making ultra-large virtual screening intractable with traditional methods. At the same time, increasingly compute-intensive scoring functions exacerbate the challenge, underscoring the need for efficient molecular search tailored to combinatorial libraries.

To address these challenges, we propose **Vector Virtual Screen** (VVS), which leverages the recent advent of chemical language models to reframe ultra-large virtual screening as *vector search in building-block space*. Using gradient-directed hill-climbing, VVS locates high-value molecules in enumerated space at the computational cost of building block search. On the Enamine Building Block benchmark, VVS identified promising molecules while evaluating only 50,000 candidates out of 10^10^ possible, running orders of magnitude faster than comparable baselines (Fig. 5).

These results are enabled by four innovations: the Embedding Decomposer model, which learns “latent retrosynthesis” to map molecules to building block embeddings; the ChemRank loss, which improves retrieval precision; the BBKNN algorithm, which performs efficient nearest-neighbor search in building block space without enumeration; and gradient estimation in CLM embedding space, enabling hill climbing. Together, these make billion-scale catalogs tractable on a single compute node.

### Implications for drug discovery

BBKNN retrieves close analogs of arbitrary molecules, while VVS rapidly optimizes combinatorial libraries under any black-box score function. Both return molecules accessible through synthesis-on-demand services, allowing medicinal chemistry teams to generate purchasable hit lists in a single workday and accelerate the cycle from in silico design to laboratory validation.

### Limitations

The performance of VVS and BBKNN is bounded by the quality of four inputs: the RoBERTa-ZINC used for embeddings, the building-block catalog, the SMARTS-based assembly rules, and the scoring function. Search coverage is confined to the chemistry spanned by the available building blocks and reaction templates. Expanding the chemical domain of the Embedding Decomposer model to new reactions and building blocks may require retraining. The scoring function governs how well in silico gains transfer to laboratory outcomes, limiting the efficacy of VVS results. Ultimately, performance scales with the quality of chemical embeddings, scoring functions, and building block coverage. Thus, improvements in these areas will directly enhance the power of Embedding Decomposer, BBKNN and VVS.

### Future Work

Extensions to multi-component, multi-step syntheses (Supp. Notes 4.10) suggest a path to broader chemical coverage. Future work could integrate VVS and BBKNN into established pipelines—such as GB-GA or active learning surrogates—and ultimately couple them with experimental feedback loops to create closed design–synthesis–test cycles.

## Reproducibility Statement

All code to download or create datasets, implement scoring functions, and run benchmarks can be found at https://github.com/DarkMatterAI/VVS_benchmarks. Code for the VVS Platform can be found at https://github.com/DarkMatterAI/VVS_platform.

## 4 Methods

### 4.1 Datasets and Chemical Libraries

We utilize several chemical datasets to train models and evaluate our algorithms:

(i) Enamine Building Block dataset from Swanson et al. [44], consisting of 130,000 Enamine building blocks and 12 two-component SMARTS assembly reactions, (ii) Lyu 140M dataset from Lyu et al. [42], consisting of 1.4 *×* 10^8^ compounds generated from 130 reactions and 70,000 Enamine building blocks, (iii) Zinc 10M dataset, consisting of 10M molecules from the ZINC-22 library [51], (iv) COCONUT 2.0 from Chandrasekhar et al. [43], consisting of 700,000 natural product molecules, (v) Chembl ErbB1, consisting of 7,000 molecules with measured binding affinities (IC_50_) against the epidermal growth factor receptor (EGFR) downloaded from the ChEMBL [52] database, and (vi) Enamine Assembled dataset (this work), consisting of 50M molecules assembled from the Enamine Building Block building blocks and reactions. See Supp. Notes 4.2 for a detailed overview of dataset sources, creation, and processing.

### 4.2 Chemical Language Model

Molecule embeddings are generated using the RoBERTa-ZINC model [41], a 102M parameter Roberta [53] style masked language model trained on 480M SMILES strings from the ZINC database [51].

### 4.3 Scoring Functions

To comprehensively compare the performance of VVS against other methods, we implement four qualitatively different scoring functions representing a diverse range of computational methods relevant to drug discovery. Details on the implementation of each score can be found in Supp. Notes 4.3.

- Structure-based virtual screening: protein-ligand docking to the JNK3 crystal structure 2ZDT
- Ligand-based virtual screening: Rapid Overlay of Chemical Structures (ROCS) to measure 3D shape and electrostatic similarity against a reference ligand pose from Klarich et al. [8]
- Classical machine learning: A random-forest classifier, trained on 2-D fingerprints to predict antibacterial activity, sourced from Swanson et al. [44]
- Modern deep learning: A 13M-parameter multilayer perceptron trained on the Chembl ErbB1 set to predict EGFR binding affinity from RoBERTa-ZINC embeddings (Supp. Notes 4.4)

### 4.4 ChemRank Loss

The Chemical Ranking (ChemRank) loss trains embeddings to preserve *relative* similarity patterns by maximizing the Pearson correlation between predicted and target similarity profiles. Given two similarity matrices **S**^pred^, **S**^target^ *∈* ℝ ^*N ×M*^ —each row lists the cosine similarities of one query embedding to the same set of *M* reference points—the loss is defined as:

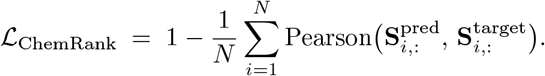

Because the loss operates on *similarity* rather than raw vectors, ChemRank is agnostic to embedding scale and dimensionality, enabling fair comparison across models with different latent sizes. The use of *correlation* over direct value comparison (i.e. MSE loss) prioritizes relative similarity ranking over exact value reconstruction, producing robust *k*-nearest-neighbors retrieval performance (Fig. 2d,e, Supp. Fig. 2). We propose two variants of the loss, based on the origin of the similarity matrices:

- **Self-similarity**. Predicted embeddings 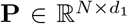 and target embeddings 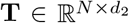 are each correlated against themselves, yielding **S**^**P**,**P**^ and **S**^**T**,**T**^. Diagonal elements (trivial self-similarities of 1) are masked. The embedding dimensions *d*_1_ and *d*_2_ may differ.
- **Reference-similarity**. Both **P** and **T** are compared to an external dictionary **R** *∈* ℝ^*M ×d*^ to obtain **S**^**P**,**R**^ and **S**^**T**,**R**^. All three sets must share the same dimension *d*, or the reference embeddings **R** must be available in both the dimensionality of **P** and **T**.

To further improve retrieval performance, we give additional weight to the top *k ∈ {* 10, 100, 256*}* highest similarity values for each row. We find ChemRank with top-*k* weighting consistently outperforms other loss functions for embedding-based retrieval (Fig. 2d,e, Supp. Fig. 2).

### 4.5 Embedding Compression Model

We observed empirically that embedding size impacts embedding retrieval latency (Supp. Fig. 3). To investigate the trade-off between retrieval speed and retrieval quality, we trained a lightweight autoencoder to compress the 768-dim RoBERTa-ZINC embeddings to five target sizes (*d ∈ {32, 64, 128, 256, 512}*) using the ChemRank loss to preserve the nearest-neighbor structure of the native embedding. The Embedding Compression model has an encoder/decoder structure (Supp. Fig. 1a), first compressing the native 768-dim CLM embedding to the target compression size with the encoder, and reconstructing the original input embedding with the decoder.

We train the Embedding Compression model on a dataset of 30M molecules created by sampling 10M molecules from the Zinc 10M, Lyu 140M, and Enamine Assembled datasets using ChemRank loss on the compressed embeddings from the encoder with top-*k* weighting for the top *k ∈ {* 10, 100, 256*}* most similar pairs and cosine similarity loss on the reconstructed embeddings from the decoder (Supp. Fig. 1b). We evaluated the model on a dataset of 1.5M validation molecules—1M of the validation molecules are used as a retrieval corpus, and the remaining 0.5M act as queries. The retrieval precision for the top-100 results (*p*@100) is computed for compressed embeddings using the retrieval results from the native embedding as the ground truth.

We conducted an ablation study on our different loss metrics and found that ChemRank loss with top-*k* weighting outperforms other metrics, providing a 5% performance boost over MSE loss with top-*k* weighting (Supp. Fig. 2a). We find that compressed embedding dims 128–512 have high retrieval precision (*>* 0.89 *p*@100) relative to the native embeddings, while performance falls off for dims 64 and 32, which achieve 0.77 and 0.59 *p*@100 respectively (Supp. Fig. 2c). In addition, we provide a qualitative assessment of retrieval performance for specific molecule queries (Supp. Fig. 14, 15). Further details on dataset creation, training, and loss ablations are provided in Supp. Notes 4.5.

### 4.6 Embedding Decomposer Model

The Embedding Decomposer model uses a product molecule CLM embedding to predict two building block embeddings corresponding to molecules that assemble into the product molecule, “decomposing” the product molecule into building blocks in latent space. The model consists of a single FFN_SwiGLU_ “in-projection” layer [54] that maps the input embedding to a 1024 dim hidden space, followed by a trunk of nine FFN_SwiGLU_ layers and an “out-projection” linear layer that predicts two output embeddings for each input embedding (Supp. Fig. 2a). The model is trained to handle all combinations of embedding dimensions in *d ∈ {* 32, 64, 128, 256, 512, 768 *}*, using different in-projection and out-projection layers for the different sizes (Supp.Fig. 2c). The Embedding Decomposer model has 43.6M total parameters.

The Embedding Decomposer model is trained on the Enamine Assembled which consists of (bb_1_, bb_2_, product) tuples. We preprocess the dataset by ordering the target building blocks in length order, such that len(bb_1_) *≤* len(bb_2_). We find that this simple heuristic improves model performance by *∼* 25% (Fig. 2d,e).

During training, the model processes a batch of 2,048 product molecule embeddings at the native size (768-dim) and all compression sizes (32, 64, 128, 256, 512). The in-projection layers map all inputs to the common hidden dimension of 1024. All inputs are processed through the trunk as an aggregate batch and then processed into different output sizes by the out-projection heads. Each input size is used to predict each output size, resulting in 36 distinct output predictions. For each output prediction, we compute the cosine similarity of the predicted building block embeddings to the target embeddings. We then randomly sample 3,072 building block “reference embeddings”. We compute **S**_pred,ref_ and **S**_targ,ref_, the pairwise similarities of the predicted and target embeddings against the reference embeddings, and compute the ChemRank loss with *k ∈ {* 10, 100*}* weighting on the similarity matrices (Supp. Fig. 2b). We evaluate the Embedding Decomposer on a hold-out set of 1M product molecule embeddings. We decompose the hold-out embeddings with all input size to output size combinations and retrieve *k* = 10 nearest neighbor results from a pre-computed set of building block embeddings. We compute the retrieval precision of these results by comparing to the retrieval results produced by the ground truth target embeddings.

We perform ablations on our different loss functions and find ChemRank loss with top-*k* weighting combined with cosine similarity gives the best performance (Fig. 2d,e). We evaluated the performance of the best model in terms of retrieval precision at *k* and retrieval accuracy at *k* (defined as the percent of queries that retrieve both ground truth building blocks within *k* results) across different input and output size combinations. We find that input size is a strong driver of performance, with input sizes 128-dim and above achieving 92%+ retrieval accuracy at *k* = 5, while 64-dim and 32-dim input sizes show 86% and 53% retrieval accuracy respectively (Supp. Fig. 4).

We assess different failure modes, and find the decomposer model struggles with disambiguating small molecules with SMILES strings *≤* 8 characters (Supp. Fig. 5b). We find that when both target building block SMILES string lengths are within *≤* 1 characters of each other, the model may “flip” the prediction, predicting the correct embeddings in the wrong order (Supp. Fig. 5a). Additionally, we provide a qualitative assessment of retrieval performance for specific molecule queries (Supp. Fig. 16, 17, 18). Full training details and loss ablations are provided in Supp. Notes 4.6.

### 4.7 Building Block K-Nearest-Neighbors (BBKNN)

The Building Block K-Nearest-Neighbors (BBKNN) algorithm enables efficient embedding-based retrieval of molecules from combinatorial libraries without requiring explicit library enumeration. The algorithm leverages the Embedding Decomposer model to decompose a query molecule embedding into constituent building block embeddings, which are then used to retrieve similar building blocks from a vector database. These retrieved building blocks are subsequently combined through in silico synthesis to generate the final result molecules.

#### 4.7.1 BBKNN Algorithm

The BBKNN algorithm (Supp. Algorithm 1, Fig. 1a) comprises the following steps:

**1. Embedding Decomposition**: Given a query embedding, we use the Embedding Decomposer model to predict two building block embeddings corresponding to the constituent parts of the query molecule.

**2. Building Block Retrieval**: For each of the two predicted building block embeddings, we query a vector database of precomputed building block embeddings to retrieve the *k*_bb_ nearest-neighbors, resulting in two pools of *k*_bb_ building blocks each.

**3. In silico Synthesis**: The retrieved building blocks are combinatorially assembled using a defined set of SMARTS-based reaction rules. For each pair of building blocks (one from each pool), we attempt to apply all compatible reaction rules, resulting in a set of assembled result molecules.

**4. Result Ranking**: The assembled molecules are embedded using the same embedding model as the query embedding. Result molecules are ranked by their embedding cosine similarity to the query molecule.

For implementation, we use the building blocks and reactions from the Enamine Building Block dataset. We embed the building blocks with the RoBERTa-ZINC model and compress to 256-dim with the Embedding Compression model. We use the Embedding Decomposer model with 256-dim input size and 256-dim output size and store the building block embeddings as a HNSW index using Usearch [55]. In-silico synthesis is performed using RDKit’s reaction SMARTS functionality.

#### 4.7.2 BBKNN Evaluation

We first evaluate the baseline behavior of BBKNN as a function of the retrieval parameter *k*_bb_. We sample 1,000 random molecules from the Enamine Assembled validation split and query these molecules against the Enamine Building Block dataset using BBKNN with *k*_bb_ *∈ {*1, … 20*}*. Across this range the number of assembled products grows roughly as 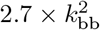, while the mean query–product cosine similarity drops from 0.89 (*k*_bb_ = 1) to 0.80 (*k*_bb_ = 20) (Supp. Fig. 6).

Next, we evaluate the generalization of the method under distribution shift. Distribution shift can impact the BBKNN method in two different ways: (i) the Embedding Decomposer may produce poor predicted embeddings for query molecules that are significantly structurally different from those seen during training, and (ii) the query molecule may contain features or substructures not present in the building block dataset used for retrieval and assembly. To probe robustness, we evaluated three query sets with increasing dissimilarity to the Embedding Decomposer training data:

- **In-distribution:** 1,000 products from the Enamine Assembled validation set.
- **Slightly OOD:** 1,000 molecules from the Lyu 140M, assembled from the same building blocks but with a different reaction set.
- **Strongly OOD:** 5,000 natural products sampled from ten structural classes in COCONUT 2.0 (stereochemistry removed to match the model’s training settings).

Each query was processed with *k*_bb_ = 10. Results were assessed via (i) cosine similarity computed on 256-dim CLM embeddings and (ii) recovery rate, the fraction of queries for which the exact molecule was reconstructed.

- Enamine Assembled queries: mean cosine similarity 0.82; recovery 97.5%.
- Lyu 140M queries: mean cosine similarity 0.81; recovery 54.5%.
- COCONUT 2.0 queries: mean cosine similarity 0.76–0.71; recovery 0–10.9% (both values are class-dependent).

Detailed class-wise numbers, similarity distributions, and representative query–result pairs are presented in Fig. 3, Supp. Figs. 19, 20.

### 4.8 BBKNN EGFR Case Study

Fifty known EGFR inhibitors from Chembl ErbB1 were docked against the EGFR structure (PDB: 6LUD) using the OpenEye docking procedure described in Supp. Notes 4.3. Each ligand was used as a BBKNN query against the Enamine Building Block dataset with *k*_bb_=20 using 256-dim embeddings. The assembled products (totaling 47,566 unique molecules) were then docked against EGFR with the same settings.

We find that each set of results reliably contains molecules that score better than the query. At a sampling depth of 100 results (*k*_bb_ = 5), the top-10 scoring results show an average 10% score improvement over the query, rising to 18% at sampling depth of 270 (*k*_bb_ = 10) and 24% at sampling depth of 1100 (*k*_bb_ = 20) (Fig. 4), demonstrating BBKNN as a practical “analog-by-retrieval” generator for lead-optimization. Example analogs are shown in Supp. Fig. 23.

### 4.9 VVS Algorithm

The VVS algorithm extends the local retrieval capabilities of BBKNN into a gradient-directed search that maximizes an arbitrary scoring function *f* (·) within a combinatorial chemical space. The core insight of VVS is that a set of scored results from BBKNN can be used to estimate the gradient of the target function with respect to the query embedding, allowing for directional movement in the embedding space. This approach enables efficient hill-climbing without requiring exhaustive enumeration of the library.

#### 4.9.1 VVS Inner Loop

The inner loop of the VVS algorithm (Supp. Algorithm 3) is executed as follows. Let **q** *∈* ℝ ^*d*^, **g** *∈* ℝ^*d*^ be the query embedding and its estimated gradient (initially **g** = **0**). For a user-defined set of learning rates ***α*** = *{*0, *α*_1_, …, *α*_*n*_*}*, we generate “gradient step queries” via **q**^(*j*)^ = **q** *− α*_*j*_**g**, run BBKNN on each **q**^(*j*)^, and score the results. This yields a set of combined results 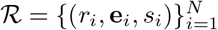 where *r*_*i*_, **e**_*i*_, and *s*_*i*_ = *f* (*r*_*i*_) are the result molecule, molecule embedding, and molecule score, respectively. The embedding of the top scoring result 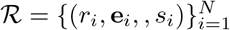 is selected as the new query 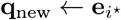, and the remaining results 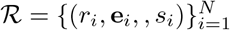 are used to estimate the gradient **g**_new_ at **q**_new_, which become the inputs to the next iteration. The process is run concurrently for a batch of *B* query embeddings for a user-defined number of iterations. We use “top-1 update” to refer to the process of selecting 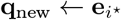 and computing the gradient at **q**_new_, in contrast to the “standard” hill-climbing approach of estimating the gradient at the original query embedding **q**.

#### 4.9.2 Advantage Weighted Loss and Gradient Estimate

The gradient of the new query is estimated with Supp. Algorithm 2. First, scores are converted to centered, normalized advantages.

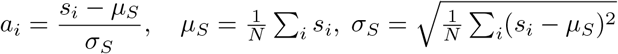

With cosine distance *d*_*i*_ = 1 *−* cos *⟨* **q**_new_, **e**_*i*_ *⟩*, we compute the surrogate loss of advantage-weighted distances:

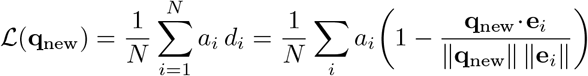

Differentiating with respect to **q**_new_ yields the gradient estimate:

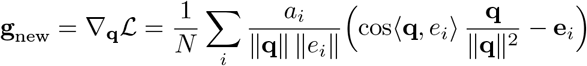

#### 4.9.3 VVS Outer Loop

The core VVS algorithm is repeated until a user-defined wall clock limit or inference limit is reached (Supp. Algorithm 4). Each new batch is initialized with *B* queries, of which *p*_explore_ percent is sampled randomly and *p*_exploit_ = 1 *− p*_explore_ is sampled from high-scoring molecules from the previous iteration. A complete overview of hyperparameters and ablations can be found in Supp. Notes 4.8.

### 4.10 VVS Benchmarks

To quantify the performance of VVS, we compared it against three state-of-the-art search methods for molecule optimization in building block libraries without enumeration: Synthemol [44], Thompson Sampling [8], and RxnFlow [45]. All methods were tasked with discovering high-scoring molecules drawn from the Enamine Building Block dataset when optimizing four qualitatively different scoring functions - OpenEye docking score against the human JNK3 (PDB: 2ZDT), OpenEye ROCS similarity to a target reference ligand, a random-forest antibiotic classifier [44], and a 13 M-parameter MLP predicting pIC_50_ binding affinity against EGFR (see Supp. Notes 4.3 for score implementation details). Each run was capped at 50k molecule evaluations and 12 wall-clock hours. Each method is implemented using the author’s published code, with as few alterations as possible. All methods were afforded identical hardware (64 CPU cores, 1 × A100 GPU); however, not every codebase was designed to use GPU compute or multicore CPU processing. We report the actual hardware used by each method in Supp. Table 4. Because algorithmic performance is sensitive to internal meta-parameters, we first conducted a full hyper-parameter sweep for every method against the EGFR pIC_50_ MLP scoring function objective; the best scoring set of parameters (by the average rank order of the top- *{*1, 10, 100*}* scores) was then frozen and applied to all four objectives with five replicas per objective. For each scoring function, we report the mean score of the top-1, top-10, and top-100 molecules (Fig. 5, Supp. Table 5). Top-100 scores for each method and objective are shown as a function of total inference and runtime in Supp. Fig. 12,13. For completeness, we also evaluate the VVS algorithm on enumerated spaces by comparing to the RAD method [47] on a subset of 100M molecules from the Lyu 140M dataset (Supp. Fig. 11, Supp. Table 6) to showcase the performance of the method at 10^8^ enumerated scale using a hierarchical navigable small worlds (HNSW) index [49] for nearest-neighbor searching in the enumerated space. Full benchmark implementation details can be found in Supp. Notes 4.9.

## Supplementary

### 1 Supplementary Figures

**Supplementary Fig. 1.**
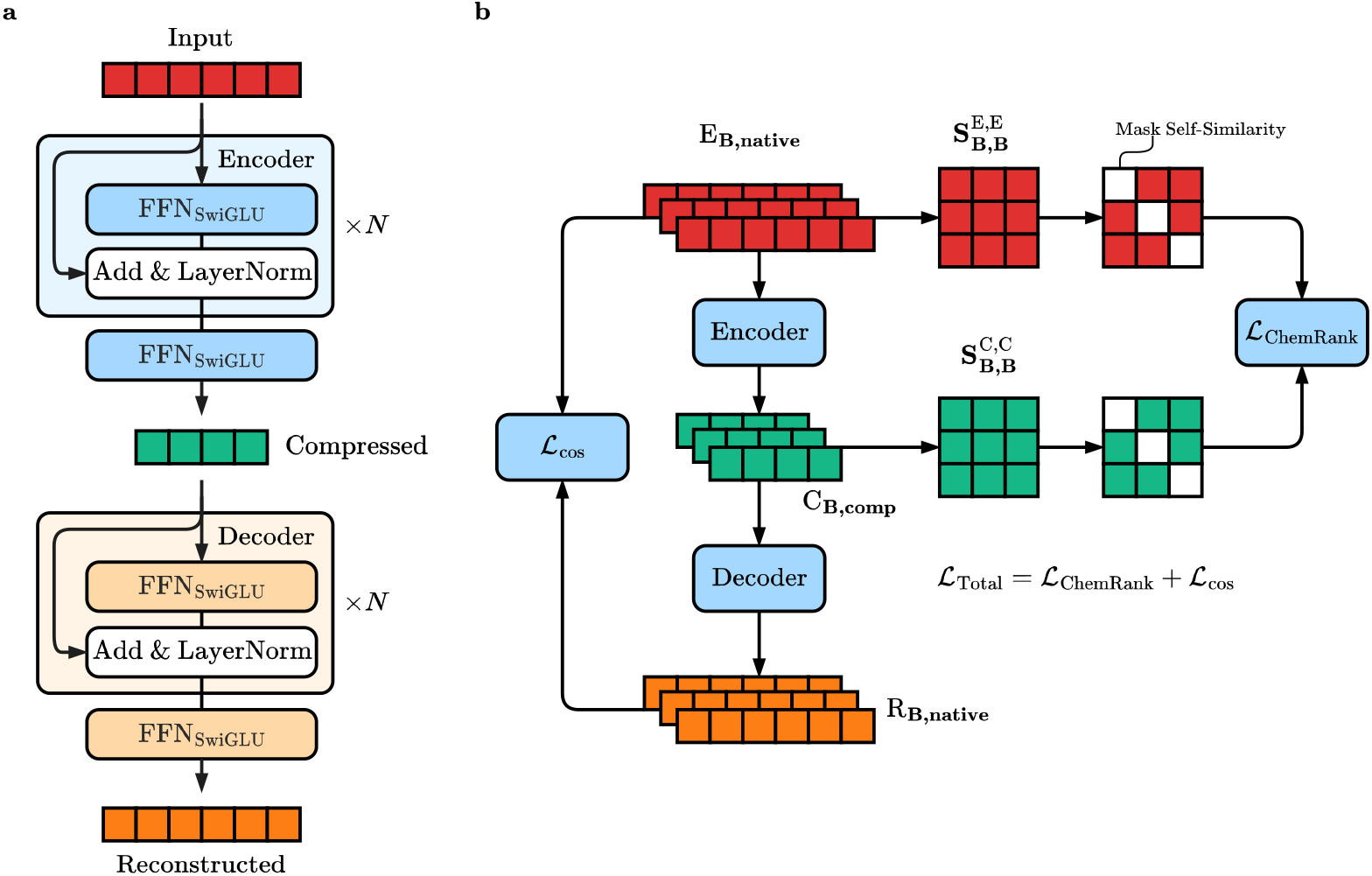
Embedding Compression model framework. **a)** Model architecture of the compression encoder and decoder, consisting of *N* blocks of FFN_SwiGLU_ layers followed by skip connection and LayerNorm, with a final FFN_SwiGLU_ layer to project the activations into the output space. The encoder maps the input embedding from the input dimension **d**_**native**_ to a compressed embedding with dimension **d**_**comp**_, and the decoder reconstructs the original embedding from the compressed embedding. **b)** Model training framework. A batch of input embeddings **E** (red) with dimension (**B, native**) is compressed by the encoder to compressed embeddings **C** (green) and reconstructed by the decoder to an output **R** (orange). We compute the cosine similarity loss *ℒ*_cos_ between the input and reconstructed embeddings. We compute in-batch cosine similarity of all pairs in **E** and **C** to yield batch similarity matrices **S**^E,E^ (red square) and **S**^C,C^ (green square), each with dimension (**B, B**). We mask the diagonal self-similarity values and compute the ChemRank loss *ℒ* _ChemRank_. The final loss *ℒ* _Total_ is the sum of *ℒ* _cos_ and *ℒ* _ChemRank_.

**Supplementary Fig. 2.**
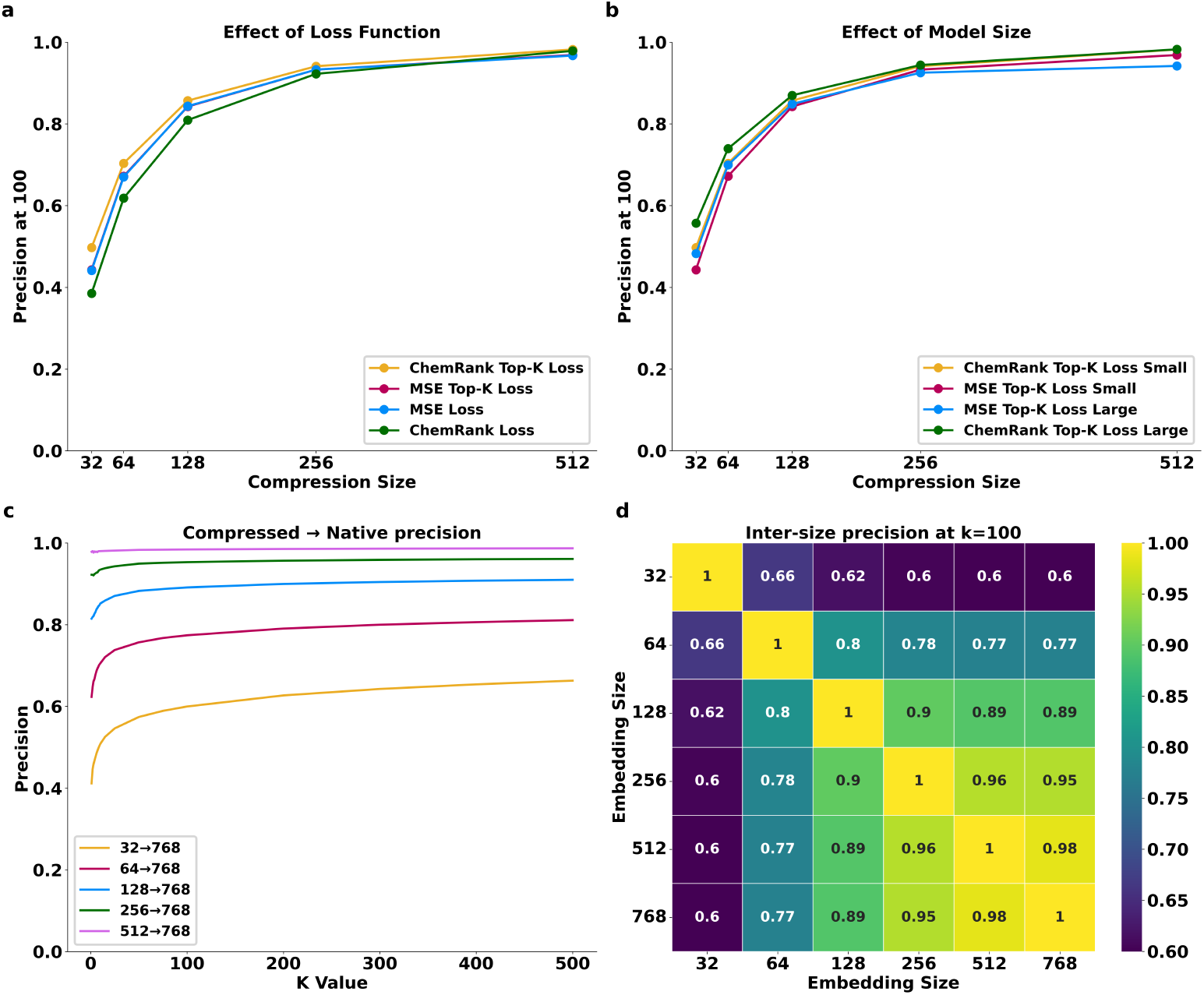
Ablations and evaluations for the embedding compression model. **(a)** Loss ablations comparing Precision@100 versus compressed embedding size for four loss functions on the compressed embedding - ChemRank, ChemRank with top-*k* weighting, MSE loss and MSE loss with top-*k* weighting for *k* = 10, 100, 256. All runs additionally use cosine similarity loss on reconstructed embeddings from the decoder. ChemRank with top-*k* weighting performs the best with an average 3%improvement over MSE loss with top-*k* weighting across all compression sizes. **(b)** Effect of model depth comparing a single-layer compression model (denoted small) and a four-layer compression model (denoted large) using ChemRank or MSE loss, both with top-*k* weighting. We see a modest improvement with depth, with the ChemRank large model showing a 2.8% improvement over the small model. **(c)** Precision vs neighbor cut-off *k* (1-500) comparing each compressed size to the native 768-dim baseline. Larger compression sizes consistently track close to the baseline, highlighting a tradeoff between the degree of compression and loss of information. **(d)** Heat-map of cross-size agreement (Precision@100) - each cell shows the fraction of top-100 neighbors shared between two compression sizes. We see high agreement for dimensions 128 and higher (*≥* 0.89), with a fall-off for dimensions 64 and lower.

**Supplementary Fig. 3.**
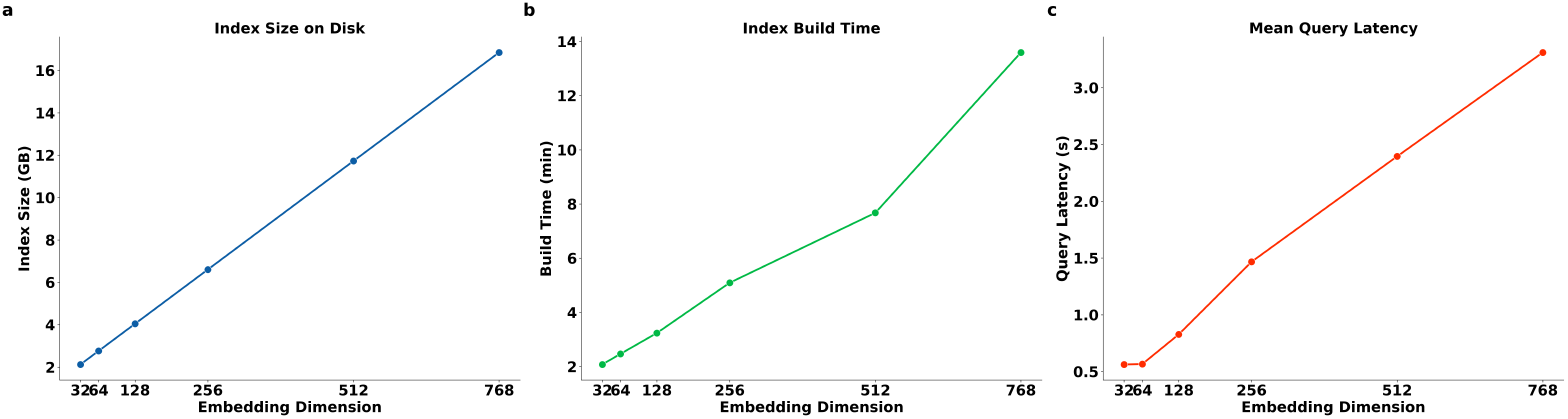
HNSW Retrieval performance as a function of embedding size. 10M molecules from the Zinc 10M dataset were embedded at all compression sizes and built into HNSW indices using Usearch [55]. Each index was evaluated on total index size, index build time, and query latency. **(a)** Index size on disk (GB) as a function of embedding size. **(b)** Index build time (minutes) as a function of embedding size. **(c)** Average query latency as a function of embedding size for a query of 1,024 embedding vectors with *k* = 100.

**Supplementary Fig. 4.**
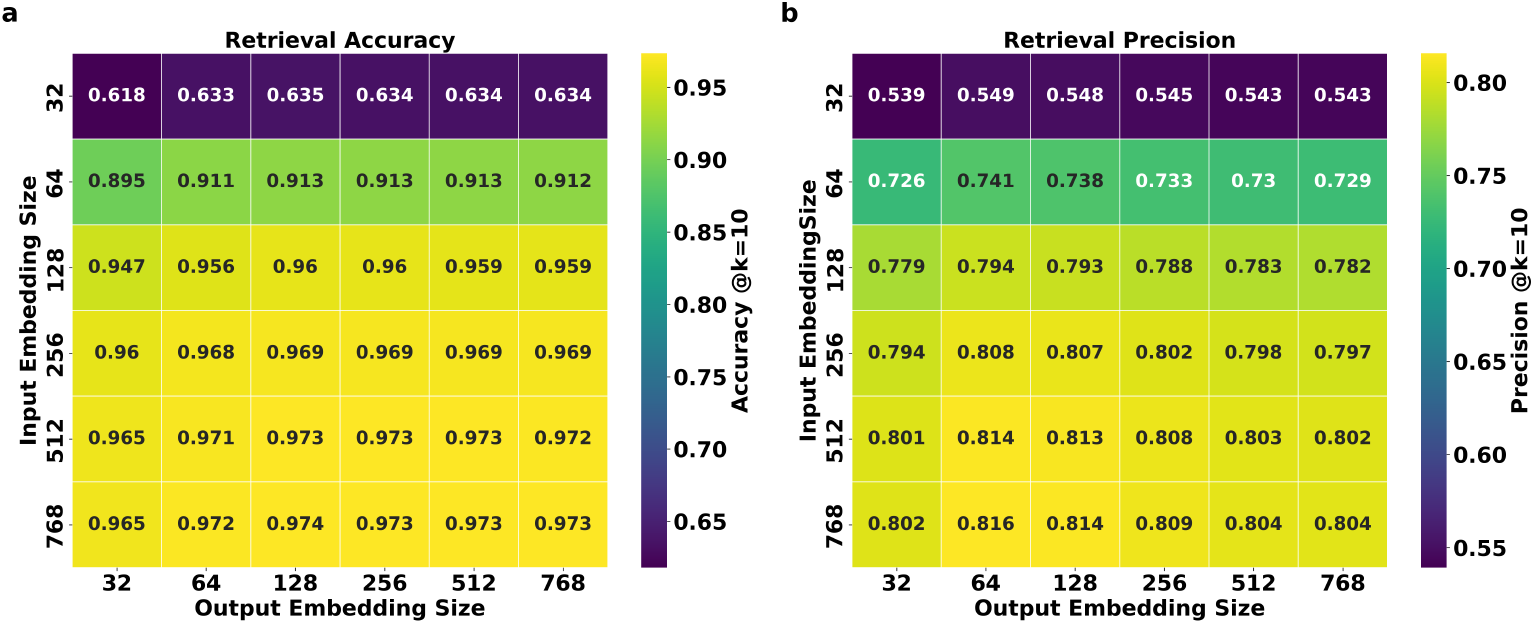
Retrieval performance for the Embedding Decomposer model using the model trained with Cosine and ChemRank loss. **(a)** Heatmap of retrieval accuracy (recovering both ground truth building blocks) at *k* = 10 for all input size to output size combinations. **(b)** Heatmap of retrieval precision at *k* = 10 (defined by comparing the building blocks retrieved by the predicted embedding to those retrieved by the ground truth embedding) for all input size to output size combinations. For both metrics, performance is dominated by the input embedding size. Input sizes 128 and above show roughly the same performance, while input sizes 64 and below show noticeably worse performance.

**Supplementary Fig. 5.**
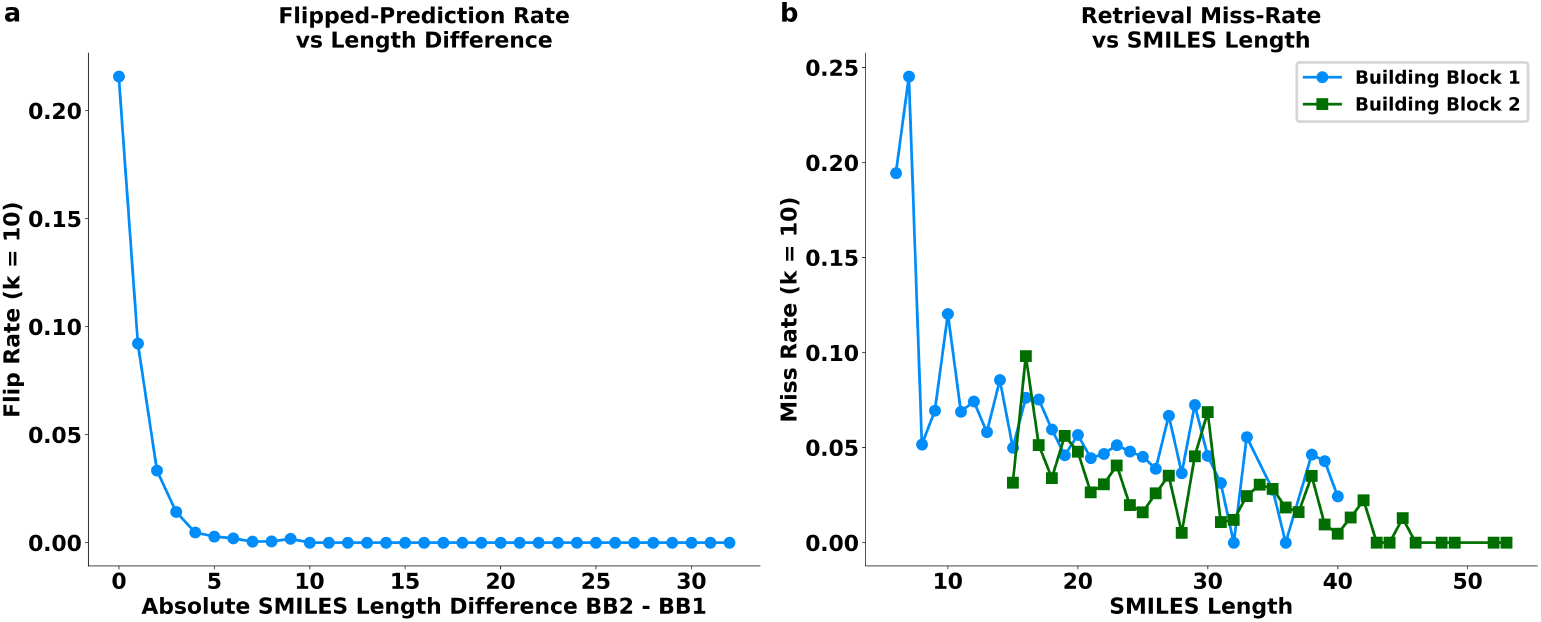
Embedding Decomposer model error modes vs building block length. **(a)** Flipped-prediction rate (both building blocks found but swapped) versus building block SMILES string length difference, showing a 21% flip rate when both building block lengths are exactly equal, dropping off to zero as the length difference increases. **(b)** Miss rate at *k* = 10 (neither block found in top-10 retrieval results) versus building block SMILES string length. Errors concentrate on very short building blocks (*≤* 8 characters).

**Supplementary Fig. 6.**
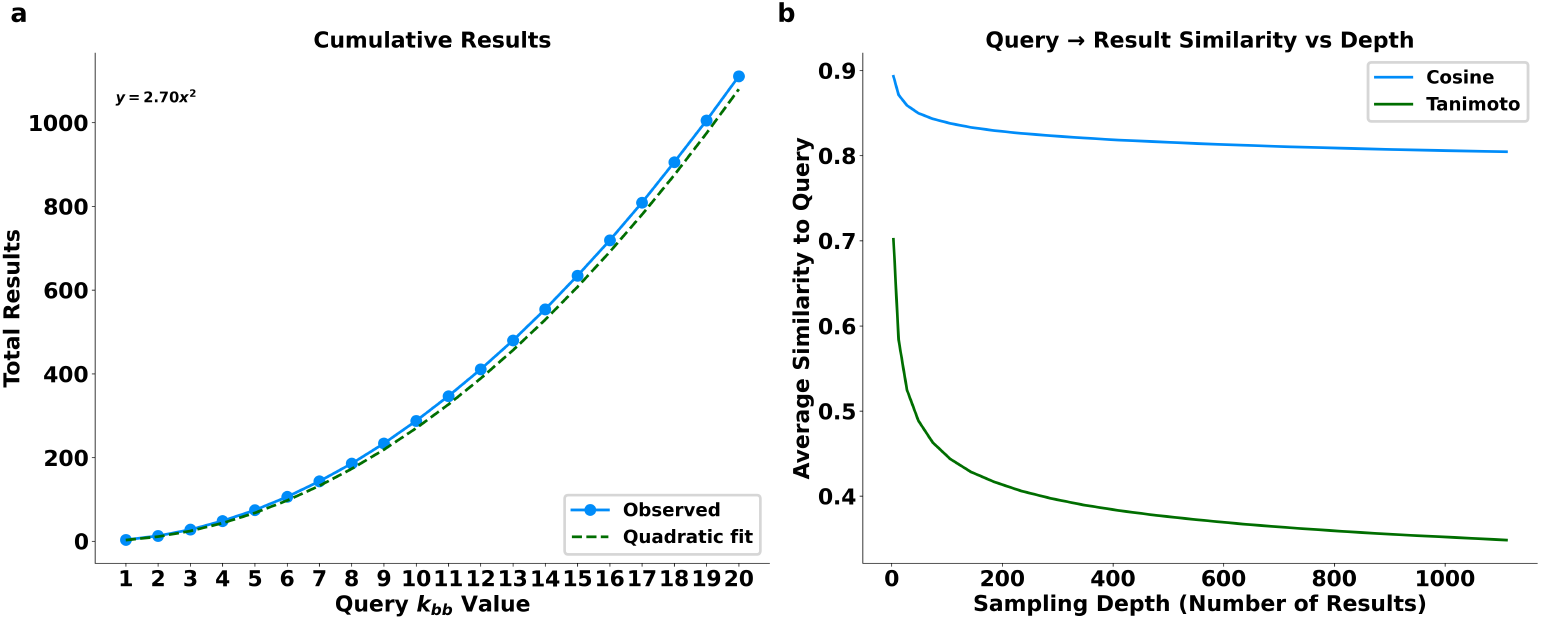
BBKNN Evaluation. **(a)** Result set size vs query *k*_bb_ value, showing approximately 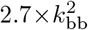 results per query. **(b)** Average query to result Tanimoto and cosine similarity vs sampling depth. Results were generated using 256-dim input and output size for the Embedding Decomposer on a set of molecules from the Enamine Assembled dataset.

**Supplementary Fig. 7.**
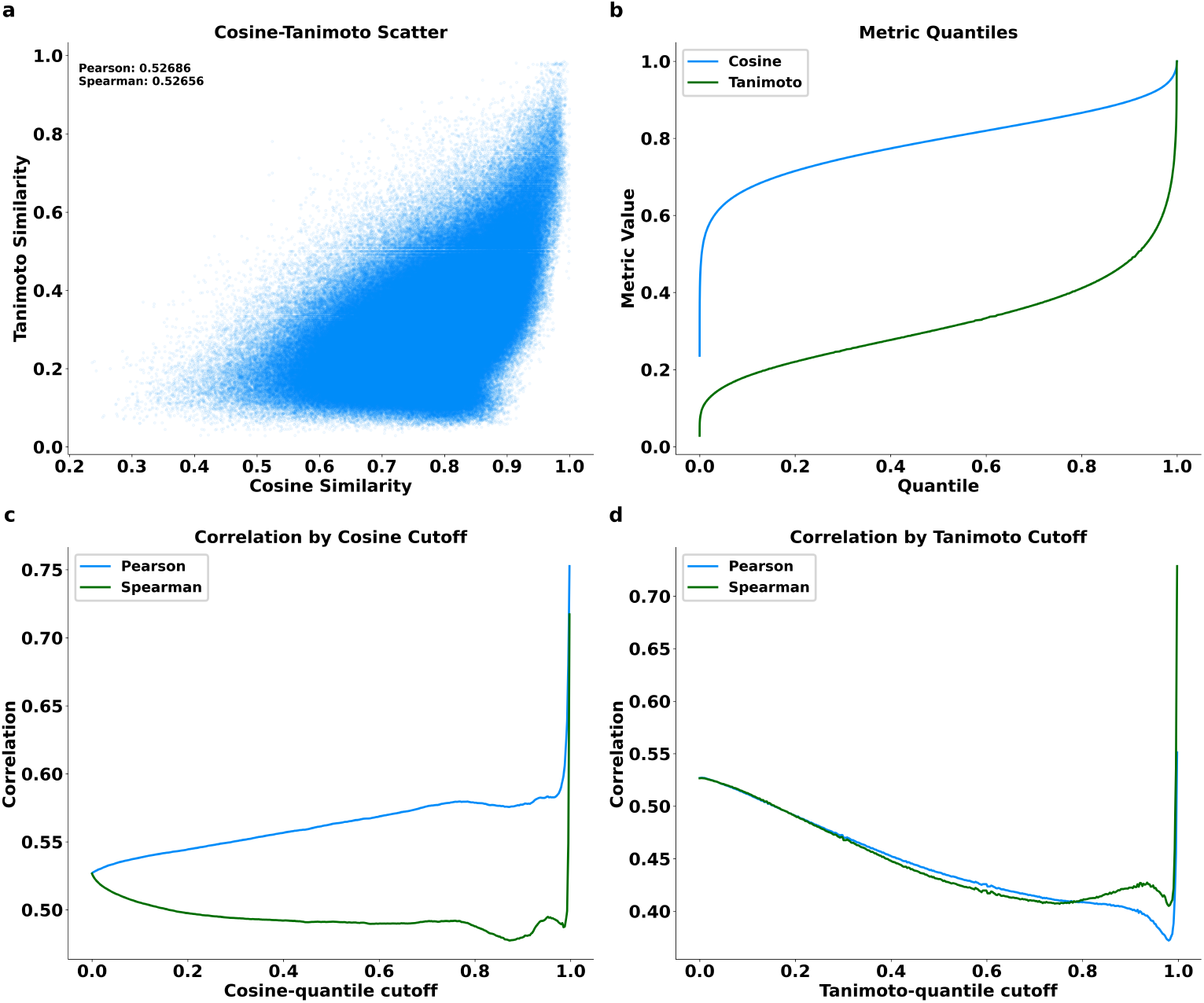
Comparison of CLM embedding cosine similarity (256-dim compressed embeddings) and Tanimoto similarity (ECFP-4) using 2.3M molecule pairs. **(a)** Scatter plot of cosine similarity and Tanimoto similarity (750,000 points shown). **(b)** Quantile curves for cosine and Tanimoto similarity. **(c)** Pearson and Spearman correlations between cosine and Tanimoto similarity at different cutoffs based on cosine similarity quantiles. **(d)** Pearson and Spearman correlations between cosine and Tanimoto similarity at different cutoffs based on Tanimoto similarity quantiles.

**Supplementary Fig. 8.**
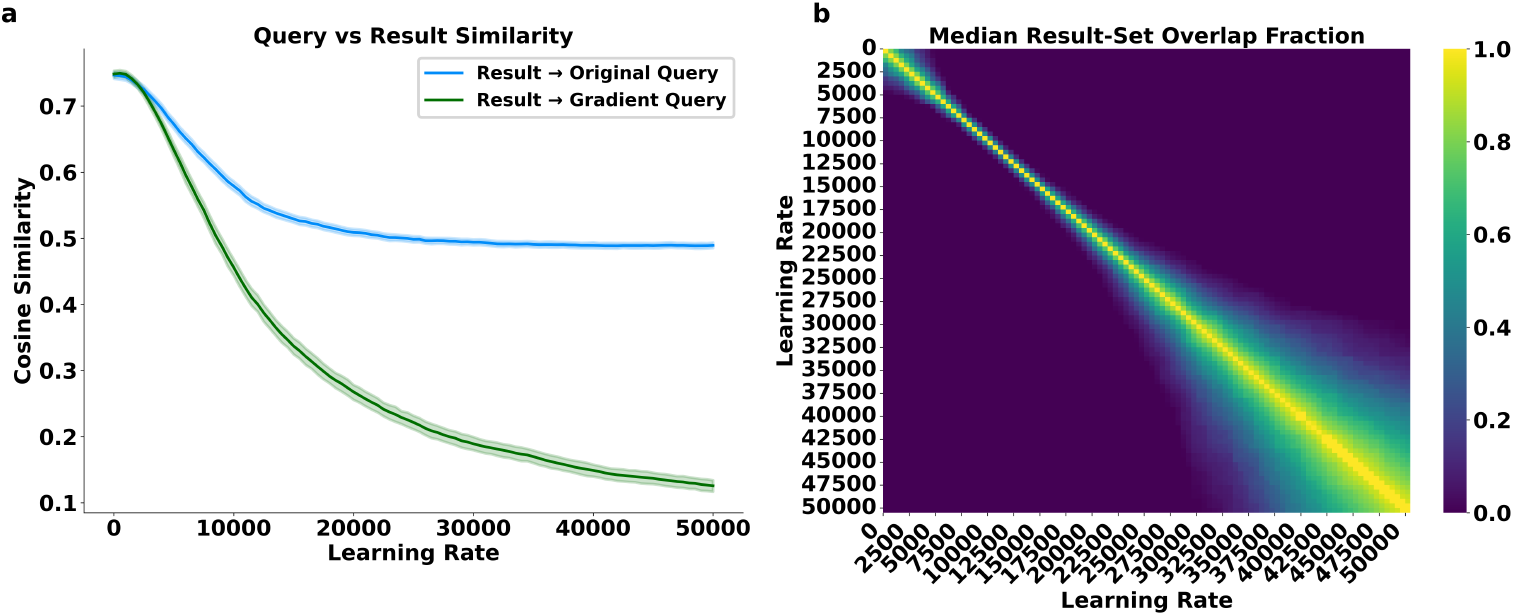
VVS learning rate sweep. 256 query embeddings with gradients were used to generate 101 gradient steps with learning rates ranging from 0 to 50, 000 in increments of 500 and used to retrieve a set of result molecules using BBKNN. **(a)** The average cosine similarity of the result molecules compared to the original query (blue) and the gradient step query (green) generated by taking a step along the gradient defined by *q*_*step*_ = *q*_*orig*_ *− α * g*. Result similarity to the original query drops modestly (0.75 *→* 0.5) while result similarity to the gradient step query plummets to 0.12. This shows that large gradient steps move “off-manifold”, but still generate valid “on-manifold” results due to the explicit retrieval of the BBKNN algorithm. **(b)** result set overlap for gradient step queries at different learning rates. Result set overlap is local (*≤ ±* 2, 000 learning rate) until *α ≈* 30, 000, where the overlap range grows much wider.

**Supplementary Fig. 9.**
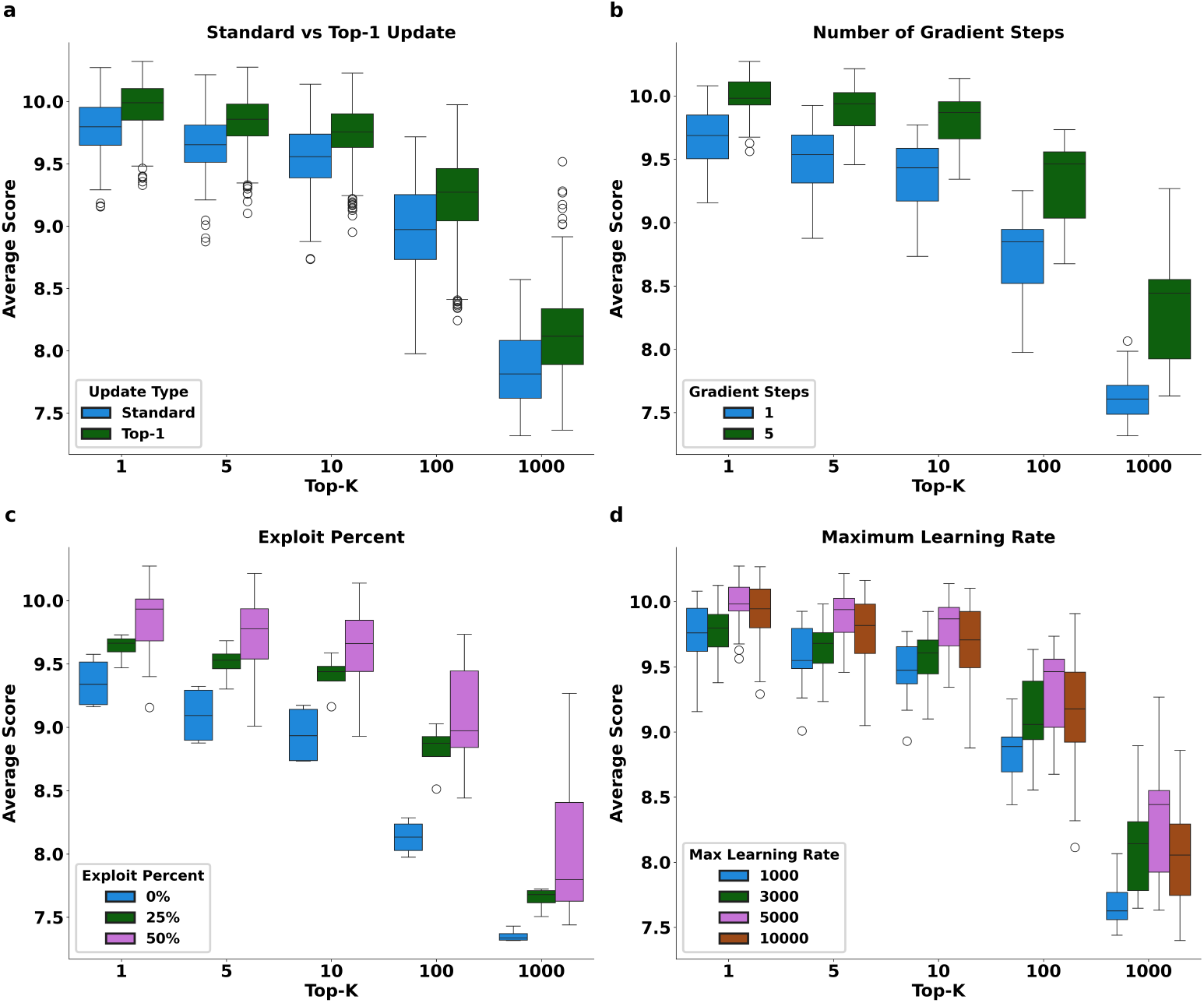
VVS hyperparameter evaluation with the EGFR score function. Each boxplot shows the average top *k* results across 1,400 independent VVS searches. **(a)** scores stratified by update type. “Standard” refers to standard hill climbing, where we estimate the gradient of the initial query embedding. “Top-1” refers to the VVS approach of selecting the top-1 scoring result in each batch as the new query embedding, and estimating the gradient at that value. **(b)** scores stratified by number of gradient steps for a single step vs multiple steps, showing improved scores from using multiple gradient steps for each VVS iteration. **(c)** scores stratified by *p*_exploit_ for the outer VVS loop, showing improved performance from re-sampling high scoring results from previous iterations. **(d)** scores stratified by maximum learning rate, showing improvement from 1,000 to 5,000, but no benefit from 5,000 to 10,000.

**Supplementary Fig. 10.**
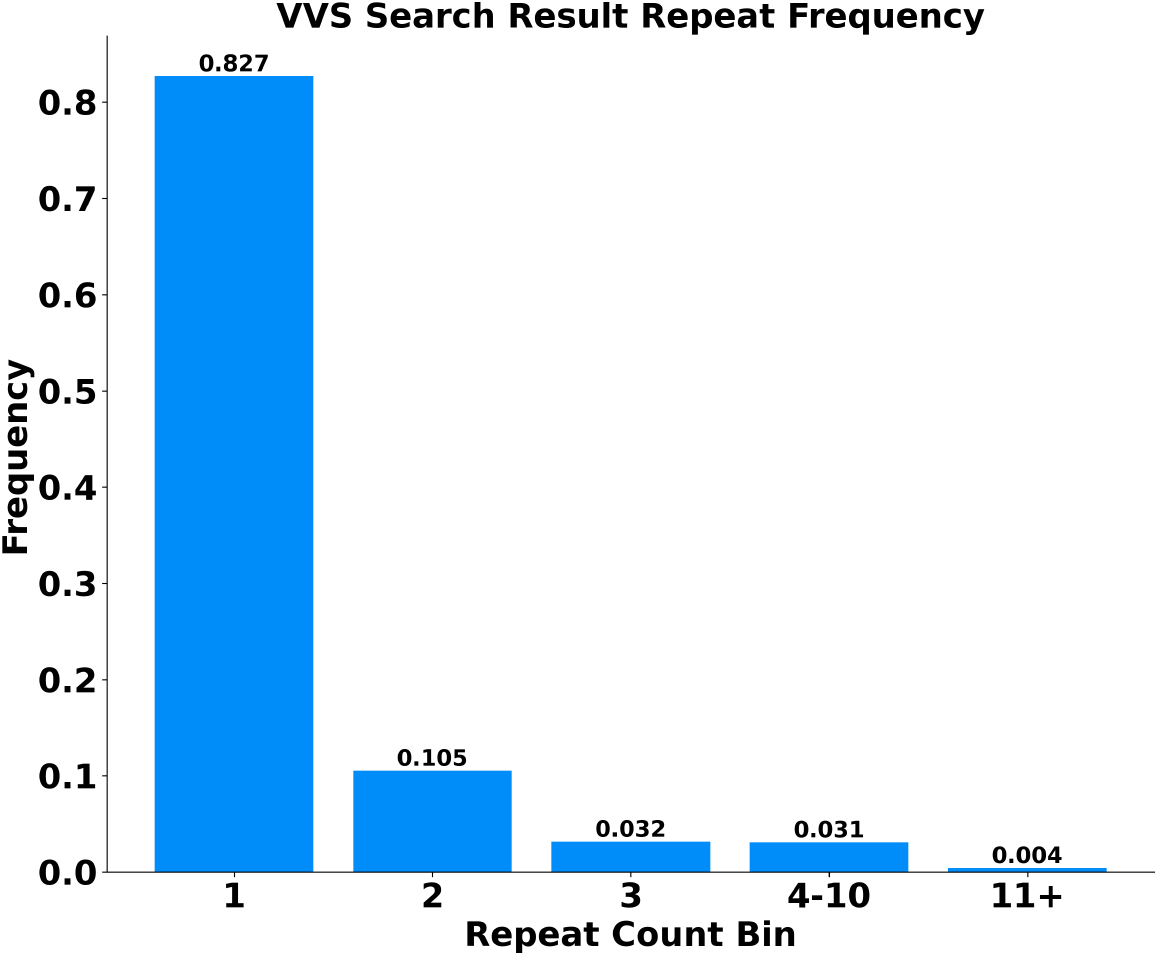
VVS result diversity of 50M unique result molecules generated over 1,400 VVS searches using the EGFR pIC_50_ MLP scoring function. The vast majority of molecules (82.7%) appear exactly once, indicating that VVS reliably produces diverse results.

**Supplementary Fig. 11.**
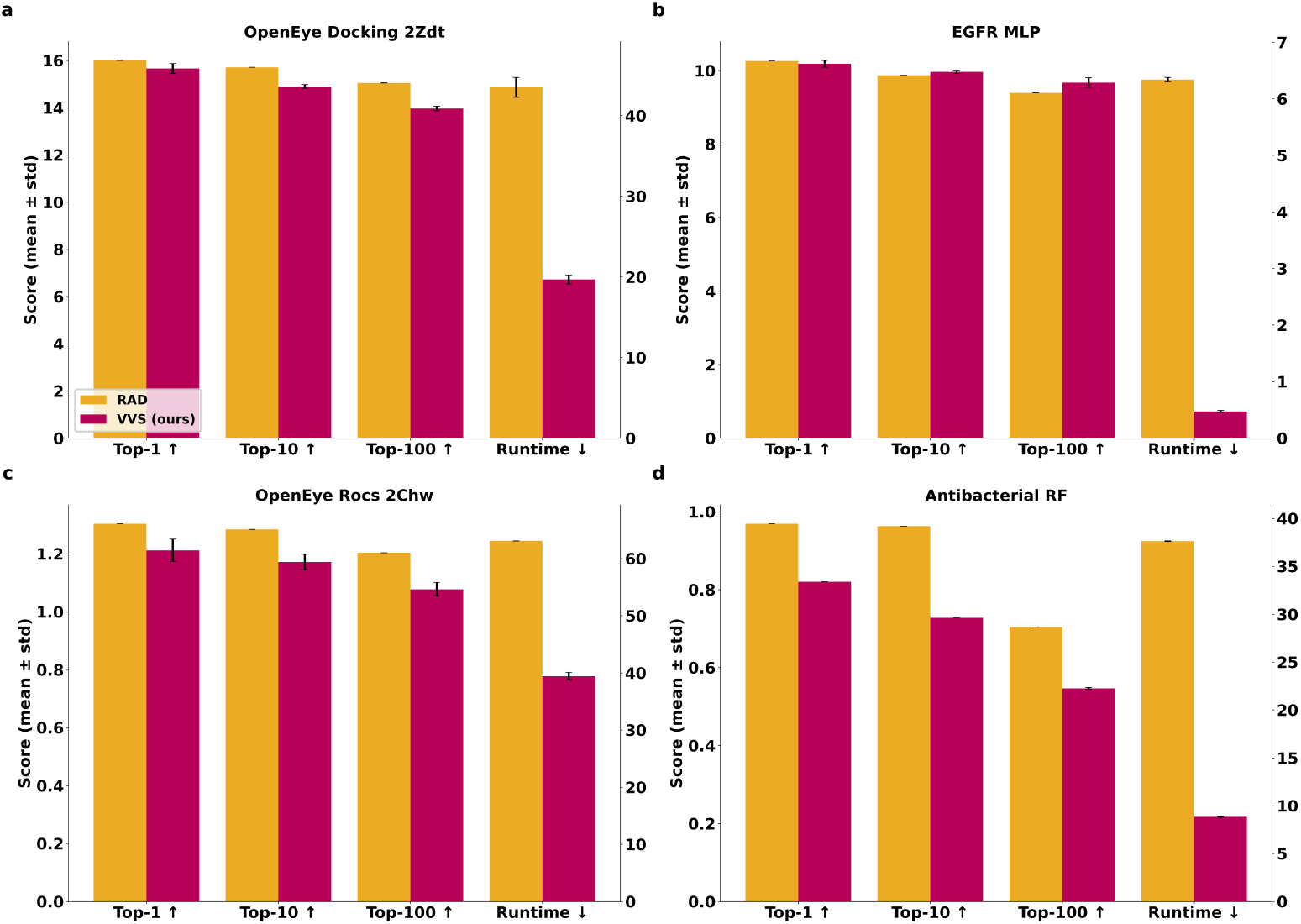
Performance of VVS and RAD search strategies on the Lyu 140M under a budget of 50,000 score evaluations and a 12-h wall-time limit. Each bar shows the mean ± s.d. of five independent runs for each search method against **(a)** OpenEye Docking, **(b)** EGFR pIC_50_ MLP, **(c)** OpenEye ROCS, and **(d)** Antibacterial Random Forest scoring functions. The “Top- *{*1, 10, 100*}*” reports the average score of the best k molecules found in each run (higher = better ↑). Score magnitudes reflect the specific scoring function used. “Runtime” is the wall-clock time in minutes until the evaluation limit is reached (lower = better ↓).

**Supplementary Fig. 12.**
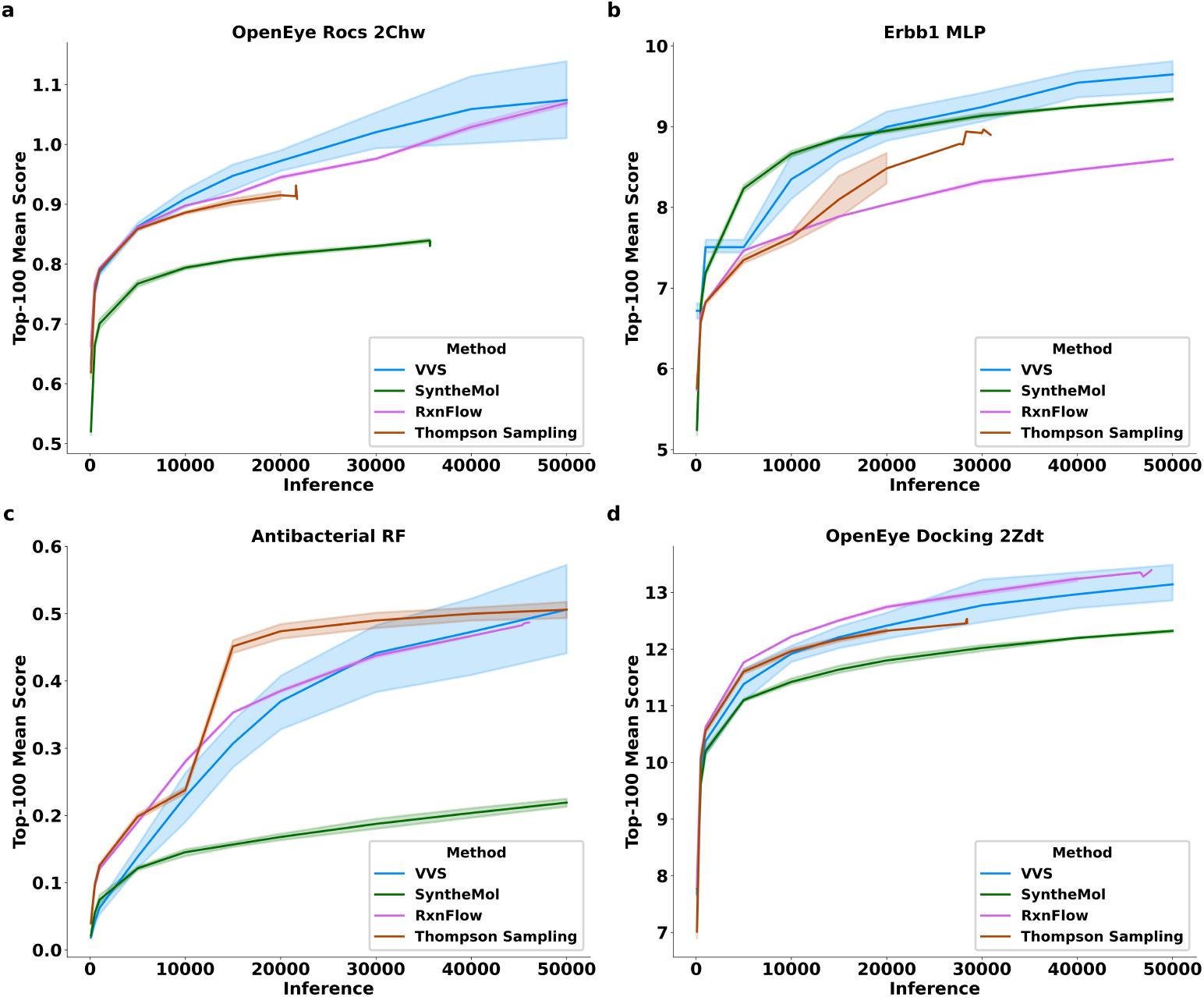
Performance of four search strategies on the Enamine Building Block dataset under a budget of 50,000 score evaluations and a 12-h wall-time limit across five independent runs. Each panel shows the average score of the top-100 molecules for each run replica as a function of total inference budget. Score magnitudes reflect the specific scoring function used. Curves that terminate before 50,000 evaluations indicate the method hit the 12-h wall-time limit before hitting the evaluation limit.

**Supplementary Fig. 13.**
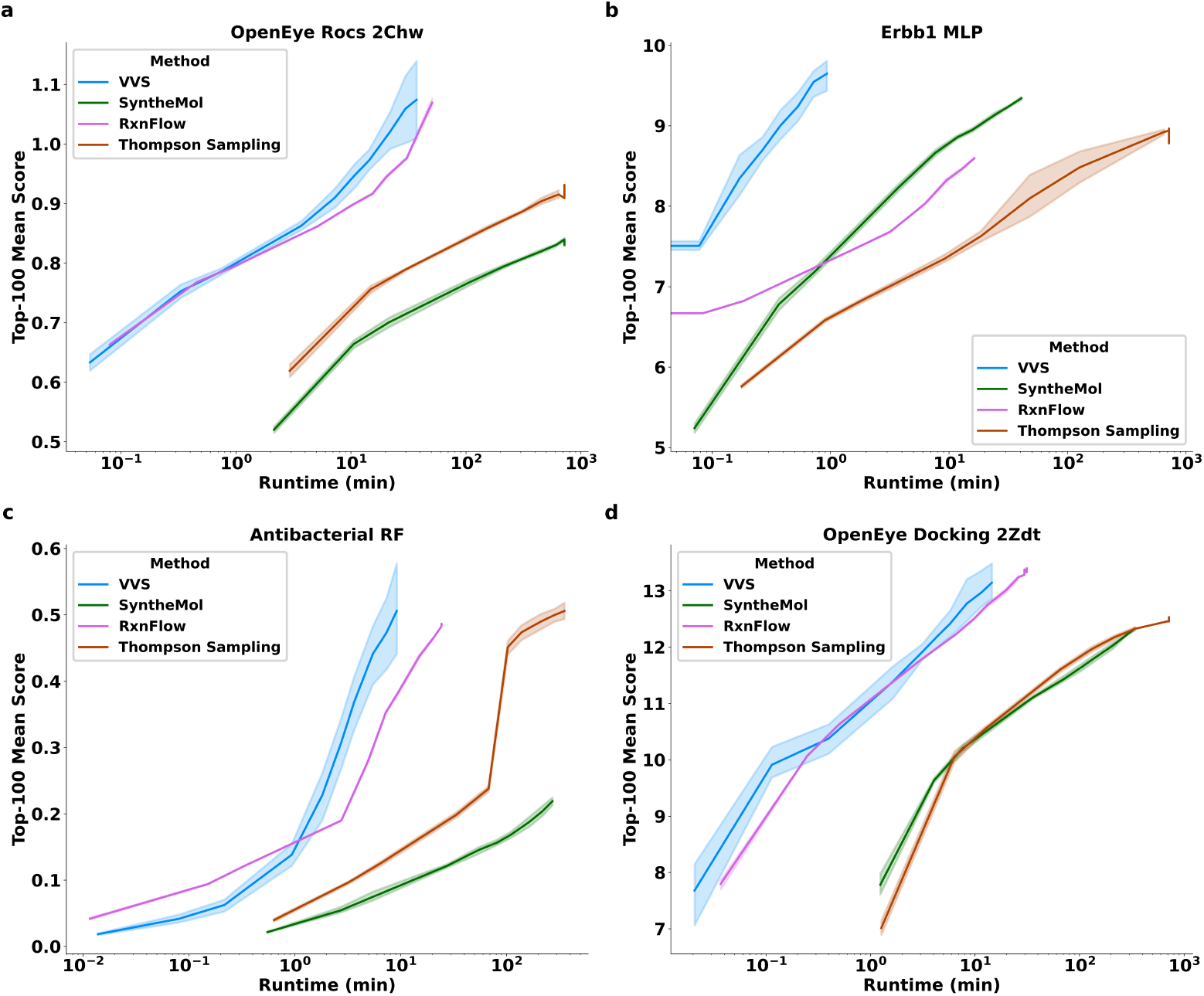
Performance of four search strategies on the Enamine Building Block dataset under a budget of 50,000 score evaluations and a 12-h wall-time limit across five independent runs. Each panel shows the average score of the top-100 molecules for each run replica as a function of runtime. Score magnitudes reflect the specific scoring function used. The starting point for each curve is the first timestamp where 100 results have been collected.

**Supplementary Fig. 14.**
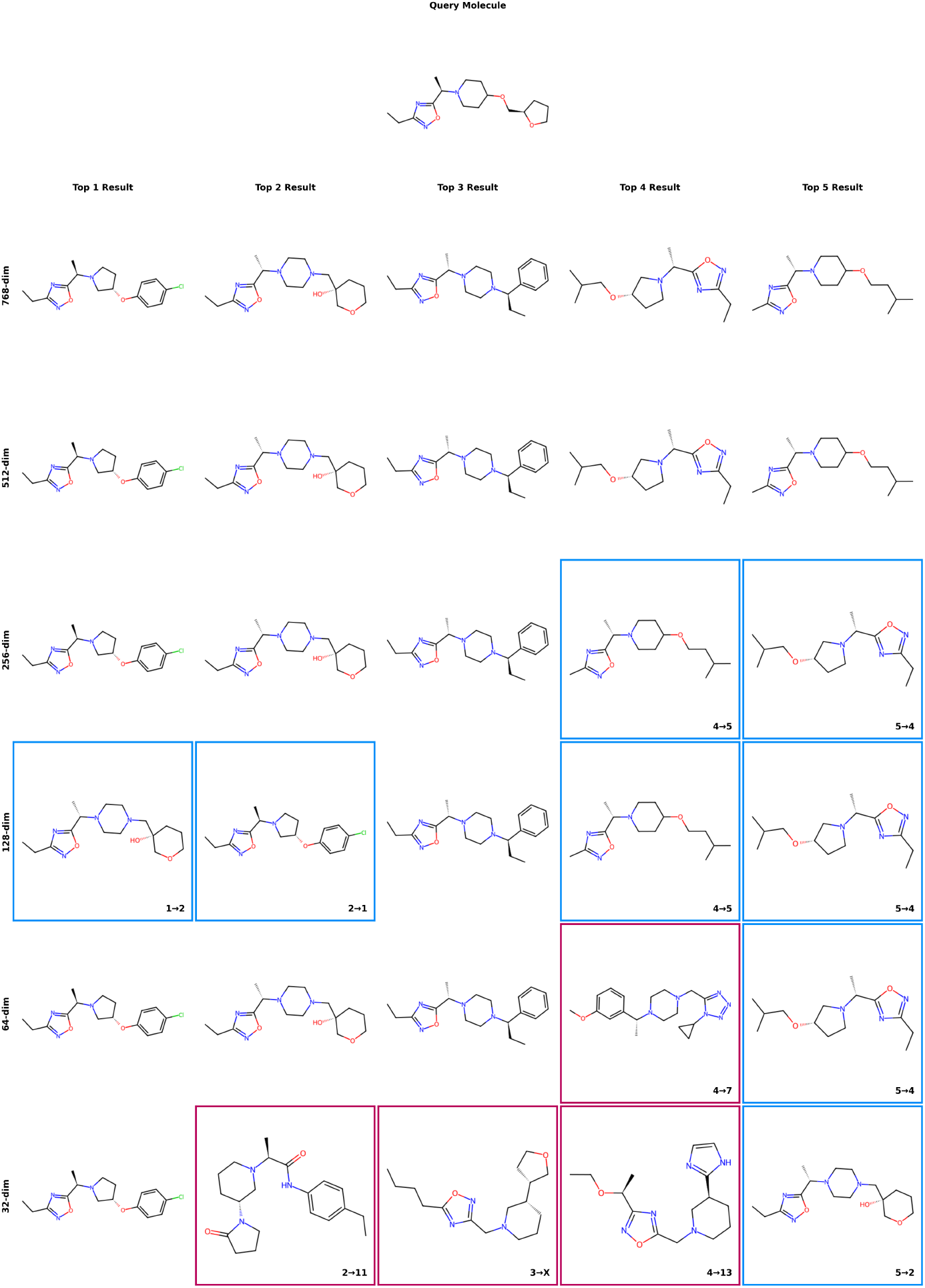
Qualitative comparison of retrieval performance for different compressed embedding sizes. **Top:** Query molecule. **Below:** Grid of retrieval results for different embedding sizes showing six rows (768 *→* 32 dims) and five columns (top-5 ranking order). The first row shows the “ground truth” top-5 nearest-neighbors retrieved from the native 768 dim embedding. Subsequent rows show the top-5 nearest-neighbors retrieved by compressed embeddings of different sizes. Cells are colored to denote comparisons to the ground truth results. Cells with no border match the molecule entity and rank of the ground truth values. Cells bordered blue are molecules from the ground truth top-5 molecules retrieved in the wrong order, with the text in the lower right of the cell denoting the compressed rank relative to the ground truth rank. Cells bordered red are molecules not found in the top-5 ground truth results, with text in the lower right cell denoting the compressed rank relative to the ground truth rank if the retrieval *k* value is extended to 20. For this example, we see perfect performance for the 512 dim compressed embedding. The 128 and 256 dim embeddings retrieve the correct molecules in the wrong order. The 64 and 32 dim embeddings retrieve some incorrect molecules, with the 32 dim top-3 result falling outside of the ground truth top-20 results (denoted by 3 *→ X*).

**Supplementary Fig. 15.**
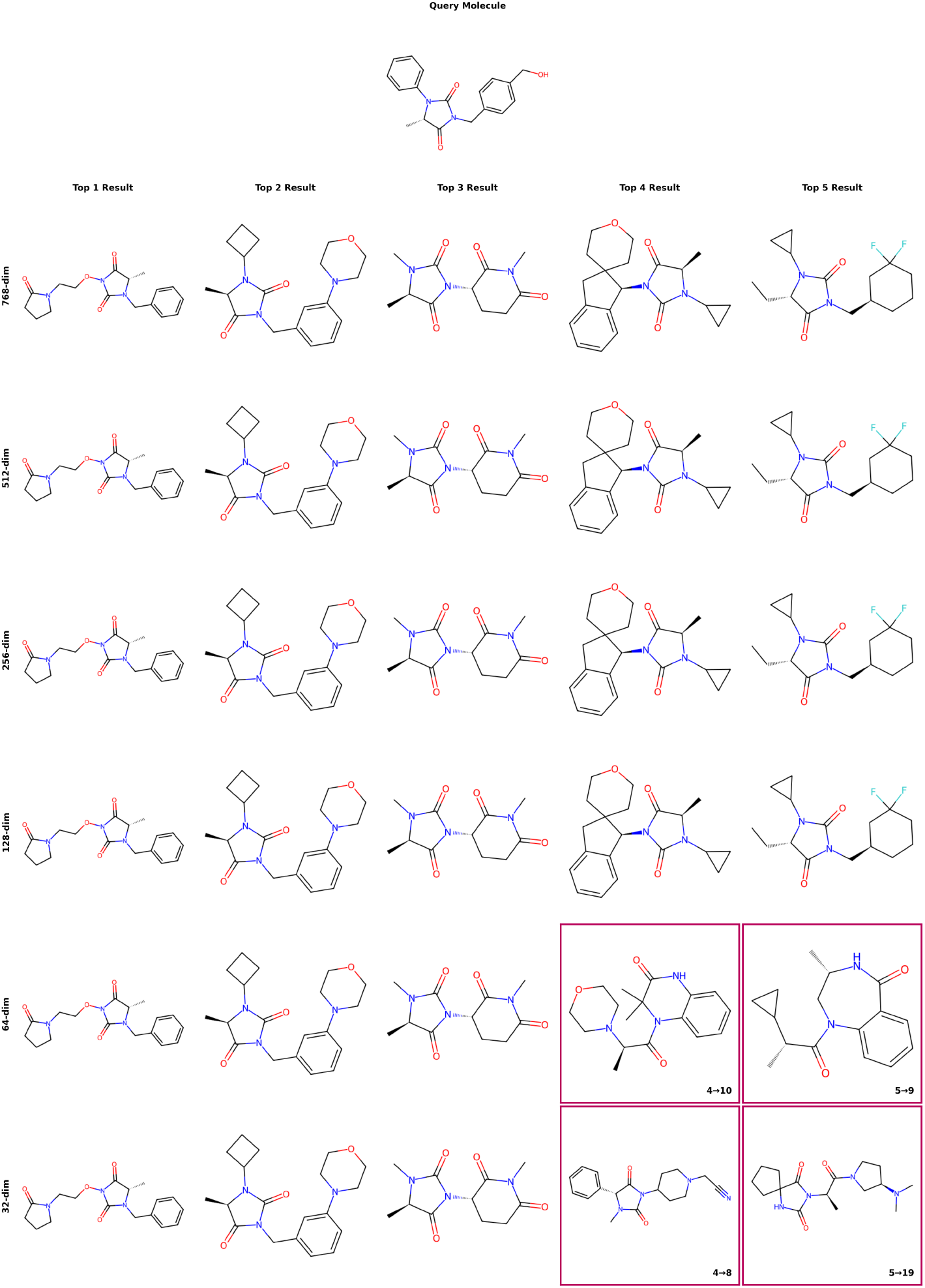
Qualitative comparison of retrieval performance for different compressed embedding sizes. **Top:** Query molecule. **Below:** Grid of retrieval results for different embedding sizes showing six rows (768 *→* 32 dims) and five columns (top-5 ranking order). The first row shows the “ground truth” top-5 nearest-neighbors retrieved from the native 768 dim embedding. Subsequent rows show the top-5 nearest neighbors retrieved by compressed embeddings of different sizes. Cells are colored to denote comparisons to the ground truth results. Cells with no border match the molecule entity and rank of the ground truth values. Cells bordered blue are molecules from the ground truth top-5 molecules re^3^tr^9^ieved in the wrong order, with the text in the lower right of the cell denoting the compressed rank relative to the ground truth rank. Cells bordered red are molecules not found in the top-5 ground truth results, with text in the lower right cell denoting the compressed rank relative to the ground truth rank if the retrieval *k* value is extended beyond 5. For this example, we see perfect performance for the 128, 256, and 512 dim embeddings, while the 32 and 64 dim embeddings retrieve molecules outside of the top 5.

**Supplementary Fig. 16.**
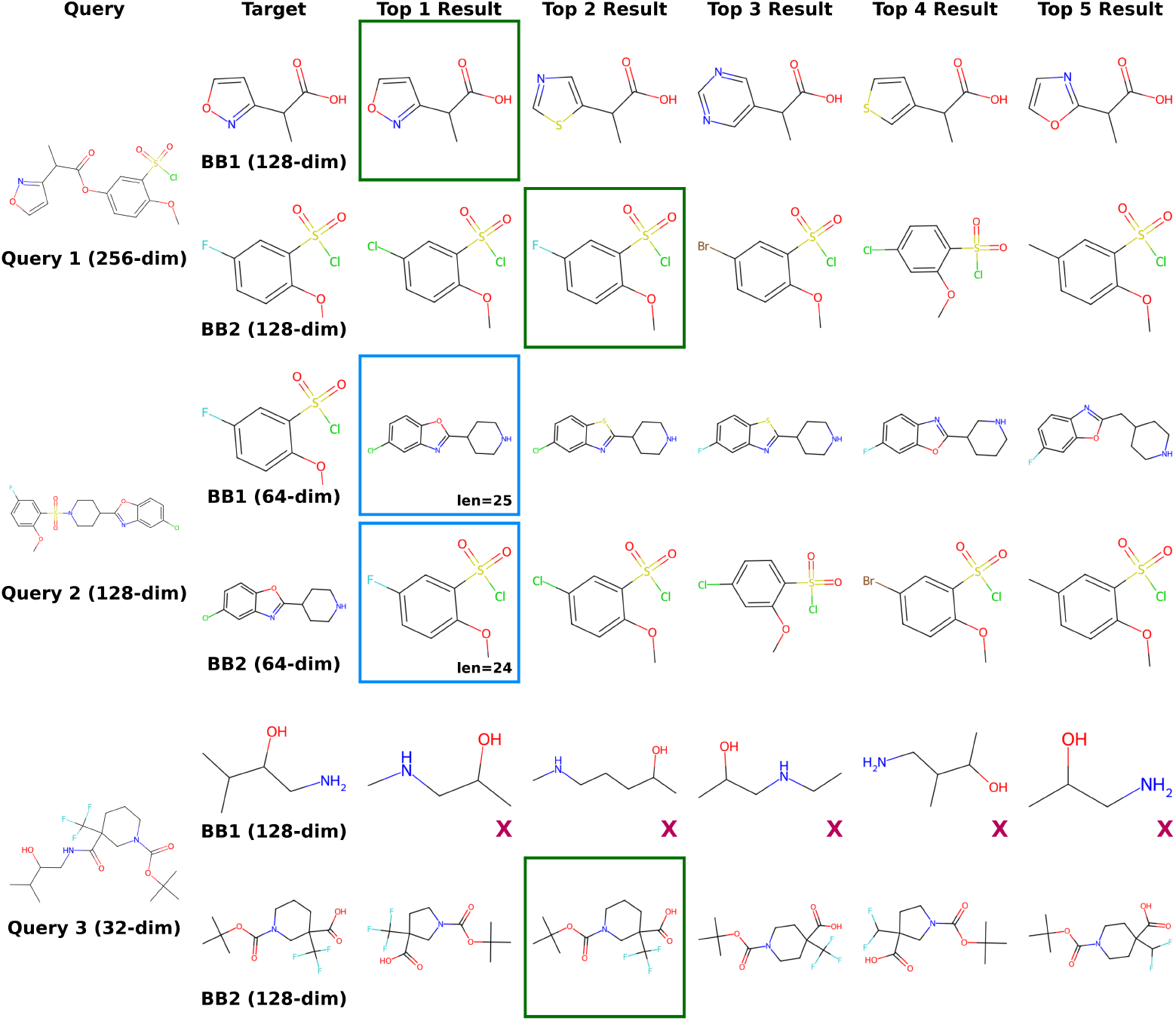
Qualitative comparison of Embedding Decomposer performance. **Query Column:** The query product molecule, annotated with the input embedding dimension. **Target Column:** The ground truth building blocks for decomposing the product molecule, annotated with the target embedding dimension. **top-***k* **Columns:** The *k* = 5 nearest-neighbors retrieved by the Embedding Decomposer model. Cells bordered in green match the target building blocks. Cells bordered in blue denote a flipped prediction, where the correct building blocks are found in the wrong order. These cells are annotated with building block SMILES string length for comparison. Cells with a red X denote that the target building block was not found within the *k* = 5 nearest-neighbor results.

**Supplementary Fig. 17.**
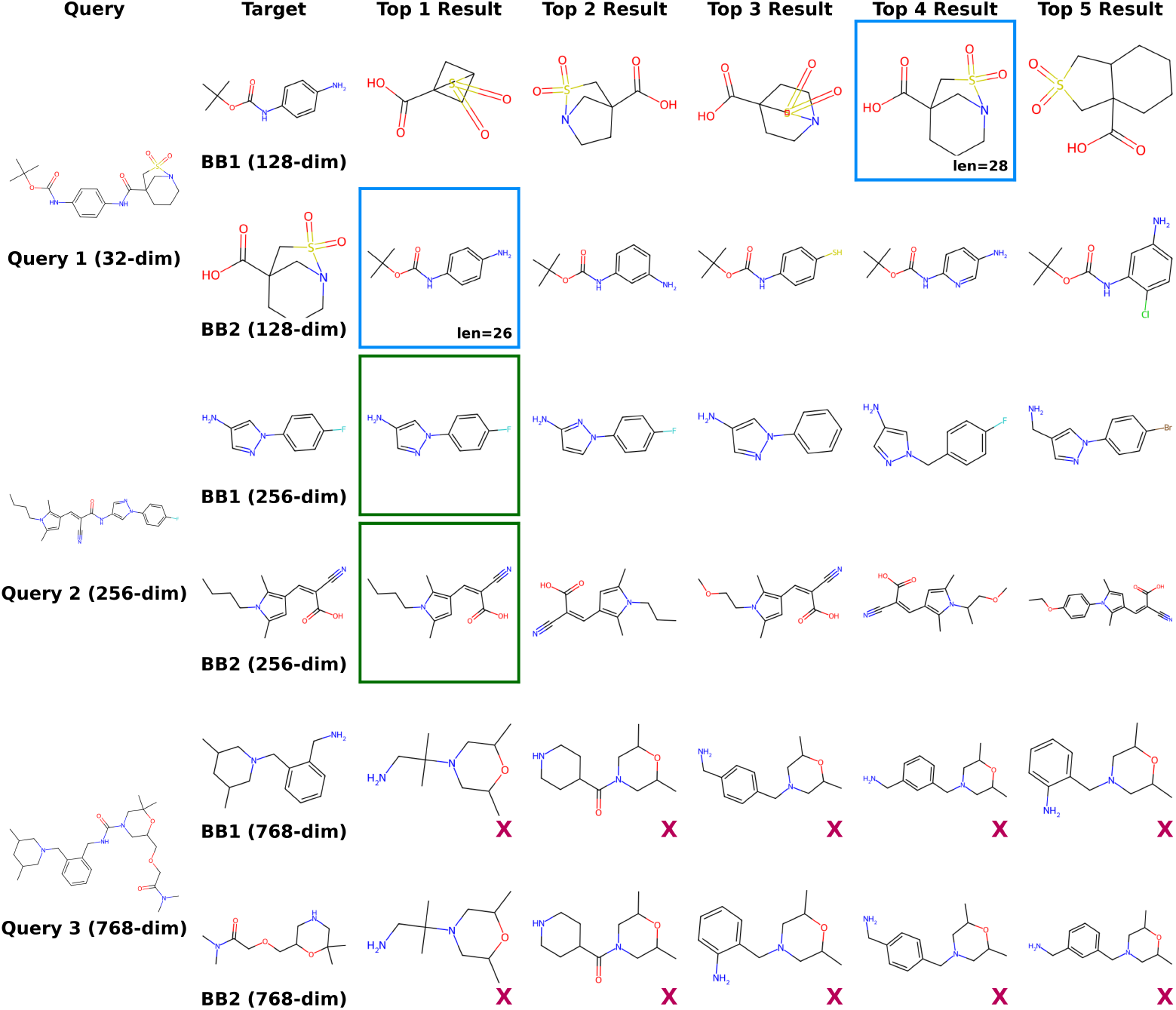
Qualitative comparison of Embedding Decomposer performance. **Query Column:** The query product molecule, annotated with the input embedding dimension. **Target Column:** The ground truth building blocks for decomposing the product molecule, annotated with the target embedding dimension. **top-***k* **Columns:** The *k* = 5 nearest-neighbors retrieved by the Embedding Decomposer model. Cells bordered in green match the target building blocks. Cells bordered in blue denote a flipped prediction, where the correct building blocks are found in the wrong order. These cells are annotated with building block SMILES string length for comparison. Cells with a red X denote that the target building block was not found within the *k* = 5 nearest-neighbor results.

**Supplementary Fig. 18.**
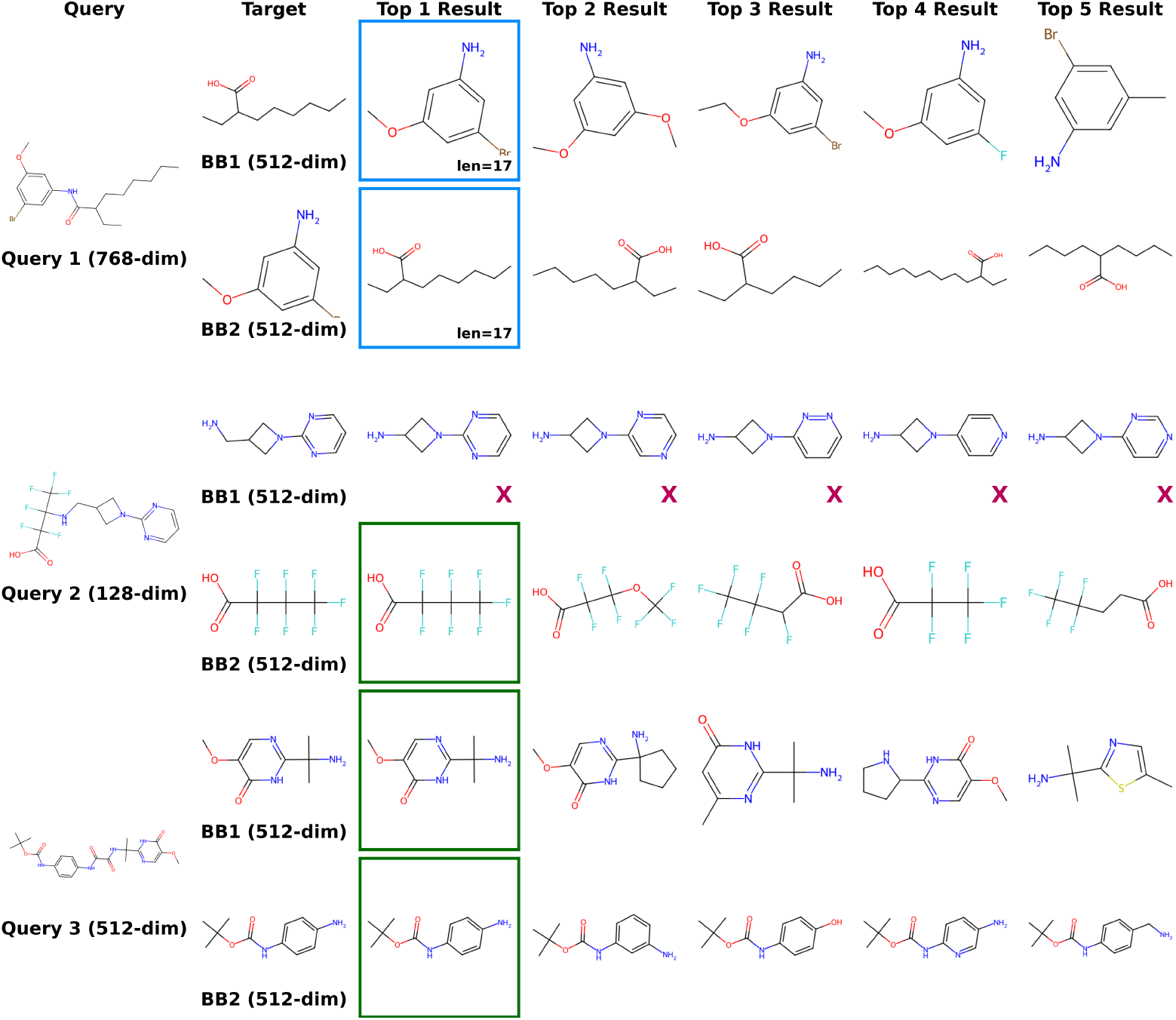
Qualitative comparison of Embedding Decomposer performance. **Query Column:** The query product molecule, annotated with the input embedding dimension. **Target Column:** The ground truth building blocks for decomposing the product molecule, annotated with the target embedding dimension. **top-***k* **Columns:** The *k* = 5 nearest-neighbors retrieved by the Embedding Decomposer model. Cells bordered in green match the target building blocks. Cells bordered in blue denote a flipped prediction, where the correct building blocks are found in the wrong order. These cells are annotated with building block SMILES string length for comparison. Cells with a red X denote that the target building block was not found within the *k* = 5 nearest-neighbor results.

**Supplementary Fig. 19.**
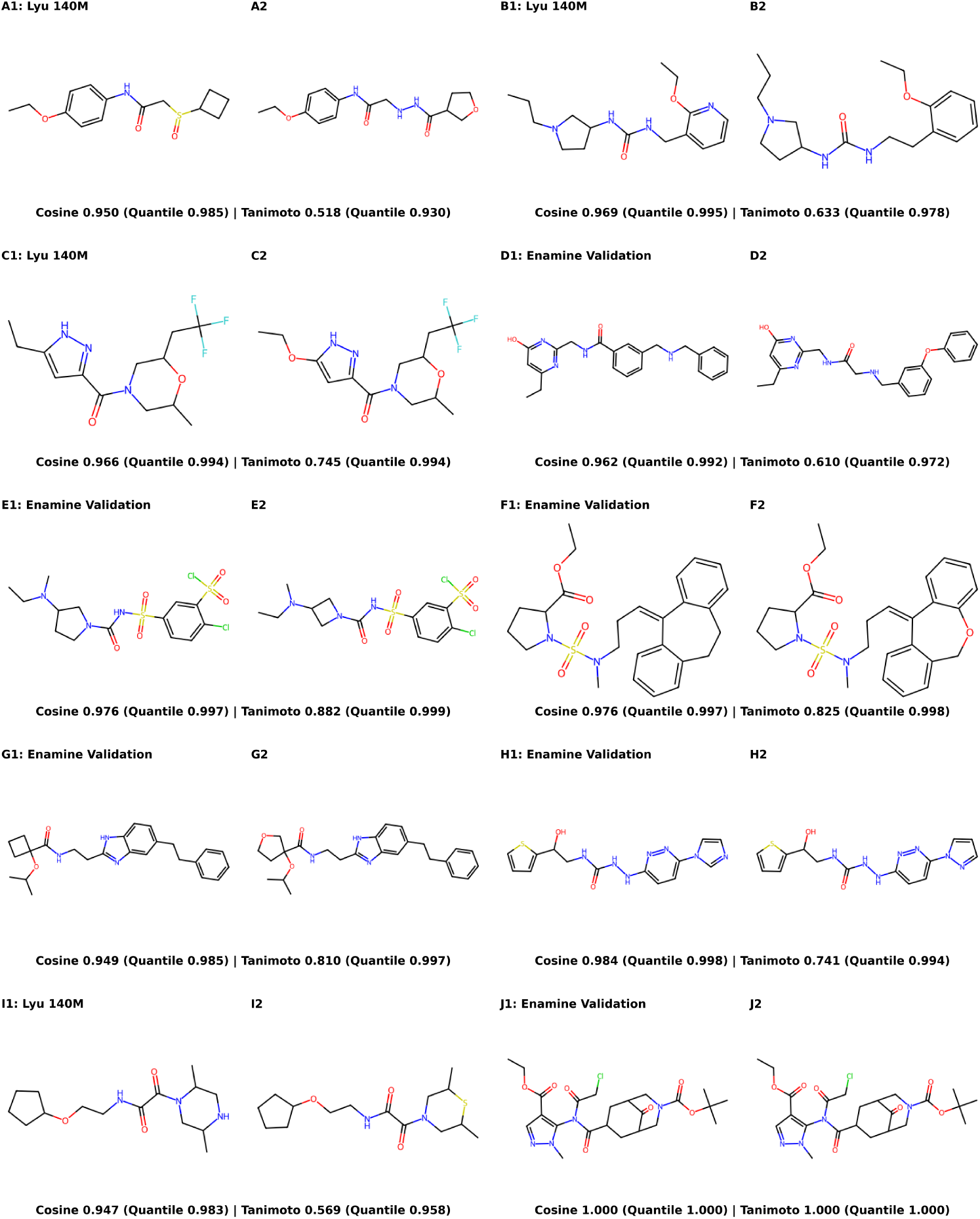
Qualitative comparison of BBKNN query and result molecules for queries from the Enamine Assembled dataset (“Enamine Validation”) and Lyu 140M dataset. Each query-result pair is annotated with their cosine and Tanimoto (ECFP-4) similarity. Each similarity metric has a quantile value denoting the ranking of the value relative to a dataset of 2.3M molecule pairs.

**Supplementary Fig. 20.**
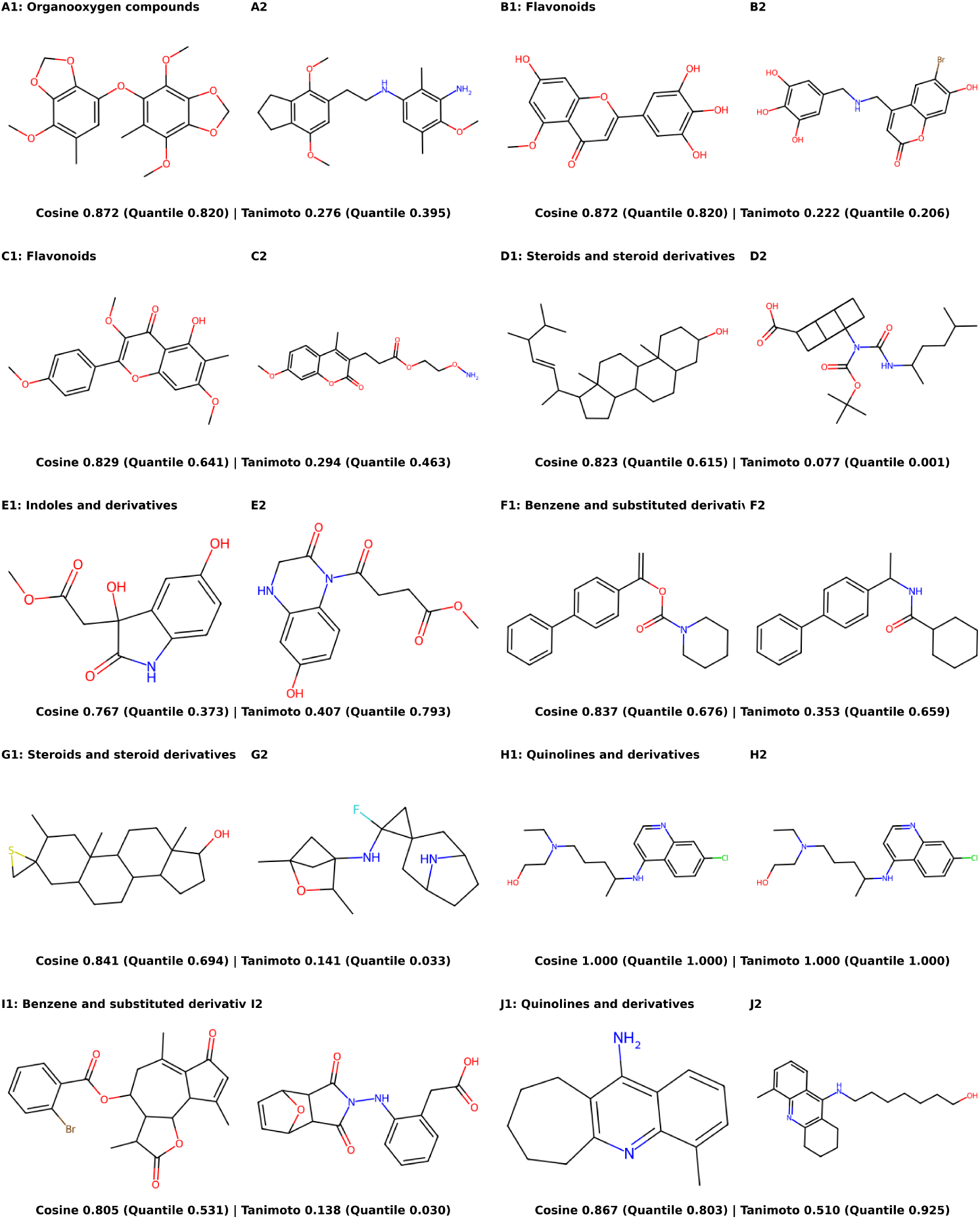
Qualitative comparison of BBKNN query and result molecules for natural product queries from the COCONUT 2.0 dataset. Each query molecule is annotated with its natural product class. Each query-result pair is annotated with their cosine and Tanimoto (ECFP-4) similarity and the quantile of each metric relative to a dataset of 2.3M molecule pairs.

**Supplementary Fig. 21.**
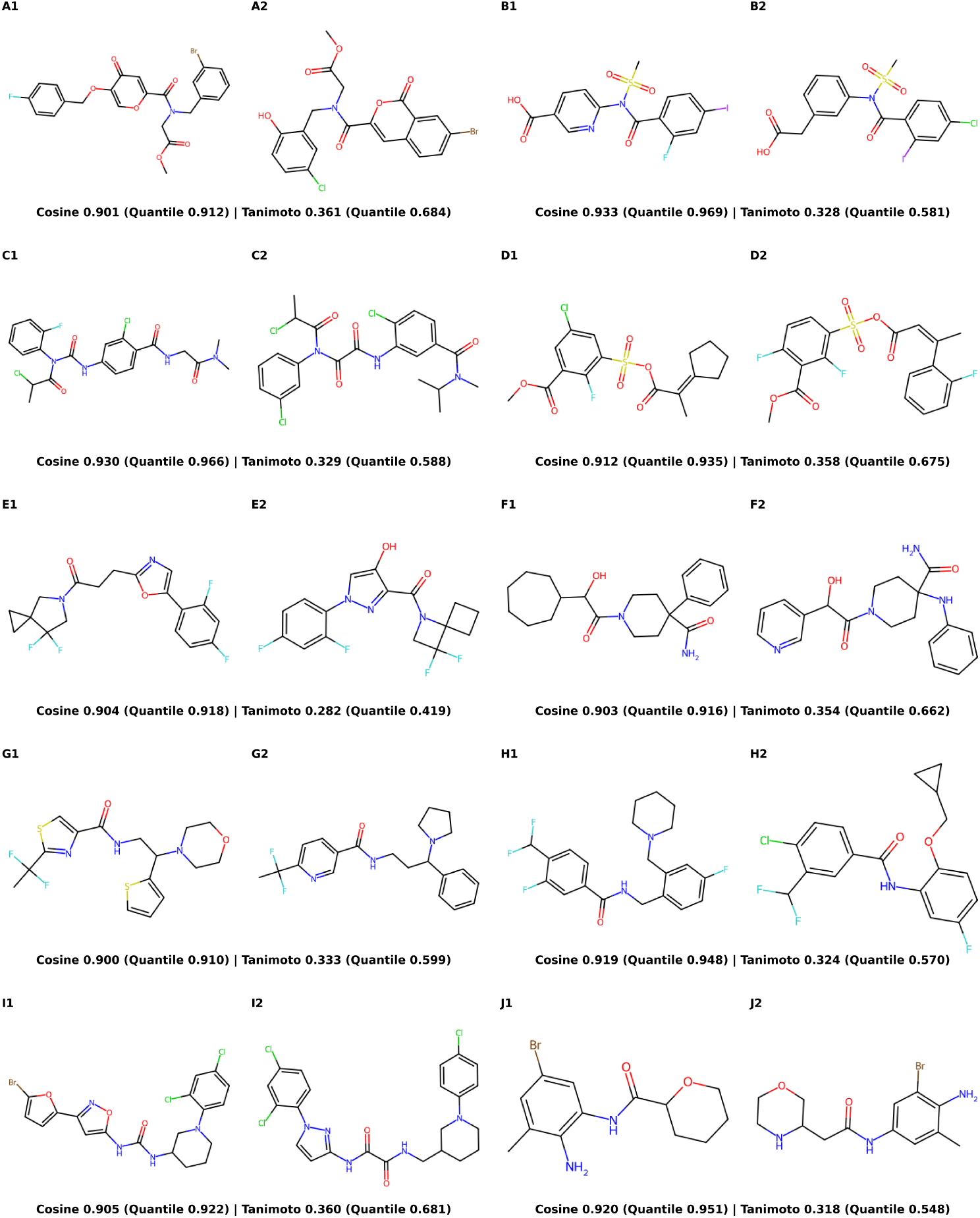
Qualitative comparison of disagreements between cosine and Tanimoto similarity showing high cosine similarity and low Tanimoto similarity. Each pair is annotated with their cosine and Tanimoto (ECFP-4) similarity and the quantile of each metric relative to a dataset of 2.3M molecule pairs.

**Supplementary Fig. 22.**
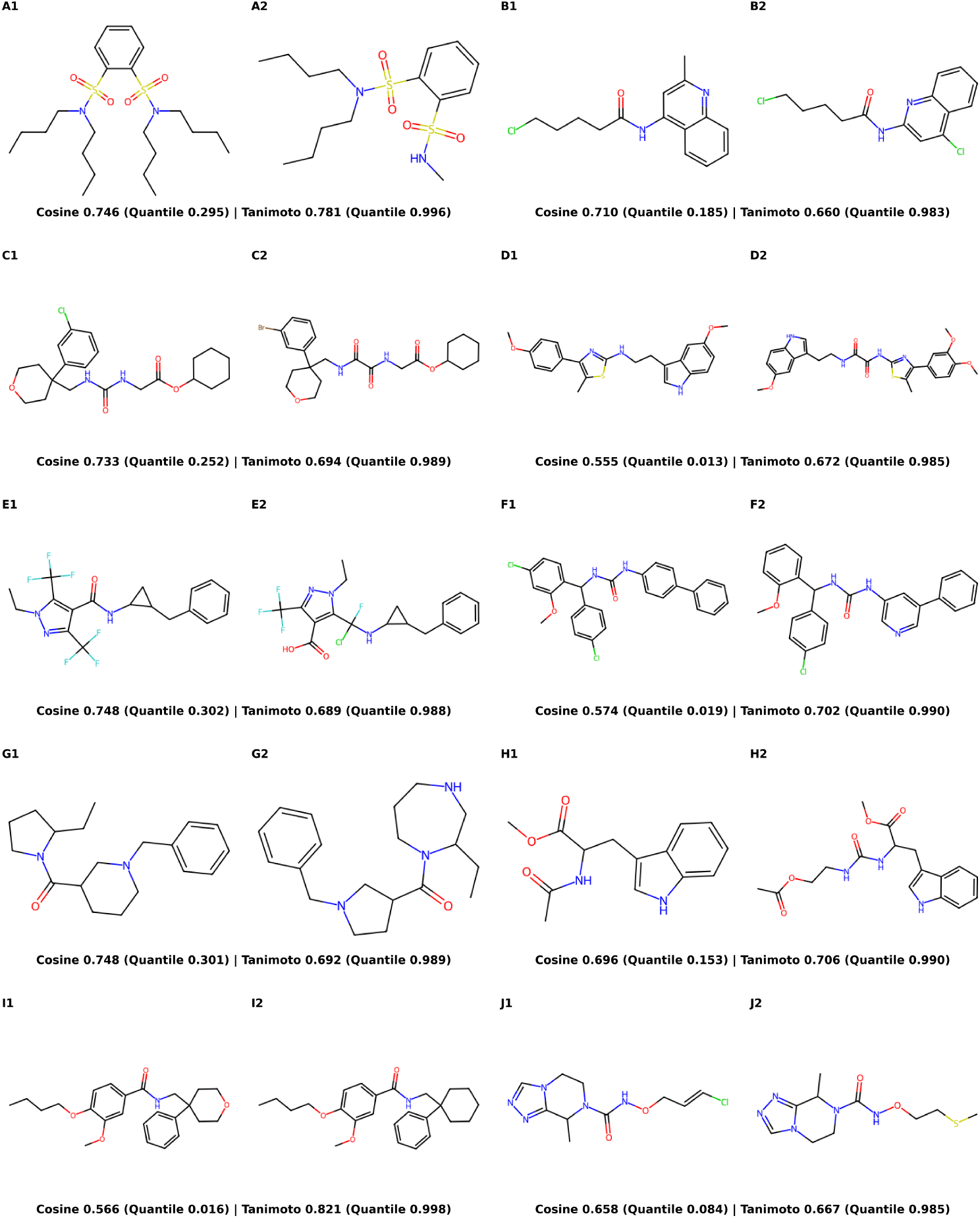
Qualitative comparison of disagreements between cosine and Tanimoto similarity showing low cosine similarity and high Tanimoto similarity. Each pair is annotated with their cosine and Tanimoto (ECFP-4) similarity and the quantile of each metric relative to a dataset of 2.3M molecule pairs.

**Supplementary Fig. 23.**
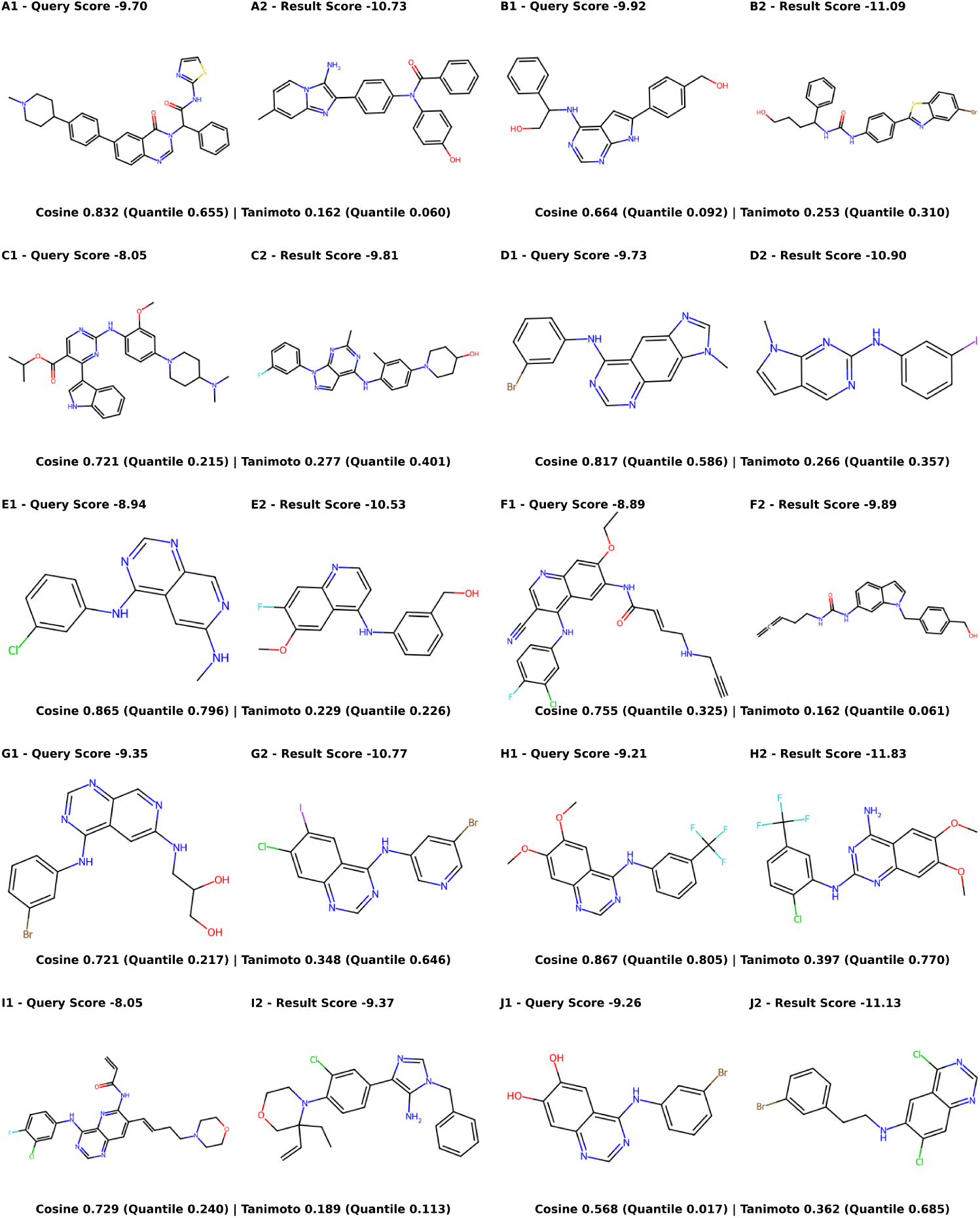
EGFR analogs retrieved using BBKNN to query 50 known EGFR ligands from the Chembl ErbB1 against the Enamine Building Block search space. Each pair of results represents a query molecule (known EGFR ligand) and a retrieved result. Each molecule is annotated with its docking score against EGFR. Each pair is annotated with their cosine and Tanimoto (ECFP-4) similarity and the quantile of each metric relative to a dataset of 2.3M molecule pairs to contextualize the value.

### 2 Supplementary Tables

**Supplementary Table 1.** Parameter count for different embedding compression models. Output size denotes the size that the input embedding (768-dim) is compressed to. The ‘Encoder parameters’ column reflects the number of parameters in the encoder used for the compression. The ‘Total parameters’ column includes the decoder parameter count, which is only used for training.

| Output Size | Encoder Parameters | Total Parameters |
| --- | --- | --- |
| 512 | 6,763,776 | 13,856,256 |
| 256 | 5,976,320 | 12,478,976 |
| 128 | 5,631,744 | 11,839,488 |
| 64 | 5,471,744 | 11,532,032 |
| 32 | 5,394,816 | 11,381,376 |

**Supplementary Table 2.**
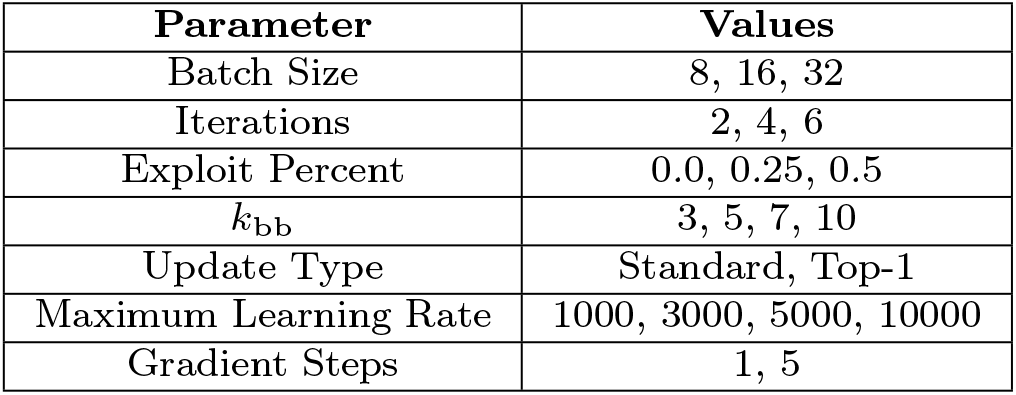
Parameter ranges explored in the VVS hyperparameter sweep.

**Supplementary Table 3.**
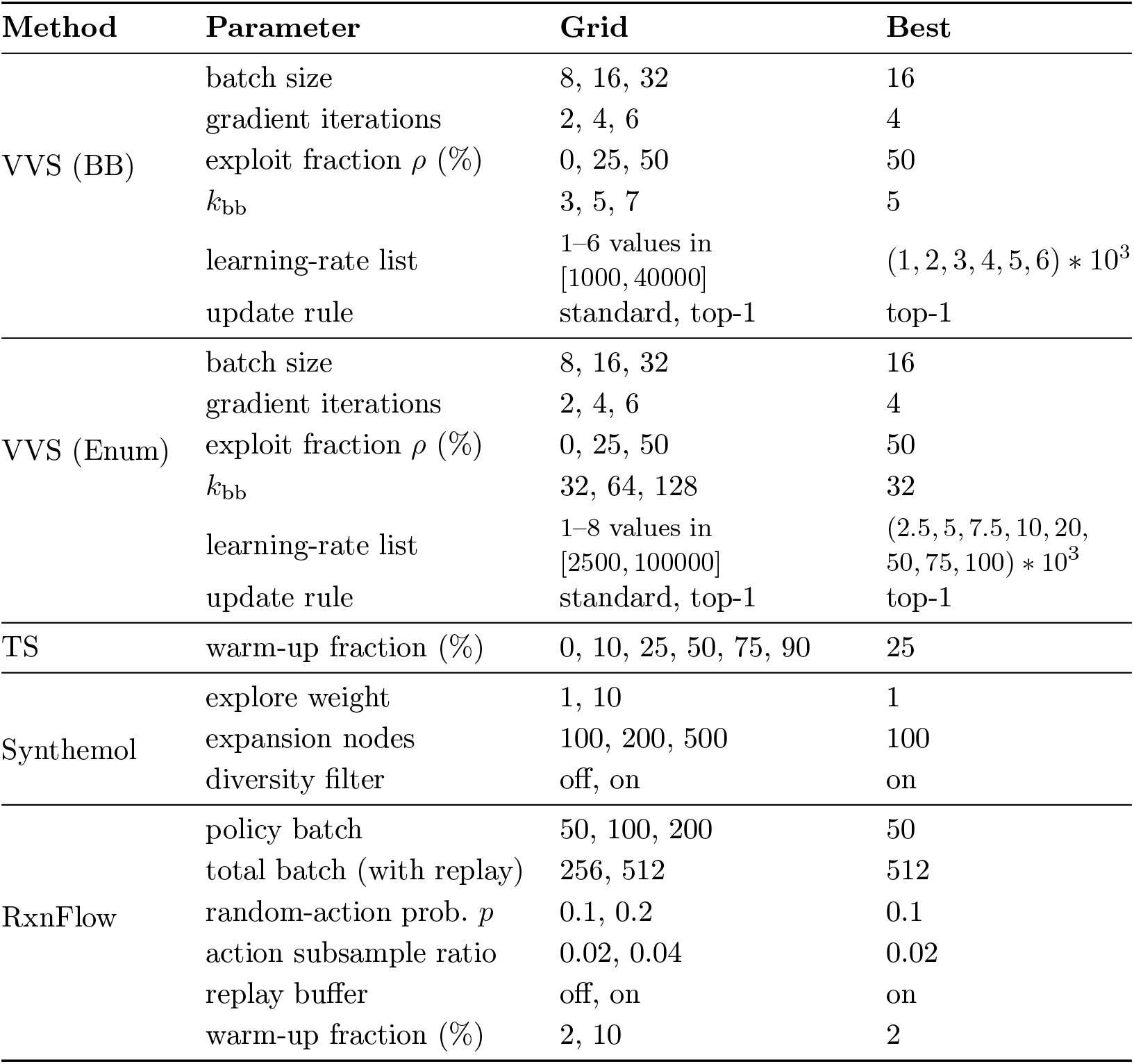
Hyperparameter grids tested for different search methods against the EGFR pIC_50_ MLP scoring function and the best value for each parameter. Best parameters were determined by running each parameter combination in triplicate and evaluating the Top-*{*1, 10, 100*}* scoring molecules in each run.

| Method | Parameter | Grid | Best |
| --- | --- | --- | --- |
| VVS (BB) | batch size | 8, 16, 32 | 16 |
|  | gradient iterations | 2, 4, 6 | 4 |
| | exploit fraction $\rho$ (%) | 0, 25, 50 | 50 |
| | $k_{bb}$ | 3, 5, 7 | 5 |
| | learning-rate list | 1-6 values in [1000, 40000] | $(1, 2, 3, 4, 5, 6) * 10^3$ |
|  | update rule | standard, top-1 | top-1 |
| VVS (Enum) | batch size | 8, 16, 32 | 16 |
|  | gradient iterations | 2, 4, 6 | 4 |
| | exploit fraction $\rho$ (%) | 0, 25, 50 | 50 |
| | $k_{bb}$ | 32, 64, 128 | 32 |
| | learning-rate list | 1-8 values in [2500, 100000] | $(2.5, 5, 7.5, 10, 20, 50, 75, 100) * 10^3$ |
|  | update rule | standard, top-1 | top-1 |
| TS | warm-up fraction (%) | 0, 10, 25, 50, 75, 90 | 25 |
| Synthemol | explore weight | 1, 10 | 1 |
|  | expansion nodes | 100, 200, 500 | 100 |
|  | diversity filter | off, on | on |
| RxnFlow | policy batch | 50, 100, 200 | 50 |
|  | total batch (with replay) | 256, 512 | 512 |
| | random-action prob. $p$ | 0.1, 0.2 | 0.1 |
|  | action subsample ratio | 0.02, 0.04 | 0.02 |
|  | replay buffer | off, on | on |
|  | warm-up fraction (%) | 2, 10 | 2 |

**Supplementary Table 4.**
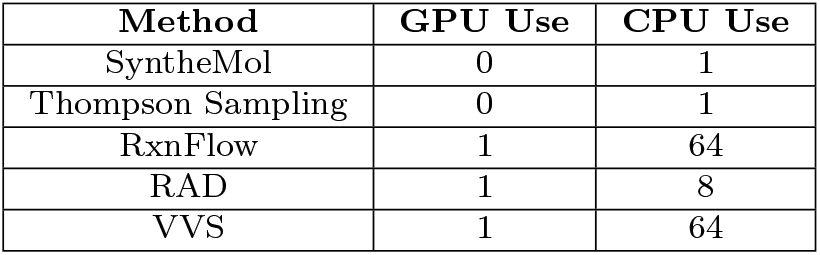
Benchmark compute usage for different methods. GPU (A100) and CPU usage was determined by the author’s code implementation. See Supp. Notes 4.9 for implementation details.

| Method | GPU Use | CPU Use |
| --- | --- | --- |
| SyntheMol | 0 | 1 |
| Thompson Sampling | 0 | 1 |
| RxnFlow | 1 | 64 |
| RAD | 1 | 8 |
| VVS | 1 | 64 |

**Supplementary Table 5.** Performance of four search strategies on the Enamine Building Block dataset under a budget of 50,000 score evaluations and a 12-h wall-time limit. Each cell shows the mean ± s.d. of five independent runs. The “Top- *{*1, 10, 100*}*” columns report the average score of the best k molecules found in each run (higher = better ↑). Score magnitudes reflect the specific scoring function used. “Runtime” is the wall-clock time in minutes until the evaluation limit is reached (lower = better ↓); a value of 720^*\**^ minutes indicates the method exhausted the 12-hour budget before completing all 50k evaluations.

| Score | Method | Top-1 ↑ | Top-10 ↑ | Top-100 ↑ | Runtime ↓ |
| --- | --- | --- | --- | --- | --- |
| Docking | RxnFlow | 15.36 ± 0.73 | 14.33 ± 0.16 | 13.35 ± 0.05 | 30.6 ± 0.8 |
| | SyntheMol | 13.83 ± 0.48 | 13.30 ± 0.12 | 12.32 ± 0.03 | 336 ± 0.5 |
| | TS | 14.75 ± 0.67 | 13.67 ± 0.17 | 12.50 ± 0.04 | 720* |
| | VVS | 15.01 ± 0.70 | 14.18 ± 0.55 | 13.14 ± 0.40 | 14.5 ± 1.4 |
| EGFR pIC<sub>50</sub> MLP | RxnFlow | 9.86 ± 0.09 | 9.35 ± 0.05 | 8.60 ± 0.01 | 16.4 ± 1.0 |
| | SyntheMol | 10.04 ± 0.15 | 9.85 ± 0.09 | 9.34 ± 0.03 | 40.8 ± 0.0 |
| | TS | 9.55 ± 0.19 | 9.22 ± 0.18 | 8.87 ± 0.09 | 720* |
| | VVS | 10.21 ± 0.20 | 10.02 ± 0.21 | 9.65 ± 0.25 | 0.93 ± 0.0 |
| ROCS | RxnFlow | 1.26 ± 0.03 | 1.19 ± 0.02 | 1.06 ± 0.01 | 50.9 ± 0.0 |
| | SyntheMol | 0.97 ± 0.01 | 0.92 ± 0.01 | 0.84 ± 0.00 | 720* |
| | TS | 1.06 ± 0.06 | 1.00 ± 0.04 | 0.92 ± 0.01 | 720* |
| | VVS | 1.20 ± 0.11 | 1.16 ± 0.11 | 1.07 ± 0.08 | 37.3 ± 0.6 |
| Anti-microbial RF | RxnFlow | 0.56 ± 0.00 | 0.53 ± 0.00 | 0.49 ± 0.00 | 24.4 ± 0.0 |
| | SyntheMol | 0.61 ± 0.15 | 0.39 ± 0.03 | 0.22 ± 0.01 | 273 ± 0.1 |
| | TS | 0.58 ± 0.03 | 0.55 ± 0.03 | 0.51 ± 0.02 | 354 ± 1.7 |
| | VVS | 0.63 ± 0.13 | 0.60 ± 0.13 | 0.51 ± 0.09 | 9.22 ± 0.0 |

**Supplementary Table 6.** Performance of VVS and RAD search strategies on the Lyu 140M under a budget of 50,000 score evaluations and a 12-h wall-time limit. Each cell shows the mean ± s.d. of five independent runs. The “Top- *{*1, 10, 100*}* columns report the average score of the best *k* molecules found in each run (higher = better ↑). Score magnitudes reflect the specific scoring function used. “Runtime” is the wall-clock time in minutes until the evaluation limit is reached (lower = better ↓).

| Score | Method | Top-1 ↑ | Top-10 ↑ | Top-100 ↑ | Runtime ↓ |
| --- | --- | --- | --- | --- | --- |
| Docking | RAD | 16.00 ± 0.00 | 15.71 ± 0.00 | 15.05 ± 0.00 | 43.50 ± 1.23 |
| | VVS | 15.66 ± 0.21 | 14.90 ± 0.08 | 13.96 ± 0.09 | 19.67 ± 0.57 |
| EGFR pIC<sub>50</sub> MLP | RAD | 10.26 ± 0.00 | 9.87 ± 0.00 | 9.40 ± 0.00 | 6.34 ± 0.04 |
| | VVS | 10.18 ± 0.09 | 9.97 ± 0.04 | 9.67 ± 0.13 | 0.47 ± 0.02 |
| ROCS | RAD | 1.30 ± 0.00 | 1.28 ± 0.00 | 1.20 ± 0.00 | 121.65 ± 0.11 |
| | VVS | 1.21 ± 0.04 | 1.17 ± 0.03 | 1.08 ± 0.02 | 39.42 ± 0.70 |
| Anti-microbial RF | RAD | 0.97 ± 0.00 | 0.96 ± 0.00 | 0.70 ± 0.00 | 37.63 ± 0.03 |
| | VVS | 0.82 ± 0.00 | 0.73 ± 0.00 | 0.55 ± 0.00 | 8.83 ± 0.05 |

### 3 Supplementary Algorithms

#### Algorithm 1

BBKNN: Building-Block *K*-Nearest-Neighbors

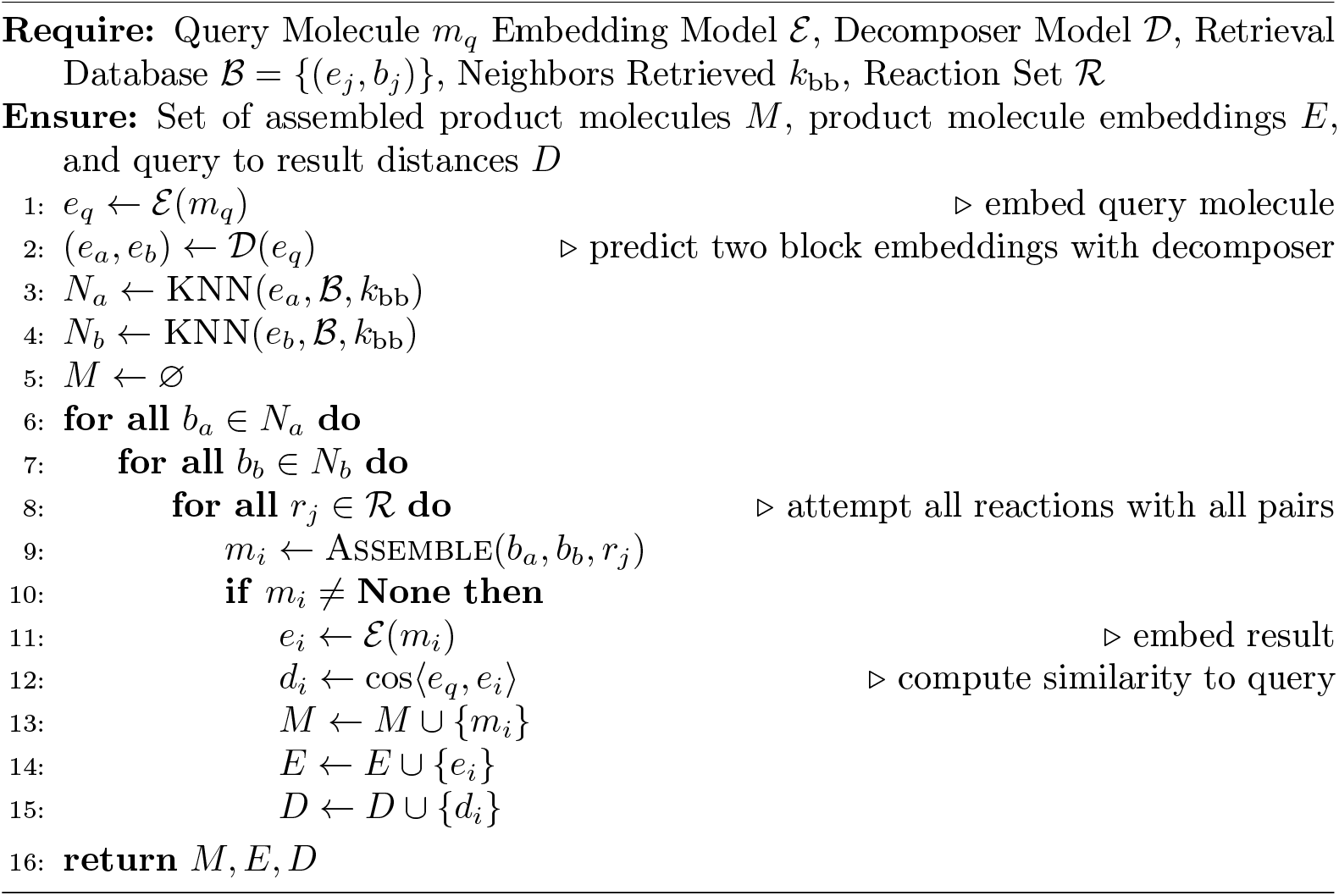

#### Algorithm 2

Gradient Estimation

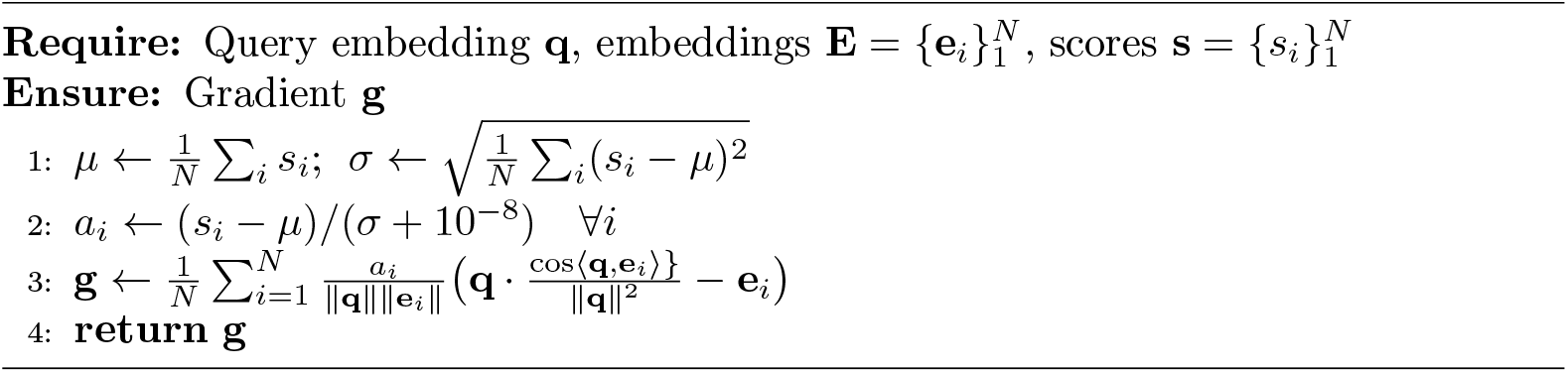

#### Algorithm 3

Single VVS Step

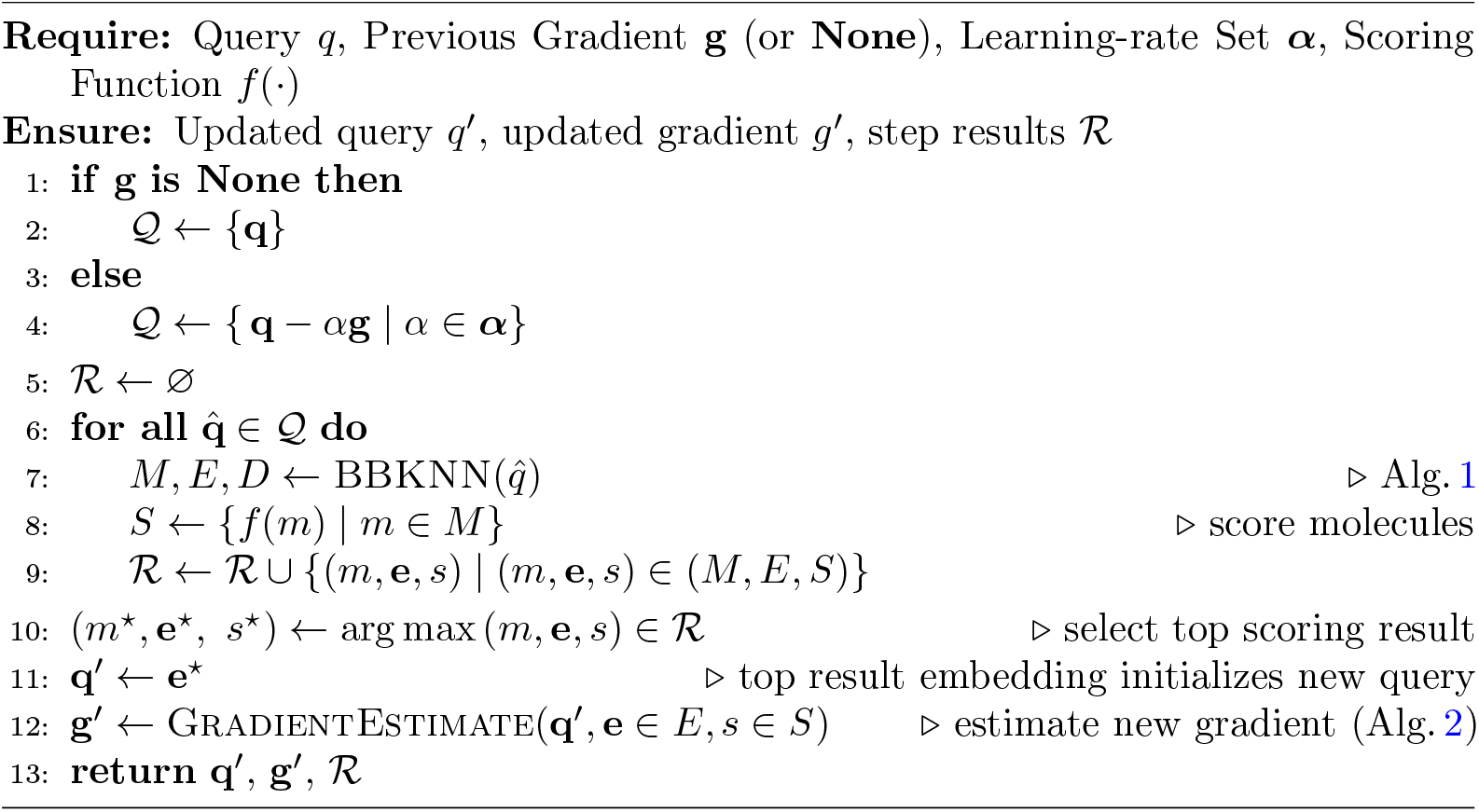

#### Algorithm 4

Batched VVS Outer Loop with Exploitation

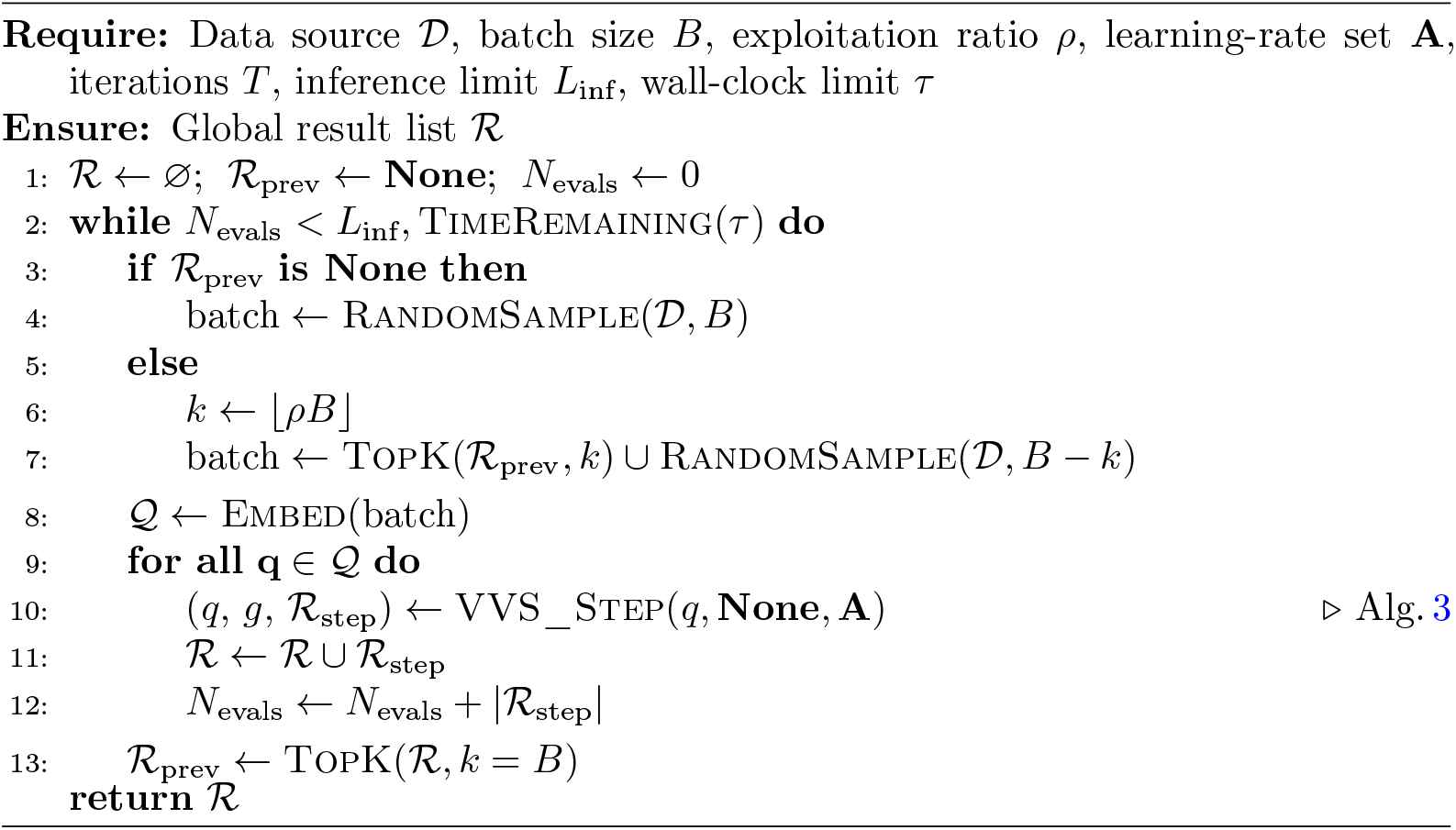

## 4 Supplementary Notes

### 4.1 Related Works

To fully realize the promise of these ultra-large building block libraries, there is a pressing need for new computational strategies that can efficiently navigate and prioritize within these vast combinatorial spaces without requiring library enumeration and exhaustive screening. Current approaches to searching and screening these libraries fall into several broad classes:

#### Analogs by Similarity

Similarity-based search in combinatorial libraries enables the retrieval of chemically similar analogs of query molecules (i.e., the nearest-neighbors that can actually be purchased or commissioned from a fixed vendor catalog) without enumerating the entire product space [7, 10–14]. One shortcoming of traditional retrosynthesis objectives, which aim to predict the reactant molecules of a query product molecule, is the inherent uncertainty in the synthesizability of the predicted reactant molecules. Instead, we aim to retrieve the most similar synthesis-on-demand alternatives given a specific set of building blocks and assembly reaction. Traditional workflows achieve this by first applying cheminformatics heuristics to break the query molecule into fragments or synthon structures [14, 15], followed by matching those structures against available building blocks using 2D fingerprint similarity [10], feature trees [13], graph-editdistance [16] or related metrics; the top-ranked blocks are re-assembled *in silico* to yield catalog-compatible analogs. Although effective, these pipelines rely on extensive rule bases and deep retrosynthetic expertise, so implementations remain largely proprietary and inaccessible to the research community.

Recent machine-learning advances offer a complementary route for similarity search: dense embeddings from chemical language models (CLMs) capture subtle structural and physicochemical relationships, enabling fast nearest-neighbor retrieval via simple vector operations such as cosine similarity [17, 18]. Coupled with modern vector-database indexes, this makes sub-second analog search feasible even in billion-scale embedding spaces [19]. However, existing embedding-based approaches require a fully enumerated molecule library, a requirement that quickly becomes intractable for the 10^10^–10^12^ products implied by typical combinatorial catalogs.

In parallel, several machine learning approaches have been developed for general retrosynthesis [20–24]. However, these methods generally focus on the total synthesis of specific molecules as opposed to finding analogs available in a specific building block catalog.

#### Fragment Methods

Fragment-based methods manage the scale of combinatorial libraries by first evaluating individual building blocks in the context of a protein pocket, then assembling molecules from high scoring fragments [25–27]. Some machine learning variants extend this concept by “growing” a ligand within a protein pocket one fragment at a time, using the docking score of the growing ligand as a reward signal [28–31]. This class of methods avoids the scale explosion of combinatorial libraries by working in building block space, but is restricted to only using structure-based scoring. Furthermore, these methods rely on the assumption that fragment scores are indicative of full molecule scores, which is often an invalid assumption [32].

#### Active Learning

Active Learning (AL) [33] methods attempt to manage the scale of chemical spaces by fitting a fast surrogate model to an expensive scoring function (e.g. docking) and iteratively refining it. The surrogate model predicts the scores for the search library, selects the most promising compounds to score with the expensive score, re-trains the surrogate on the results, and then repeats. Although this approach has been applied successfully to virtual screening [34–38], it requires the search space to be fully enumerated. As the size of the search space grows, the computational overhead induced by multiple iterations of scoring the search space with the surrogate function becomes prohibitive, even with an extremely cheap surrogate function. Although there are methods to reduce the inference burden of active learning [39, 40], it is not sufficient to overcome the scale of modern building block libraries.

### Datasets and Chemical Libraries

We utilize several chemical datasets to train models and evaluate our algorithms. Specific use cases of the different datasets are discussed in the relevant methods sections. For all datasets, SMILES strings were canonicalized with RDKit [56].

#### Enamine Building Block

The Enamine Building Block dataset consists of 130,000 building blocks and 12 two-component SMARTS assembly reactions from the Enamine REAL space [5] (Enamine reactions 11, 22, 27, 40, 527, 1458, 2230, 2430, 2708, 2718, 240690, and 271948) sourced from Swanson et al. [44]. The building block dataset represents a combinatorial space of approximately 10^10^ synthesizable molecules when theoretically enumerated. All reactions use two building blocks, but may also utilize other reagents or components that add additional matter.

#### Lyu 140M

The Lyu 140M dataset contains approximately 1.4 *×* 10^8^ synthesis-on-demand compounds generated from 130 reactions and 70,000 Enamine building blocks and 130 assembly reactions sourced from Lyu et al. [42]. This dataset consists of only product molecules, so the exact overlap between the building blocks and reactions used to create the Lyu 140M dataset with the building blocks and reactions in the Enamine Building Block dataset is unknown.

#### Zinc 10M

The Zinc 10M dataset contains 10M molecules from the ZINC-22 library [51] provided by the DeepChem [57] python library.

#### COCONUT 2.0

The COCONUT 2.0 dataset consists of 700,000 natural products spanning several molecule classes sourced from the COCONUT natural product database [43].

#### Chembl ErbB1

The Chembl ErbB1 dataset consists of 7,000 molecules with measured binding affinities (IC_50_) against the epidermal growth factor receptor (EGFR) downloaded from the ChEMBL [52] database. The dataset was created using the ChEMBL web client [58] to download all ligands with EGFR binding affinity measurements (target ChEMBL ID CHEMBL203, measurement type IC_50_, relation =, assay type *B*). The compound set was filtered for assays with the target organism *Homo sapiens* and assay measurements in nanomolar units. Finally, IC_50_ values were converted to pIC_50_ values. After processing and filtering, the dataset contains 7, 327 data points.

#### Enamine Assembled (This work)

The Enamine Assembled dataset consists of 50M molecules assembled from the Enamine Building Block dataset, stored as *{* (bb_1_, bb_2_) *}*, product triples. The dataset was created by randomly sampling pairs of building blocks from the Enamine Building Block dataset and assembling the pairs with the set of 12 SMARTS reactions (also from the Enamine Building Block dataset). For each sampled pair, in silico assembly is attempted with all 12 SMARTS reactions. In the case where multiple products are generated (either from one SMARTS reaction yielding multiple valid products, or the sampled pair matching multiple SMARTS reaction templates), three products from the total product set are randomly selected, and the rest are discarded. This process is executed in parallel on 64 CPUs until the target dataset reaches 50M product molecules.

### 4.3 Scoring Functions

We use several scoring functions to comprehensively evaluate the performance of VVS across a diverse set of optimization tasks. We selected score functions representing computational chemistry, machine learning, and deep learning to represent the range of score functions commonly used in virtual screening.

#### 2ZDT Protein-Ligand Docking

Protein-ligand docking is selected as a score function to represent structure-based methods in virtual screening. We implement docking against the 2ZDT crystal structure of JNK3 using the OpenEye [59] toolkit with Python scripts and protein crystal structure from Klarich et al. [8]. To make docking compatible with a hill-climbing maximization framework, we flip the sign of docking scores so that higher values denote a better score. For each molecule evaluated, we dock 10 poses and select the best-scoring result as the final score.

#### 3D Shape and Electrostatic Similarity

Rapid Overlay of Chemical Structures (ROCS) [60] is selected as a score function to represent ligand-based methods in virtual screening. We implement ROCS using the OpenEye [59] toolkit with Python scripts from Klarich et al. [8]. The target ligand pose is also sourced from Klarich et al. [8]. For each molecule evaluated, we evaluated 10 poses and selected the best-scoring result as the final score.

#### Antibacterial Random Forest

A random forest classifier is selected as a score function to represent classical machine learning methods. Specifically, we use a trained random forest classifier from Swanson et al. [44] trained to predict antibiotic activity from 2D fingerprints.

#### EGFR MLP

A dense 13M parameter multilayer perceptron model is selected as a score function to represent modern deep learning scoring functions used in virtual screening. The model is trained on the Chembl ErbB1 dataset to predict molecule binding affinity against the protein EGFR, using RoBERTa-ZINC embeddings as input features. The model architecture, training, and evaluation details for this model are found in Supp. Notes 4.4.

### 4.4 EGFR pIC_50_ MLP Training

We train a 13M-parameter multilayer perceptron model to predict molecule binding affinity against the protein EGFR. This model is used as a scoring function for VVS benchmarks.

#### Dataset and Featurization

We use the Chembl ErbB1 dataset (Supp. Notes 4.2) and convert molecules using embeddings from the RoBERTa-ZINC model [41], mapping each molecule to a 768-dim embedding vector. We randomly split the dataset into 80% training and 20% validation. Target pIC_50_ values are normalized to zero mean and unit variance for training. During inference, the mean and variance of the training data are used to denormalize the model predictions to return standard pIC_50_ values.

#### Architecture

The model consists of four blocks, each containing a FFN_SwiGLU_ layer [54] with skip connection followed by LayerNorm [61] and dropout [62]. We use a hidden dimension of 1024, LayerNorm eps of 1e *−* 12, and dropout of 0.1. A linear head outputs a single prediction of the normalized pIC_50_ values. The model has approximately 13M parameters.

#### Training and Evaluation

We use the mean squared error loss, Adam optimizer [63], and a batch size of 32. We train for 30 epochs with 10% warm-up, maximum learning rate of 1*e −* 3, and cosine decay. The final model achieves a validation MSE loss of 0.367 and validation *R*^2^ metric of 0.636.

### 4.5 Embedding Compression Model Training and Evaluation

We train several Embedding Compression models used to compress CLM embeddings to a smaller dimension while maintaining relative embedding-to-embedding similarity relationships as defined by the native embeddings.

#### Dataset and Featurization

We create a training dataset of 30M molecules by randomly sampling 10M molecules from the Zinc 10M, Lyu 140M, and Enamine Assembled datasets. We convert molecules using embeddings from the RoBERTa-ZINC model [41], mapping each molecule to a 768-dim embedding vector.

#### Model Architecture

The Embedding Compression model consists of several encoder layers, which map the input embedding to the compressed size, followed by several decoder layers that reconstruct the compressed embedding back to the input size. The encoder and decoder both use 4 blocks consisting of a FFN_SwiGLU_ layer with skip connection followed by LayerNorm and dropout (Supp. Fig. 1**a**). Each encoder block preserves a hidden size of 768 (the native embedding size) before the final layer maps directly from 768 to the compressed size. The first decoder layer maps the compressed size back to the hidden size of 768, and preserves that size through the decoder layers. Supp. Table 1 shows the parameter count for the different compressor models.

#### Loss

The Embedding Compression model loss consists of two terms: cosine similarity loss on the decoder output and ChemRank loss with top-*K* weighting (Supp. Fig. 1**b**). We first compute the cosine similarity loss of the input native embeddings to the reconstructed embeddings from the decoder. Next, we compute the ChemRank loss with top *K ∈ {* 10, 100, 256*}* weighting between the native input embeddings and the compressed embeddings from the encoder. For both the native and compressed embeddings, we compute in-batch pairwise cosine similarity matrices *S*^target^ and *S*^pred^. We mask the diagonal of each similarity matrix to remove the self-similarity values (all equal to 1 by definition) and compute the ChemRank loss on the remaining values.

#### Training

We trained five independent compression models for compression sizes *d ∈{* 32, 64, 128, 256, 512 *}*. Each model is trained for one epoch with a batch size of 3072. We use the Adam optimizer with a maximum learning rate of 1e *− e*, 10% learning rate warmup, and cosine learning rate decay.

#### Validation Metrics

Given the downstream use case of nearest-neighbor retrieval, we designed an evaluation process to measure the performance of the compressed embeddings at this task. We embed a validation dataset of 1.5M molecules with the native embedding and all compression sizes (*d ∈ {* 32, 64, 128, 256, 512, 768 *}*). We use 1M molecules as a retrieval corpus, and the remaining 0.5M as retrieval queries. For each query, we first compute the *k* = 100 nearest neighbors using the native embedding. We then repeat the process for each compressed embedding size and compute the retrieval precision via the intersection of the native-embedding nearest neighbor list and the compressed-embedding nearest neighbor.

#### Loss Ablations and Analysis

We compare training the embedding compression model with different losses on the compressed embeddings, using ChemRank loss versus MSE loss (both computed on the pairwise similarity matrices, given the unequal sizes of the input and compressed embeddings), with and without top-*k* weighting. We find ChemRank loss with top-*k* weighting gives the best performance (Supp. Fig. 2a). We compare using a 1-layer model to a 4-layer model and find a modest but nonzero improvement from increasing model depth (Supp. Fig. 2b).

We find that adding top-*k* weighting to the ChemRank loss provides an average 7% improvement in precision@100 across all compression sizes, increasing model size from one to four layers provides an average 2.8% improvement over all compression sizes, and ChemRank with top-*k* weighting provides an average 5% improvement across all compression sizes compared to MSE loss with top-*k* weighting. We use the 4-layer model trained with ChemRank loss with top-*k* weighting for downstream applications.

#### Retrieval Evaluation

To evaluate retrieval performance, we embedded 10M molecules from the Zinc 10M dataset at all compression sizes. For each size, we built an HNSW index using Usearch [55]. We evaluated each index by index disk size, index build time, and average query latency for a query of 1024 embeddings with *k*=100. We find that index disk size and index build time scale linearly with embedding size, query latency is flat from size 32 to size 64 and scales linearly from size 64 to size 768, and query latency for 768-dim embeddings is roughly 5.9x slower compared to 64-dim embeddings. Results are summarized in Supp. Fig. 3. For downstream applications, we use 256-dim compressed embeddings for building block spaces and 128-dim compressed embeddings for enumerated spaces. Additionally, Supp. Fig. 14 and 15 provide a qualitative evaluation of comparative retrieval performance for different levels of compression.

### 4.6 Embedding Decomposer Training Details

#### Dataset and Featurization

The Embedding Decomposer model is trained on the Enamine Assembled dataset, which is composed of *{* (bb_1_, bb_2_), product*}* triples. The objective of the Embedding Decomposer model is to use a product molecule embedding to predict two building block embeddings. The dataset is preprocessed by canonicalizing all smiles and sorting building blocks by SMILES string length such that (len(bb1) *≤* len(bb2)). We find sorting building blocks by length provides an average 25% boost in performance across embedding sizes (Fig. 2d).

Each *{* (bb_1_, bb_2_), product*}* triple is embedded with the RoBERTa-ZINC model to 768-dim embeddings. The set of building blocks used to create the Enamine Assembled dataset (130k unique building blocks from the Enamine Building Block dataset) and embedded at all sizes (*d ∈ {* 32, 64, 128, 256, 512, 768*}*) and stored as lookup tables for the loss computation.

#### Model Architecture

The Embedding Decomposer model consists of three stages: input projection, trunk, and output projection (Fig. 2a). The input projection consists of several heads that map the input embedding from the input size (*d ∈ {* 32, 64, 128, 256, 512, 768 *}*) to a common hidden size of 1024. Each input projection head is a single FFN_SwiGLU_ layer. The trunk consists of 9 FFN_SwiGLU_ layers. The output projection consists of several linear layers, one for each output dimension (*d* = 32, 64, 128, 256, 512, 768).

Each output projection head maps the 1024 dim hidden vector to two embeddings of the corresponding output dimension. The model has 43.6M trainable parameters.

#### Multi-Size Batching

We train the Embedding Decomposer model to decompose any input embedding size (*d ∈ {* 32, 64, 128, 256, 512, 768 *}*) to any output size embedding. All size combinations are trained simultaneously (Supp. Fig. 2c). At training time, the model receives a batch of native product molecule embeddings of size (*B*, 768). We use the frozen Embedding Compression model encoders to compress the batch to all compression sizes. The Embedding Decomposer model then receives six separate input batches, one for each compression size. The input projection heads are used to map all input batches to the common hidden size of 1024. The batches are then stacked together to form an aggregate batch of size (6, *B*, 1024). The aggregate batch is processed by the shared trunk. The aggregate batch is then processed by six different output projection heads, each outputting a different embedding size. The final output is six output batches of size (6, *B*, 2, *D*) for a total of 36 output predictions per batch, representing the mapping of all input sizes to all output sizes.

#### Loss

For each output batch, we look up the ground-truth building block embeddings of the appropriate dimension from the pre-computed building block embedding tables. We compute the cosine similarity between the predicted embeddings and the ground-truth embeddings. Next, we randomly sample a batch of reference building block embeddings from the pre-computed embedding table. We compute the pairwise similarities of the predicted and ground-truth embeddings against the reference embeddings, yielding two similarity matrices. We compute the ChemRank loss on these similarity matrices (Fig. 2b). The loss is computed individually for each input size to output size combination. The cosine similarity and ChemRank loss terms are then summed with equal weight.

#### Training

We train for one epoch with a batch size of 2,048 and 3,072 reference embeddings for the loss computation. We use the Adam optimizer with a maximum learning rate of 1*e −* 3, 10% warm-up, and cosine learning rate decay. During training, we have 43.6M active parameters in the model, plus 29M frozen parameters in the Embedding Compression models and 140M frozen parameters for the precomputed building block embeddings.

#### Evaluation

We evaluate the model with a held-out set of 11M product molecules. For each item in the validation set, we compute predicted building block embeddings for all input to output size combinations. We use these embeddings to retrieve the *k* = 10 nearest neighbors building blocks from the Enamine Building Block dataset. We measure retrieval precision against the ground truth building block embeddings.

#### Ablations and Analysis

We compare average precision@10 performance for all input to output size combinations for different losses - cosine similarity, MSE loss, ChemRank with top-*k* weighting, and combinations therein (Fig. 2d,e). Cosine similarity and MSE loss directly compare predicted building block embeddings to the ground truth embeddings, while the ChemRank variants compare using the similarity matrix against the reference embeddings. We find that using cosine similarity with ChemRank performs best. We evaluate the impact of sorting building blocks by SMILES string length prior to training and find that sorting by length gives an average 25% performance boost across all embedding sizes.

We evaluate retrieval-precision-at-*k* and retrieval-accuracy-at-*k* (defined as the percent of predictions that recovered both target building block molecules within *k* results) for all input size to output size combinations (Supp. Fig. 4). We find that input size is the main driver of retrieval performance, while output size has minimal impact. Overall, we find input embedding sizes of 128 and above have roughly the same performance, while input embedding sizes 64 and below show a sharp performance drop.

We investigate retrieval failures and find two dominant failure modes: (i) “flipped predictions”, where both building blocks are recovered, but in the opposite order as expected by the dataset design, and (ii) complete failures, where the target building block is not retrieved by either predicted embedding within *k* results. We find that “flipped predictions” occur most often when the target building block SMILES strings are almost the same length, with a flip rate of 21% when the target SMILES strings are exactly the same length, and dropping to near zero when the target SMILES string lengths differ by 5 characters or more. We find that missed predictions are more common for short SMILES strings, suggesting the model struggles to differentiate building blocks with less than 10 atoms (Supp. Fig. 5b).

### 4.7 Cosine Similarity and Tanimoto Similarity Comparison

Cosine similarity computed on dense CLM embeddings and Tanimoto similarity computed on sparse molecular fingerprints are different computational approaches to evaluating molecular similarity. To assess the similarities and differences between these two metrics as a means of evaluating chemical similarity, we randomly sample 2.3M unique molecule pairs from the Enamine Assembled, Lyu 140M, and COCONUT 2.0 datasets. For each pair, we evaluate cosine similarity on 256-dim CLM embeddings and Tanimoto similarity using ECFP-4 fingerprints. We find a moderate correlation between the metrics, with Pearson and Spearman correlations of 0.5268 and 0.5265 respectively. We find that the correlation increases for highly similar molecule pairs (Supp. Fig. 7c,d). We also note that cosine similarity values tend to be consistently higher than Tanimoto similarity values (Supp. Fig. 7b). When plotting these two similarity metrics against each other, we observe a distinct “boomerang” shape (Supp. Fig. 7a).

Qualitative analysis of discordant pairs revealed fundamental differences in what these metrics capture. Cases with high cosine and low Tanimoto similarity (Supp. Fig. 21) display high-level molecule similarity with substantial atom-level differences, indicating that cosine similarity favors global molecular assessment, while Tanimoto similarity is more sensitive to atom-level features. Conversely, cases with low cosine and high Tanimoto similarity (Supp. Fig. 22) display molecules with repeated substructures. This discrepancy is likely caused by chemical fingerprints—generated by feature hashing—not accounting for feature frequency, effectively making Tanimoto similarity blind to this facet of molecular structures. Cosine similarity, derived from contextual embeddings, is more sensitive to repeat substructure patterns and their contribution to the overall molecular identity.

### 4.8 VVS Hyperparameter Evaluation

#### Learning Rate Sweeps

We assess the impact of learning rate magnitude on the gradient-step results with the following procedure:

1. Randomly sample 256 query molecules from the Enamine Assembled dataset
2. Embed each molecule with the RoBERTa-ZINC model and compress to 256-dim with the Embedding Compression model
3. Run BBKNN on each query embedding Enamine Building Block dataset with *k*_bb_ = 10
4. Score the result molecules with the EGFR MLP model
5. Use the result molecule embeddings and scores to estimate the gradient of the query embedding via Supp. Algorithm 2
6. For each (query, gradient) pair, take 100 steps along the gradient with learning rates ranging from 500 to 50, 000 in increments of 500
7. For each gradient-step embedding, run BBKNN to retrieve a set of result molecules and corresponding embeddings
8. Evaluate the cosine similarity of the gradient-step results against the gradient-step query embedding and the original query embedding

We find that result similarity to the original query drops modestly with learning rate, decreasing from 0.75 *→* 0.5 from learning rates 0 *→* 50, 000, while result similarity to the gradient-step embedding drops significantly, from 0.75*− >* 0.12 on the same learning rate interval (Supp. Fig. 8a). The low similarity between results and the gradient-step query that generated the results suggests that high learning rates result in gradient-step queries that are “off-manifold”, yielding results that are highly dissimilar from the gradient-step query. This highlights an advantage of BBKNN and VVS-grounding BBKNN results in retrieval from an explicit set of building blocks always results in valid, “on-manifold” results. This allows VVS to use large learning rates to rapidly explore disjoint regions of chemical space. Using a range of learning rates in the multi-scale query allows for both local area exploitation and distant chemical space exploration.

#### Hyperparameter evaluation

To investigate the impact of different hyperparameters, we ran 1,400 VVS searches on the Enamine Building Block dataset using the EGFR MLP score function with an inference budget of 50,000. The complete set of parameters tested can be found in Supp. Table 2. For each search, we evaluate the average score of the top *k ∈ {* 1, 5, 10, 100, 1000*}* results.

Parameters that strongly influence result scores are summarized in Supp. Fig. 9. With our hill climbing algorithm, taking multiple steps along the gradient outperforms single-step updates. (Supp. Fig. 9b). We find improvements increasing the maximum learning rate from 1,000 to 5,000, but little benefit to increasing from 5,000 to 10,000 (Supp. Fig. 9d). We also find re-sampling *p*_exploit_ high-scoring results from previous iterations improves results over not re-sampling (Supp. Fig. 9c). We show that a greedy “top-1” approach — choosing the top scoring result in each VVS iteration to be the new query embedding — outperforms the “standard” hill climbing approach of computing the gradient at the original query embedding (Supp. Fig. 9a).

While tuning the number of iterations, batch size, and *k*_*bb*_ values do not strongly impact the final scores, they do impact overall runtime. Larger values for these parameters result in more results scored per iteration, causing the overall process to reach the inference limit faster.

#### Result Diversity

We evaluated the total molecule result set from the 1,400 VVS hyperparameter sweeps using the EGFR scoring function to determine the diversity of results. Of 50M unique result molecules, 82.7% of results appear once, 10.5% twice, 3.1% of results appear three times, 3.1% of results appear between 4–10 times, and 0.44% of results appear more than 10 times (Supp. Fig. 10).

### 4.9 VVS Benchmark Details

#### Method Implementation

Implementation details for different benchmark search methods.

**SyntheMol**: We use the author’s GitHub implementation as-is with a custom score function.

**RxnFlow**: We use the author’s GitHub implementation as-is with a custom score function.

**RAD**: We use the author’s GitHub implementation as-is with a custom score function. In testing the author’s codebase, we found the method ran substantially faster when using 8 CPUs concurrently instead of the maximum available 64 CPUs. Consequently, we run all RAD benchmarks with 8 CPUs in a good-faith effort to showcase the best performance of the method.

##### Thompson Sampling

The author’s implementation uses a single SMARTS reaction and requires a minimum of one inference scoring per building block as part of a warm-up step, which would exceed the 50*k* inference budget. We modify the author’s method to use multiple SMARTS reactions and use a partial warm-up. To implement the partial warm-up, we use a hyperparameter *p*_warmup_ to allocate a percentage of the total inference budget to the warm-up process. Any molecules not seen during the warm-up have their prior distribution initialized with the average of all molecules seen during the warm-up phase. We test different values of *p*_warmup_ in the hyperparameter sweep to determine what value yields the best performance.

###### VVS

We implemented VVS as a single Python program, using PyTorch for models and embedding operations, RDKit for reaction assembly, and basic Python multiprocessing for parallelization. We use 256-dim embeddings for the Enamine Building Block building block dataset and 128-dim embeddings for the the much larger (*∼* 10^8^ embeddings) enumerated Lyu 140M dataset, both stored has a HNSW index using Usearch [55].

###### Reaction Implementation

All methods were given the same SMARTS reaction set and limited to a single reaction per product so that all product molecules are the result of assembling exactly two building blocks.

###### Compute

All methods were provided with the same compute resources - 1 A100 GPU and 64 CPUs - however, actual compute usage depended on the author’s code implementation. VVS and RxnFlow both used 1 GPU and 64 CPUs, while SyntheMol and Thompson Sampling each used a single CPU (see Supp. Table 4).

#### Scoring Back-End

All four score functions were implemented as consumers of a RabbitMQ queue. Benchmark engines submit SMILES strings to the score consumers via RPC, guaranteeing identical code paths and equal CPU/GPU resources for each method. All scoring clients are implemented with caching so that redundant scoring does not contribute to the total inference limit. The final benchmarks are run individually to avoid runtime impacts from different processes competing for the same scoring resources.

#### Hyperparameter Sweeps and Final Evaluation

For all benchmarked methods, we evaluated a range of hyperparameters using the EGFR MLP score function. Each test is run in triplicate and is evaluated on the average top-*k* = 1, 10, 100 scoring results in each run. The best results are summarized in Supp. Table 3.

The best hyperparameters from each sweep were run in 5 replicates across all score functions. The average results for searching the Enamine Building Block building block dataset are presented in Fig. 5, Supp. Table 5, and results for searching the Lyu 140M enumerated dataset are presented in Supp. Fig. 11, Supp. Table 6.

### 4.10 Extension of BBKNN to Multi-Step Assembly

Here we propose a theoretical framework for extending the Embedding Decomposer and BBKNN algorithm to multi-step reactions and molecules composed of three or more building blocks. The two-block version of the Algorithm 1 can be lifted to an arbitrary but bounded number of building blocks *𝒯* by (i) expanding the Embedding Decomposer model to map the query embedding **e**_*q*_ to predict *𝒯* total embeddings via (**e**_1_, **e**_2_, …, **e** _*𝒯*_) = *𝒟* (**e**_*q*_) and (ii) adding an auxiliary softmax classification head that predicts *𝒯* ^*\**^, the number of building blocks in the query. At inference time, only the first *𝒯* ^*\**^ *≤ 𝒯* predicted embeddings are used for downstream retrieval and assembly. Each predicted embedding 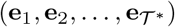 is used for a separate k-nearest-neighbors search to yield *𝒯* ^*\**^ pools of candidate building blocks via *N*_*t*_ = ???(**e**_*t*_, *ℬ, k*_bb_).

Assembly proceeds as a left-deep reaction chain. Let *ℛ*^(*s*)^ denote the set of SMARTS assembly reactions for stage *s ∈ {*1, …, *𝒯* ^*\**^ *−* 1*}*. Stage *s* = 1 enumerates every triple (*b*_*a*_, *b*_*b*_, *r*_*j*_) with *b*_*a*_ *∈ N*_1_, *b*_*b*_ *∈ N*_2_, *r*_*j*_ *∈ ℛ*^(1)^ and keeps the valid products *𝒫* ^(1)^. Stage *s* = 2 treats each product *p ∈ 𝒫* ^(1)^ as the “left” reactant and pairs it with every block *b ∈ N*_3_ under every reaction *r ∈ ℛ*^(2)^, producing a set of products *𝒫* ^(2)^. This process repeats until stage *𝒯* ^*\**^. Formally,

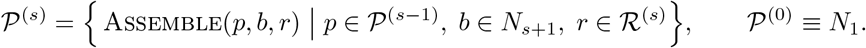

The final products 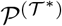 are embedded and evaluated by their cosine similarity to the query embedding **e**_*q*_. It may be beneficial to return all products 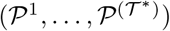; this determination is left to future research.

This framework preserves the key invariants of the Embedding Decomposer model and the BBKNN algorithm, while generalizing to *𝒯*-component synthesis. The classification head on the Embedding Decomposer explicitly predicts how many building blocks are needed for each query, keeping assembly efficient for simpler (1- or 2-block) targets, while also scaling up to the architectural limit *𝒯*.

## References

[1] Hilten, N., Chevillard, F., Kolb, P.: Virtual compound libraries in computerassisted drug discovery. J. Chem. Inf. Model. 59(2), 644–651 (2019)

[2] Walters, W.P.: Virtual chemical libraries. J. Med. Chem. 62(3), 1116–1124 (2019)

[3] Sadybekov, A.V., Katritch, V.: Computational approaches streamlining drug discovery. Nature 616(7958), 673–685 (2023)

[4] Warr, W.A., Nicklaus, M.C., Nicolaou, C.A., Rarey, M.: Exploration of ultralarge compound collections for drug discovery. J. Chem. Inf. Model. 62(9), 2021–2034 (2022)

[5] Enamine Ltd.: REAL Database: REAL Compounds. https://enamine.net/compound-collections/real-compounds/real-database Accessed 2025-06-11

[6] WuXi AppTec: GalaXi Molecules: Virtual Screening Services. https://wuxibiology.com/drug-discovery-services/hit-finding-and-screening-services/virtual-screening/ Accessed 2025-06-11

[7] Korn, M., Ehrt, C., Ruggiu, F., Gastreich, M., Rarey, M.: Navigating large chemical spaces in early-phase drug discovery. Current Opinion in Structural Biology 80, 102578 (2023) 10.1016/j.sbi.2023.102578

[8] Klarich, K., Goldman, B., Kramer, T., Riley, P., Walters, W.P.: Thompson samplingan efficient method for searching ultralarge synthesis on demand databases. Journal of Chemical Information and Modeling 64(4), 1158–1171 (2024)

[9] Kuan, J., Radaeva, M., Avenido, A., Cherkasov, A., Gentile, F.: Keeping pace with the explosive growth of chemical libraries with structure-based virtual screening. Wiley Interdiscip. Rev. Comput. Mol. Sci. 13(6) (2023)

[10] Bellmann, L., Penner, P., Rarey, M.: Topological similarity search in large combinatorial fragment spaces. Journal of Chemical Information and Modeling 61(1), 238–251 (2020)

[11] Schmidt, R., Klein, R., Rarey, M.: Maximum common substructure searching in combinatorial make-on-demand compound spaces. Journal of Chemical Information and Modeling 62(9), 2133–2150 (2021) 10.1021/acs.jcim.1c00640

[12] Liphardt, T., Sander, T.: Fast substructure search in combinatorial library spaces. J. Chem. Inf. Model. 63(16), 5133–5141 (2023)

[13] Rarey, M., Stahl, M.: Similarity searching in large combinatorial chemistry spaces. J. Comput. Aided Mol. Des. 15(6), 497–520 (2001)

[14] Sadybekov, A.A., Sadybekov, A.V., Liu, Y., Iliopoulos-Tsoutsouvas, C., Huang, X.-P., Pickett, J., Houser, B., Patel, N., Tran, N.K., Tong, F., Zvonok, N., Jain, M.K., Savych, O., Radchenko, D.S., Nikas, S.P., Petasis, N.A., Moroz, Y.S., Roth, B.L., Makriyannis, A., Katritch, V.: Synthon-based ligand discovery in virtual libraries of over 11 billion compounds. Nature 601(7893), 452–459 (2022)

[15] Zabolotna, Y., Volochnyuk, D.M., Ryabukhin, S.V., Gavrylenko, K., Horvath, D., Klimchuk, O., Oksiuta, O., Marcou, G., Varnek, A.: SynthI: A new open-source tool for synthon-based library design. J. Chem. Inf. Model. 62(9), 2151–2163 (2022)

[16] Software, N.: SmallWorld. https://www.nextmovesoftware.com/smallworld.html Accessed 2025-06-11

[17] Das, M., Ghosh, A., Sunoj, R.B.: Advances in machine learning with chemical language models in molecular property and reaction outcome predictions. J. Comput. Chem. 45(14), 1160–1176 (2024)

[18] Kirchoff, K.E., Wellnitz, J., Hochuli, J.E., Maxfield, T., Popov, K.I., Gomez, S., Tropsha, A.: Utilizing low-dimensional molecular embeddings for rapid chemical similarity search. In: European Conference on Information Retrieval, pp. 34–49 (2024). Springer

[19] Zhao, X., Wang, M., Zhao, X., Li, J., Zhou, S., Yin, D., Li, Q., Tang, J., Guo, R.: Embedding in Recommender Systems: A Survey (2023). https://arxiv.org/abs/2310.18608

[20] Long, L., Li, R., Zhang, J.: Artificial intelligence in retrosynthesis prediction and its applications in medicinal chemistry. J. Med. Chem. 68(3), 2333–2355 (2025)

[21] Jiang, Y., Yu, Y., Kong, M., Mei, Y., Yuan, L., Huang, Z., Kuang, K., Wang, Z., Yao, H., Zou, J., Coley, C.W., Wei, Y.: Artificial intelligence for retrosynthesis prediction. Engineering (Beijing) (2022)

[22] Yan, C., Ding, Q., Zhao, P., Zheng, S., Yang, J., Yu, Y., Huang, J.: Retroxpert: Decompose retrosynthesis prediction like a chemist. Advances in Neural Information Processing Systems 33, 11248–11258 (2020)

[23] Dai, H., Li, C., Coley, C., Dai, B., Song, L.: Retrosynthesis prediction with conditional graph logic network. Advances in Neural Information Processing Systems 32 (2019)

[24] Tavakoli, M., Baldi, P., Carlton, A.M., Chiu, Y.T., Shmakov, A., Van Vranken, D.: Ai for interpretable chemistry: Predicting radical mechanistic pathways via contrastive learning. Advances in Neural Information Processing Systems 36, 4080–4096 (2023)

[25] Sándor, M., Kiss, R., Keseru, G.M.: Virtual fragment docking by glide: a validation study on 190 protein-fragment complexes. J. Chem. Inf. Model. 50(6), 1165–1172 (2010)

[26] Oliveira, T.A., Silva, M.P., Maia, E.H.B., Silva, A.M., Taranto, A.G.: Virtual Screening Algorithms in Drug Discovery: A Review Focused on Machine and Deep Learning Methods. Drugs and Drug Candidates 2(2), 311–334 (2023) 10.3390/ddc2020017

[27] Ferla, M.P., Sánchez-García, R., Skyner, R.E., Gahbauer, S., Taylor, J.C., Delft, F., Marsden, B.D., Deane, C.M.: Fragmenstein: Predicting Protein-Ligand Structures of Compounds Derived from Known Crystallographic Fragment Hits Using a Strict Conserved-Binding–Based Methodology. ChemRxiv (2024). 10.26434/chemrxiv-2024-17w01

[28] Wang, M., Li, S., Wang, J., Zhang, O., Du, H., Jiang, D., Wu, Z., Deng, Y., Kang, Y., Pan, P., Li, D., Wang, X., Yao, X., Hou, T., Hsieh, C.-Y.: ClickGen: Directed exploration of synthesizable chemical space via modular reactions and reinforcement learning. Nat. Commun. 15(1), 10127 (2024)

[29] Powers, A.S., Yu, H.H., Suriana, P., Koodli, R.V., Lu, T., Paggi, J.M., Dror, R.O.: Geometric deep learning for structure-based ligand design. ACS Cent. Sci. 9(12), 2257–2267 (2023)

[30] Zhang, O., Huang, Y., Cheng, S., Yu, M., Zhang, X., Lin, H., Zeng, Y., Wang, M., Wu, Z., Zhao, H., Zhang, Z., Hua, C., Kang, Y., Cui, S., Pan, P., Hsieh, C.-Y., Hou, T.: FragGen: towards 3D geometry reliable fragment-based molecular generation. Chem. Sci. 15(46), 19452–19465 (2024)

[31] Beroza, P., Crawford, J.J., Ganichkin, O., Gendelev, L., Harris, S.F., Klein, R., Miu, A., Steinbacher, S., Klingler, F.-M., Lemmen, C.: Chemical space docking enables large-scale structure-based virtual screening to discover rock1 kinase inhibitors. Nature Communications 13(1), 6447 (2022)

[32] Verdonk, M.L., Giangreco, I., Hall, R.J., Korb, O., Mortenson, P.N., Murray, C.W.: Docking performance of fragments and druglike compounds. J. Med. Chem. 54(15), 5422–5431 (2011)

[33] Cohn, D., Atlas, L., Ladner, R.: Improving generalization with active learning. Machine learning 15, 201–221 (1994)

[34] Graff, D.E., Shakhnovich, E.I., Coley, C.W.: Accelerating high-throughput virtual screening through molecular pool-based active learning. Chemical Science 12(22), 7866–7881 (2021) 10.1039/D0SC06805E

[35] Korablyov, M., Liu, C.-H., Jain, M., van der Sloot, A.M., Jolicoeur, E., Ruediger, E., Nica, A.C., Bengio, E., Lapchevskyi, K., St-Cyr, D., Schuetz, D.A., Butoi, V.I., Rector-Brooks, J., Blackburn, S., Feng, L., Nekoei, H., Gottipati, S., Vijayan, P., Gupta, P., Rampášek, L., Avancha, S., Bacon, P.-L., Hamilton, W.L., Paige, B., Misra, S., Jastrzebski, S.K., Kaul, B., Precup, D., Hernández-Lobato, J.M., Segler, M., Bronstein, M., Marinier, A., Tyers, M., Bengio, Y.: Generative Active Learning for the Search of Small-molecule Protein Binders. arXiv (2024)

[36] Kozyrev, V., Sindt, F., Rognan, D.: Active learning to select the most suitable reagents and one-step organic chemistry reactions for prioritizing target-specific hits from ultralarge chemical spaces. J. Chem. Inf. Model. 65(2), 693–704 (2025)

[37] Khalak, Y., Tresadern, G., Hahn, D.F., Groot, B.L., Gapsys, V.: Chemical space exploration with active learning and alchemical free energies. Journal of Chemical Theory and Computation 18(10), 6259–6270 (2022)

[38] Yang, Y., Yao, K., Repasky, M.P., Leswing, K., Abel, R., Shoichet, B.K., Jerome, S.V.: Efficient exploration of chemical space with docking and deep learning. J. Chem. Theory Comput. 17(11), 7106–7119 (2021)

[39] Smith, J.S., Nebgen, B., Lubbers, N., Isayev, O., Roitberg, A.E.: Less is more: Sampling chemical space with active learning. The Journal of chemical physics 148(24) (2018)

[40] Graff, D.E., Aldeghi, M., Morrone, J.A., Jordan, K.E., Pyzer-Knapp, E.O., Coley, C.W.: Self-focusing virtual screening with active design space pruning. J. Chem. Inf. Model. 62(16), 3854–3862 (2022)

[41] Heyer, K.: Roberta-zinc-480m (2023). https://huggingface.co/entropy/roberta_zinc_480m

[42] Lyu, J., Wang, S., Balius, T.E., Singh, I., Levit, A., Moroz, Y.S., O’Meara, M.J., Che, T., Algaa, E., Tolmachova, K., et al.: Ultra-large library docking for discovering new chemotypes. Nature 566(7743), 224–229 (2019)

[43] Chandrasekhar, V., Rajan, K., Kanakam, S.R.S., Sharma, N., Weißenborn, V., Schaub, J., Steinbeck, C.: Coconut 2.0: a comprehensive overhaul and curation of the collection of open natural products database. Nucleic Acids Research 53(D1), 634–643 (2024) 10.1093/nar/gkae1063 https://academic.oup.com/nar/article-pdf/53/D1/D634/60816582/gkae1063.pdf

[44] Swanson, K., Liu, G., Catacutan, D.B., Arnold, A., Zou, J., Stokes, J.M.: Generative ai for designing and validating easily synthesizable and structurally novel antibiotics. Nature Machine Intelligence 6(3), 338–353 (2024)

[45] Seo, S., Kim, M., Shen, T., Ester, M., Park, J., Ahn, S., Kim, W.Y.: Generative flows on synthetic pathway for drug design. arXiv preprint arXiv:2410.04542 (2024)

[46] Asano, Y., Kitamura, S., Ohra, T., Itoh, F., Kajino, M., Tamura, T., Kaneko, M., Ikeda, S., Igata, H., Kawamoto, T., et al.: Discovery, synthesis and biological evaluation of isoquinolones as novel and highly selective jnk inhibitors (2). Bioorganic & medicinal chemistry 16(8), 4699–4714 (2008)

[47] Hall, B., Keiser, M.: Retrieval Augmented Docking Using Hierarchical Navigable Small Worlds (2024). 10.26434/chemrxiv-2024-qsdd1

[48] Team, Q.: Qdrant: High-performance, massive-scale Vector Database (2020). https://github.com/qdrant/qdrant

[49] Malkov, Y.A., Yashunin, D.A.: Efficient and robust approximate nearest neighbor search using hierarchical navigable small world graphs. IEEE transactions on pattern analysis and machine intelligence 42(4), 824–836 (2018)

[50] Labs, D.: Dagster. https://dagster.io/ Accessed 2025-06-11

[51] Tingle, B.I., Tang, K.G., Castanon, M., Gutierrez, J.J., Khurelbaatar, M., Dandarchuluun, C., Moroz, Y.S., Irwin, J.J.: ZINC-22-A Free Multi-Billion-Scale Database of Tangible Compounds for Ligand Discovery. Journal of Chemical Information and Modeling 63(4), 1166–1176 (2023) 10.1021/acs.jcim.2c01253

[52] Gaulton, A., Hersey, A., Nowotka, M., Bento, A.P., Chambers, J., Mendez, D., Mutowo, P., Atkinson, F., Bellis, L.J., Cibrián-Uhalte, E., et al.: The chembl database in 2017. Nucleic acids research 45(D1), 945–954 (2017)

[53] Liu, Y.: Roberta: A robustly optimized bert pretraining approach. arXiv preprint arXiv:1907.11692 (2019)

[54] Shazeer, N.: Glu variants improve transformer. arXiv preprint arXiv:2002.05202 (2020)

[55] Vardanian, A.: USearch by Unum Cloud (2023). 10.5281/zenodo.7949416. https://github.com/unum-cloud/usearch

[56] RDKit: Open-source cheminformatics (2025). https://www.rdkit.org

[57] Ramsundar, B., Eastman, P., Walters, P., Pande, V., Leswing, K., Wu, Z.: Deep Learning for the Life Sciences. O’Reilly Media, ??? (2019). https://www.amazon.com/Deep-Learning-Life-Sciences-Microscopy/dp/1492039837

[58] Davies, M., Nowotka, M., Papadatos, G., Dedman, N., Gaulton, A., Atkinson, F., Bellis, L., Overington, J.P.: ChEMBL web services: streamlining access to drug discovery data and utilities. Nucleic Acids Research 43(W1), 612–620 (2015) 10.1093/nar/gkv352

[59] OpenEye, C.M.S.: ROCS, Santa Fe, NM (YEAR). http://www.eyesopen.com

[60] Rush, T.S. 3rd, Grant, J.A., Mosyak, L., Nicholls, A.: A shape-based 3-D scaffold hopping method and its application to a bacterial protein-protein interaction. J. Med. Chem. 48(5), 1489–1495 (2005)

[61] Ba, J.L., Kiros, J.R., Hinton, G.E.: Layer Normalization (2016). https://arxiv.org/abs/1607.06450

[62] Srivastava, N., Hinton, G., Krizhevsky, A., Sutskever, I., Salakhutdinov, R.: Dropout: A simple way to prevent neural networks from overfitting. Journal of Machine Learning Research 15(56), 1929–1958 (2014)

[63] Kingma, D.P., Ba, J.: Adam: A Method for Stochastic Optimization (2017). https://arxiv.org/abs/1412.6980

